# Integrated proteogenomics uncovers ancestry-specific and shared molecular drivers in localized prostate cancer

**DOI:** 10.1101/2025.11.28.691220

**Authors:** Cara C. Schafer, Tamara S. Abulez, Xijun Zhang, Kun-Lin Ho, Jiji Jiang, Denise Young, Jesse Fox, Kelly A. Conrads, Brian L. Hood, Gauthaman Sukumar, Darryl Nousome, Praveen-Kumar Raj-Kumar, Mariano Russo, Ayesha A. Shafi, Xiaofeng A. Su, Albert Dobi, Amina Ali, Sally Elsamanoudi, Jennifer Cullen, William D. Figg, Gyorgy Petrovics, Clifton L. Dalgard, Matthew D. Wilkerson, Nicholas W. Bateman, Thomas P. Conrads, Isabell A. Sesterhenn, APOLLO Research Network, Leigh Ellis, Craig D. Shriver, Gregory T. Chesnut, Shyh-Han Tan

## Abstract

Our integrative proteogenomics (genome, proteome and phosphoproteome) of localized prostate cancer (PCa) in an equal-access Military Health System patient cohort (57 Black and 55 White) revealed significant ancestry-associated differences. Somatic and germline regulatory differences converged on androgen, metabolic, PI3K/AKT/mTOR, and DNA damage response (DDR) pathways, with ancestry-specific immune- and stromal-associated signals. Black patients displayed greater genomic variability, enhanced androgen response, fatty-acid metabolism, and epithelial-mesenchymal transition, while White patients showed prevalent DDRG alterations, activated oncogenic signaling (MYC, E2F, mTORC1), and cell cycle regulation. Phosphoproteomics highlighted distinct kinase activities and candidate druggable dependencies. Multiomics integration revealed three exploratory tumor subtypes whose distinct biological programs were reproducibly validated. Ancestry-associated eQTLs supported inherited regulation of the proteome independent of CNAs. Ancestry-specific CNA and protein panels improved progression risk prediction beyond PSA and pathology models. These findings provide a framework for ancestry-informed prognostic models and generate testable hypotheses for precision therapies to reduce outcome disparities.

## INTRODUCTION

Prostate cancer (PCa) is the most common non-cutaneous cancer and the second leading cause of cancer-related death among men in the United States^1^. Black men experience disproportionately higher incidences and mortality rates^2–5^, a disparity that echoes global patterns in regions with large populations of African descent, such as the Caribbean and sub-Saharan Africa^6^. These consistent trends suggests underlying ancestry-related biological differences contribute to disease progression and outcomes. Although prior studies have pointed to inherited susceptibility alleles^7,8^, epigenetic modulation^9^, mitochondrial DNA variation^10–12^, driver gene defects^13–15^, and androgen receptor (AR) signaling differences as possible contributors^16–18^, a significant knowledge gap persists. Many large-scale proteogenomic studies^19–23^ have focused on patients of European ancestry, limiting generalizability and slowing progress towards equitable, ancestry-informed clinical strategies. Moreover, few studies have leveraged equal-access cohorts, where socioeconomic differences are minimized, and biological signals can be more confidently distinguished.

To addresses this gap, we conducted an integrative proteogenomics analysis of primary prostate tumors from a unique equal-access Military Healthcare System (MHS) cohort, comprising balanced representation of Black and White patients. This multi-omics approach, encompassing single-nucleotide variants (SNVs), somatic copy number alterations (CNAs), protein, and phosphoprotein profiles, together with well-annotated clinicopathologic and long-term follow-up data, provides an opportunity to uncover ancestry associated molecular pathways in clinically relevant setting. Rather than defining causal mechanism, our study is designed to generate testable hypothesis about the biological drivers of outcome disparities. Concurrently, the integrative design highlights potential biomarkers, therapeutic targets, and risk models that may ultimately inform precision medicine approaches for both Black and White men.

## RESULTS

To investigate how genetic ancestry affects PCa biology, the Applied Proteogenomic OrganizationaL Learning and Outcomes (APOLLO) Research Network conducted a proteogenomic analysis of localized PCas from a diverse cohort of 57 Black and 55 White patients treated at the Center for Prostate Disease Research (CPDR), Walter Reed National Military Medical Center (WRNMMC). Tumors underwent whole genome sequencing (WGS) and global proteome and phosphoproteome quantitative tandem-mass tag (TMT) mass spectrometry (MS). Patients were matched for tumor stage and grade (**Tables 1 & S1A**). Consistent with the trend over the past 30 years, Black men in the CPDR-APOLLO cohort were significantly younger at the time of surgery (p = 6.45E^-05^)^24^. A median follow-up time of over 10 years enabled meaningful analysis of clinical endpoints. The overall biochemical recurrence (BCR) rate of 26% and metastasis rate of 11% are consistent with other recorded cohorts^25,26^.

### Comparative analyses of prostate cancer genomes

We performed WGS on fresh-frozen specimens from 103 patients (52 Black and 51 White), using matched germline DNA from blood samples for accurate somatic mutation detection. Nearly all tumor and normal samples achieved their targeted coverage of 90X and 30X, respectively, with tumor purity averaging 41.0% (**Figure S1A-B**). Principal Component Analysis (PCA) of germline single-nucleotide polymorphism (SNP) markers, alongside HapMap and 1000 Genomes Project (1KGP) reference panels^27^, confirmed genetic ancestry largely correlated with self-identified race. As shown in **Figure S1C**, analysis of principal component 1 (PC1) quantified the primary ancestral mixture, revealing a broader diversity within the Black patient cohort, with varying levels of European admixture. While patient genetic ancestry largely matched self-identified race, we used the latter to describe patient groups for clinical contexts, and ancestry as a genomic variable to assess biological and molecular associations. This approach allows us to relate ancestry to biological mechanisms while maintaining consistency with existing terminology for health disparities. We profiled all samples for somatic SNVs, genomic rearrangements, and copy number variations (CNV). To investigate ancestry-associated etiologies, we evaluated genomes for *de novo* somatic mutation signatures. Of the ten signatures identified, seven were significantly associated with self-identified race: SNV2, SNV4, and IND2 were more frequent in Black patients; SNV3, IND1, IND3, and SV3, in White patients (p < 0.05; **Figure 1A**, Somatic signatures, p < 0.05; **Figure S1D, Tables S1B-C).**

**Figure 1.**
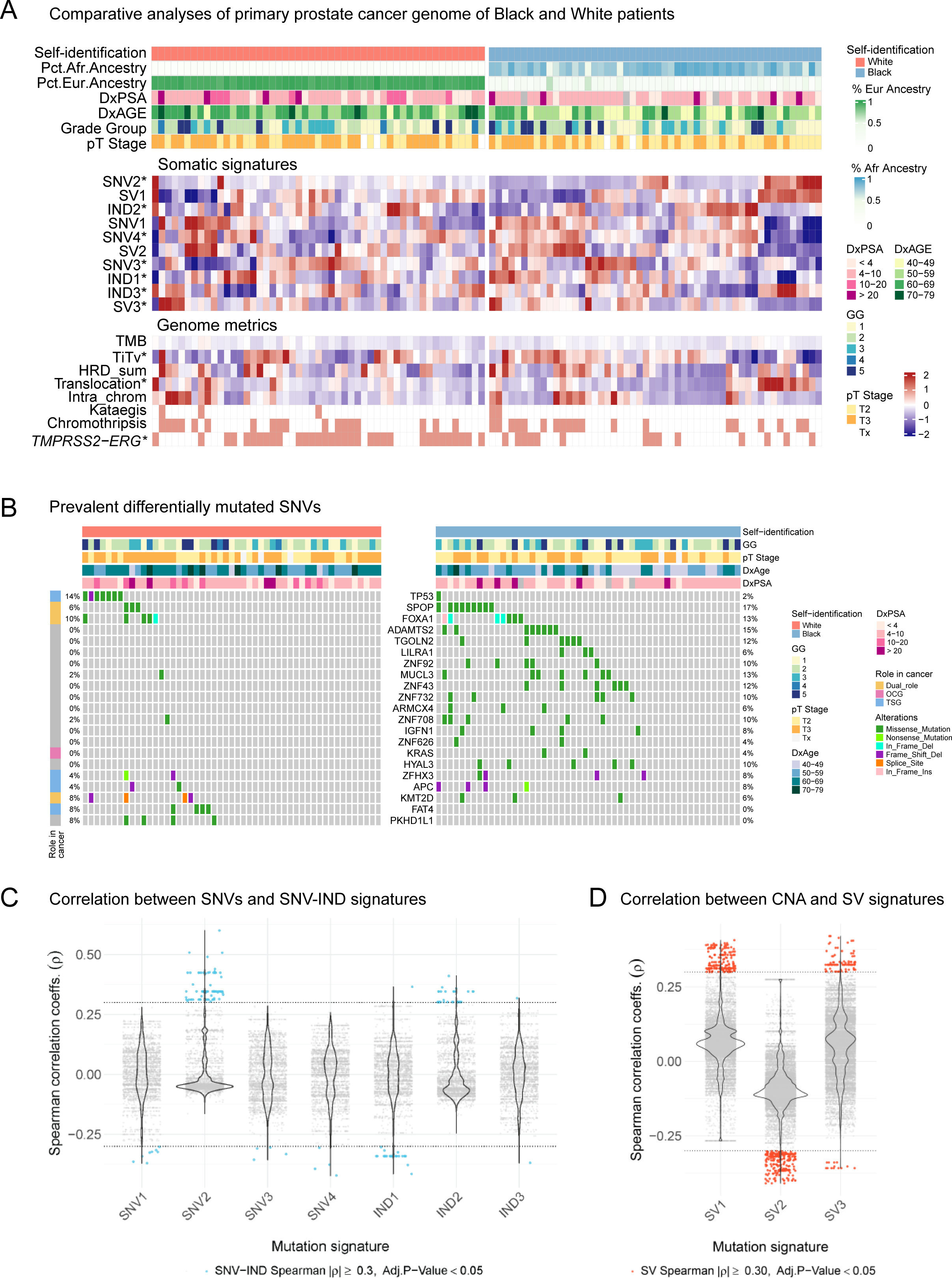
Differential genomic alterations in Black and White prostate cancer patients. (**A**) Differentially altered somatic mutational signatures and genomic metrics by self-identified race, *(p<0.05). (**B**) Recurrent SNVs, highlighting most prevalent and disproportionately represented mutations across groups (Fisher or Firth p < 0.05). Spearman correlation of SNV mutations vs. SNV and IND signatures (**C**), and sCNA vs. SV signatures (**D**; cut-off, ρ ≥ 0.3).

Our assessment of the global mutational properties revealed that while overall tumor mutation burden (TMB) was comparable, Black patients had significantly higher overall mutation counts (p = 0.0018), higher rates of translocation (p = 0.009) and transversion (p = 1.32E^-9^; **Figure 1A**, Genome metrics). We found no significant differences in homologous recombination deficiency score or intra-chromosomal structural variation. Overall chromosomal instability, including kataegis and chromothripsis events, was comparable. As expected, *TMPRSS2-ERG* fusions were more frequent in White patients (p < 0.001; **Figures S1E-F**).

The SNV ad IND signatures closely resembled those from the Pan-Cancer Analysis of Whole Genomes (PCAWG) Network^28–30^ (**Figures S1G-I**). Clock-like aging-related signatures, SNV3 and IND1, were more frequent in White patients. DNA mismatch repair deficiency (MMRd)-associated signatures were enriched in both groups: SNV4 in Black patients and IND3 in White patients. Of particular novelty, SNV2, characterized by a prominent TTA>TAA transversion peak, was enriched in Black patients, mirroring elevated transversion rates; additionally, SNV2 and IND2 signatures exhibited significantly greater genomic variability in Black patients (**Figures S1J-K**). Our evaluation of these mutational signatures showed a significant association between the IND3 signature and progression to BCR or metastasis, with this association remaining significant after adjustment for age, PSA levels, and percent African ancestry (**Figure S1L, Table S1D).**

### Ancestry-associated mutational landscapes

Across the entire cohort, we detected recurrent SNV alterations in known driver genes, including *FOXA1, SPOP*, *TP53*, and *APC* (**Figures 1B, S2A**, **Tables S1E-F**). In Black patients we detected higher mutation rates for *SPOP* (Firth p < 0.05; age- and PSA-adjusted) and *ADAMTS2, TGOLN2,* and *ZNF43* (Fisher and Firth p < 0.05), while *TP53* mutations were significantly more frequent in White patients (Fisher p < 0.05). Recurrent *FOXA1* mutations and frame-shift deletions in *APC* and *ZFHX3* were also observed in Black patients, though not significant. Notably, mutations were found in specific functional domains: TP53 within its DNA-binding domain, *SPOP* in the MATH domain^31^, and *FOXA1* in the forkhead DNA-binding α-helix 3^32^ and Wing2 domains^33^ (**Figures S2B-D**). Recurrent *ADAMTS2* and *HYAL3* variants resulting in identical amino-acid substitutions suggest potential novel hotspots (**Figures S2E-G**).

Spearman correlation revealed that the Black patient-enriched, transversion-associated SNV2 signature showed the strongest association with recurrent SNVs (ρ > 0.3, adj. p < 0.05), indicating that transversions are the dominant SNV class in this group (**Figure 1C, Table S1G**). SV1 and SV2 signatures had opposing associations with deletion-rich regions, while the amplification-associated SV3 signature was more prevalent in White patients (**Figure 1D, Tables S1H**).

### Ancestry-specific copy number alterations

At the focal level, recurrent deletions at 8p21 and 8q24 gains were common across both groups. White patients exhibited a higher burden of hemizygous deletions, with significantly more frequent deletions at 16q24.1, 17p13.1-2, 18q22.1, and 21q22.2 (Wilcoxon p < 0.05, **Figures 2A, S2H-I, Table S2A-C**). Using a combination of Fisher’s exact test and Elastic Net regression, we identified 1,631 genes with significant ancestry specific CNAs (408 in Black, 1,223 in White), spanning 48 loci in Black patients and 95 in White patients (**Figures S2J-K, Tables S2D-E)**. We confirmed recurrent amplifications (*MYC*, *NCOA2*), and deletions (*NRG1*, *NKX3-1*, *RB1*, and *CHD1*) in both ancestries. Notably, in Black patients Fisher’s exact test identified focal alterations, including 1q41 (*TGFB2*) and 3q13 (*GSK3B/LSAMP*), consistent with our prior finding of *LSAMP* deletions in African American PCa^14^. Elastic Net further revealed ancestry-associated differences in event types despite similar frequencies, including mixed gains or losses at 7q36.1(*KMT2C*) and 8q24.3 (*RECQL4*), versus predominantly gains in White patients. Additional recurrent alterations in Black patients included homozygous deletions at 2q22 (*LRP1B*), 3p11 (*EPHA3*), 5q12-14 (*RASA1, CENPK*), and 13q14 (*FOXO1*). Besides prevalent *PTEN* loss and *TMPRSS2-ERG* fusion, White patients showed a significantly higher rate of deletions at 16q23-24 (*ZFHX3*, *HSD17B2*, *FANCA*), and 17p12-13 (*MAP2K4*, *TP53, PIK3R5*) (Fisher p < 0.05). They also exhibited recurrent mixed alterations at 3p13 (*FOXP1*, *EIF4E3, SHQ1*)^34,35^, at 16p12.1-2 (*PALB2*, *PRKCB*, *IL21R*), and frequent homozygous deletions at 8p23 (*GATA4, NEIL2, ARHGEF10*) and 12p12.1 (*RECQL*, *KRAS*). These differentially altered genes prominently affected key pathways regulating oncogenic signaling, metabolism, cell proliferation, and inflammation. (**Figures 2B-C, Tables S2F**).

**Figure 2.**
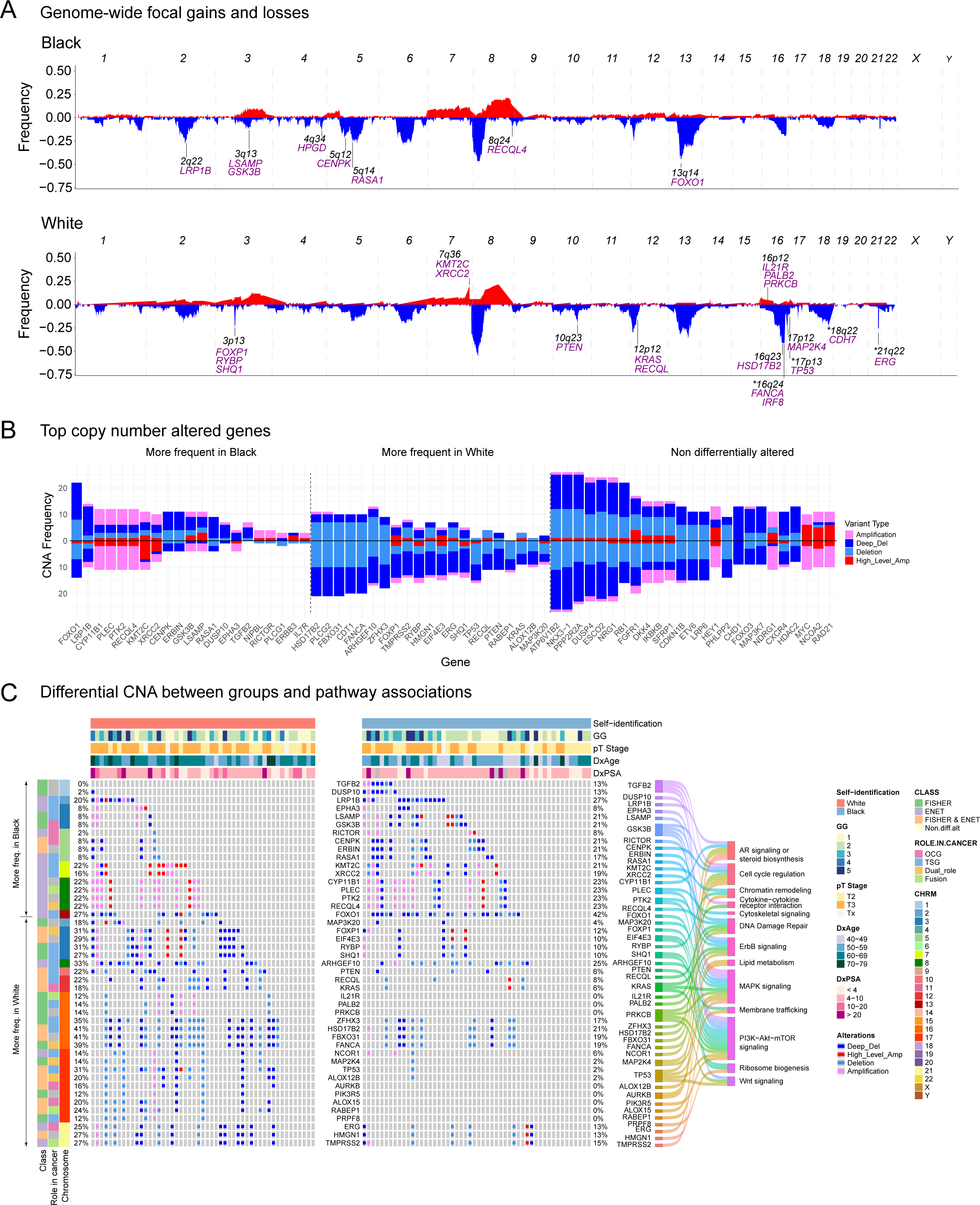
Gene level sCNA differences and pathway impact. (**A**) Genome-wide focal gains (red) and losses (blue) in Black and White patients. (**B**) Most frequent ancestry-enriched and commonly altered sCNA genes, ordered by prevalence (above midline, Black enriched; below, White enriched). (**C**) Differentially altered genes classified by method, linked to key cell signaling pathways.

### Ancestry-associated proteomic and phosphoproteomic differences

Our proteomics analysis of 101 fresh-frozen prostate tumors (48 Black, 53 White) identified 8,846 proteins, with 6,976 consistently quantified across all samples. Among these, 422 proteins were significantly altered (adj. p <0.05): 195 elevated in Black patients and 227 in White. Hierarchical clustering broadly separated samples by race. After adjusting for age and PSA, 236 proteins remained significant (103 in Black, 133 in White; adj. p <0.05) with strong concordance between adjusted and unadjusted results (71% overlap at nominal significance; 49% at FDR; **Figures S3A-D, Tables S3A-B)**. Most proteins were minimally affected by covariates, though several, including HPGD, COL2A1, and PLA2G2A, showed marked shifts in abundance or significance (**Figures S3E-F**).

Analysis of normal prostate epithelium (FFPE) from a subset of 25 patients (10 Black, 15 White) from this same cohort, quantified 5,219 proteins, with 578 significantly altered (379 elevated in White, 199 in Black, p <0.05). Tumor-normal comparisons confirmed ancestry-associated signatures that persisted after covariate adjustment, with GREB1, ANPEP, NQO2, and ADAMTS1 enriched in Black patients and REXO2, PLA2G7, TSN, and UDGH enriched in White patients (**Figures 3A**, **S3G-I, Tables S3C-D**).

**Figure 3.**
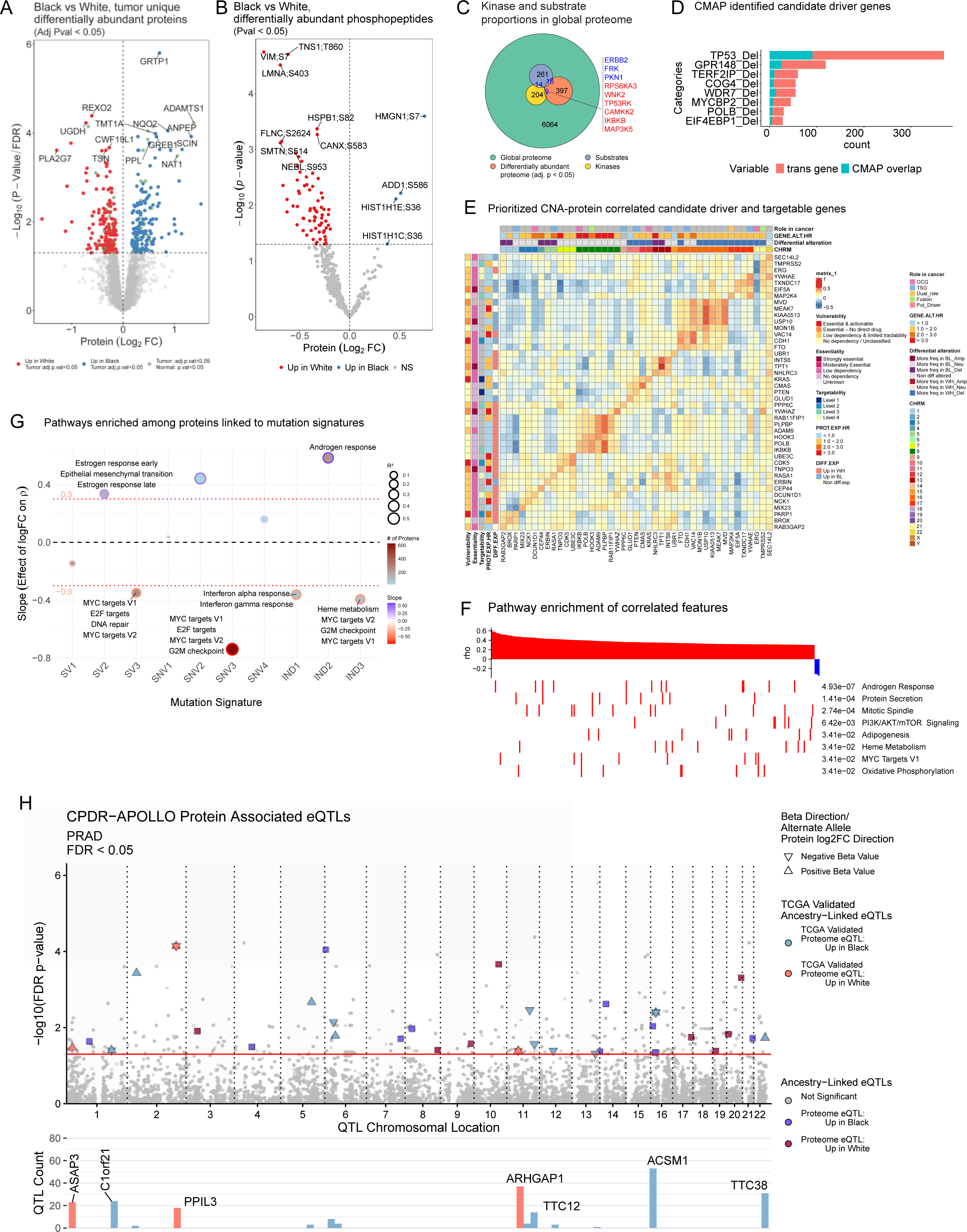
Proteomic and phosphoproteomic differences. **(A-B)** Volcano plots showing differential abundances of tumor-unique proteome (adj. p < 0.05), and phospho-proteome (p < 0.05) between Black and White tumors without age and PSA adjustment. **(C)** Kinase and phosphoprotein substrate proportions in the quantified proteome, including overlaps with differentially abundant proteins (adj. p < 0.05). **(D)** Candidate driver genes identified by LINCS/CMAP analysis (permutation FDR < 0.25) from CNA genes. (**E**) Heatmap of CNA-protein correlation (|ρ| > 0.3) for genes showing differential expression or alteration and strong prognostic or targetable features (|HR| ≥3 or annotated driver). **(F)** Hallmark pathways enriched among 271 *cis*-correlated features (|ρ| > 0.3). **(G)** Significantly enriched pathways of proteins associated with mutation-signatures (adj. p < 0.05; Slope: positive, Black-associated; negative, White-associated). **(H)** Top: PCa specific eQTLs mapping to proteins differentially abundant between Black and White patients, with allele frequency divergence between African and European populations. Squares indicate proteins elevated in Black (purple) or White (maroon) patients; triangles denote candidates with concordant transcript-level differences in TCGA (blue: elevated in Black; coral: elevated in White). Bottom: Number of validated eQTLs per gene; labeled genes have ≥10 associated eQTLs.

Phosphoproteome profiling quantified 6,456 phosphosites, of which 444 detected across all tumors and 79 were significantly altered (75 elevated in White patients, four in Black; p < 0.05; **Figure 3B**). These phosphosites mapped to 291 substrate proteins, 16 of which also differed at the global protein level, while nine of the 227 quantified kinases also showed differential abundance (adj. p < 0.05; **Figure 3C**). Notable phosphosites included TNS1;T860 and LMNA:S403 in White patients, and HMGN1;S7 and ADD1;S586 in Black patients. Phosphorylation differences remained robust after covariate adjustments, with 81 sites significant (**Figure S3J, Tables S3E-F)**.

### CNA–Protein concordance identifies known and novel candidate drivers

We used Spearman correlations to assess the impact of CNAs on protein abundance, distinguishing *cis*-effects (CNAs and protein changes at the same locus), from *trans*-effects (CNAs influence proteins encoded elsewhere) (**Figure S4A**). After filtering low variance features, 5,977 directly comparable CNA-protein pairs were retained. Of these 69.4% (4,146) showed positive correlation, with 271 (4.5%) being statistically significant (|ρ| > 0.3, p < 0.05; **Figures S4B-C, Table S4A**). Most significant *cis*-correlated pairs (172/271) were not differentially altered in copy number (CN) between ancestries and localized to contiguous hotspot on chromosomes 6, 8, 16, and 18^36,37^.

While *cis*-correlations directly link CNAs to protein abundance, many significant associations arose via *trans*-effects. Integration with LINCS knockdown profiles revealed *trans*-protein signatures for 26% of CNA genes^38,39^. CMAP analysis identified 407 candidate drivers supported by both gains and losses. Permutation-based testing (FDR < 0.249) confirmed known drivers such as *TP53* and nominated seven novel candidates (*GPR148*, *TERF2IP*, *COG4*, *WDR7*, *MYCBP2*, *POLB*, *EIF4EBP1*) as potential regulators of *trans*-effects (**Figure 3D**).

High confidence *cis*-correlated CNA-protein pairs (|ρ| ≥ 0.3) comprising candidate driver and targetable genes showed coordinated genomic-proteomic alterations with prognostic relevance across core oncogenic pathways, including RAS/MAPK, PI3K/AKT/mTOR, DNA damage response (DDR), ubiquitination, vesicular trafficking, EMT, and RNA processing (**Figure 3E, Tables S4B-C**). Integration with DepMap highlighted dependencies for EIF5A, USP10, TPT1, YWHAZ, and TNPO3, suggesting testable therapeutic vulnerabilities. Pathway enrichment further confirmed significant involvement of androgen signaling, PI3K/AKT/mTOR, cell cycle regulation, and MYC-driven proliferation (**Figure 3F, Tables S4D**). Protein abundance stratified by CNA status showed clear directional effects - amplifications increased and deletions decreased protein levels-with focal events at 8p11 (*IKBKB*, *HOOK3*) and 16q22 (*CDH1*) displaying pronounced gradients (**Figure S4D**).

### Mutational signatures and proteomic pathway activation

We next examined the association between mutational signatures and proteomic changes by using linear regression analysis of Spearman correlations (ρ) and protein log2 fold changes We considered slopes >0.3 and <−0.3 as significant (**Figure S4E, Table S4E**). Positive slopes (IND2, SNV2, SV2) were associated with proteins elevated in Black patients; negative slopes (IND1, IND3, SNV3, SV3), proteins elevated in White patients. Pathway enrichment analysis on signature-correlated proteins (ρ > 0.3) revealed that IND2 associated with androgen response and SV2 with estrogen responses and EMT in Black patients. In White patients, SV3 and SNV3 were associated with MYC and E2F targets, IND1 with interferon α/γ response, and IND3 with heme metabolism, MYC targets V1/V2 and G2M checkpoint pathways (**Figure 3G, Table S4F**). These findings demonstrate that ancestry-associated mutational processes affect distinct proteomic pathways.

### Ancestry-associated QTLs mapping to differentially abundant proteins

To assess germline contributions to proteomic differences beyond CNAs, we mapped differentially abundant proteins to PCa expression quantitative trait loci (eQTLs) from PanCanQTL, with pan-cancer corroboration **(Figures 3H, S5A, Table S5A-B**)^40^. Prioritized loci showed directionally concordant eQTL betas and protein changes by ancestry (MWU p<0.05), and ≥40% allele frequency differences across African and European populations in gnomAD, 1KGP, and CPDR-APOLLO germline data. Notable eQTL regulated proteins included ACSM1, TTC38, and C1ORF21 in Black patients, and ARHGAP1, ASAP3, and PPIL3 in White patients. Several candidates, (PPIL3, TTC38, ARHGAP1, ACSM1) were validated at the transcript level in TCGA-PRAD and in independent proteogenomic datasets generated from African and European cohorts^41^ (**Figures S5B-C, Table S5C)**. Among these, LGALS8 was associated with poorer prognosis, whereas FECH showed a protective effect, highlighting that germline regulatory variation can also shape the tumor proteome.

### Ancestry-associated CNA and protein pathway enrichment

Ancestry-associated CNA pathway enrichment revealed distinct profiles (**Figure 4A, Table S6A**), reflecting recurrent focal events such as 1q21, 8q23-24, and 12q13-q21 in Black patients, and 16q24 gain and 17p13 loss in White patients. Black patient profiles were enriched for immune-related programs, lineage-specific or oncogenic pathways, and epigenetic regulators (HDAC2, COBRA1, FOXP3 targets); White patient profiles showed enrichment in Polycomb- and SWI/SNF-associated clusters, chemokine-mediated signaling, and carbohydrate metabolism.

**Figure 4.**
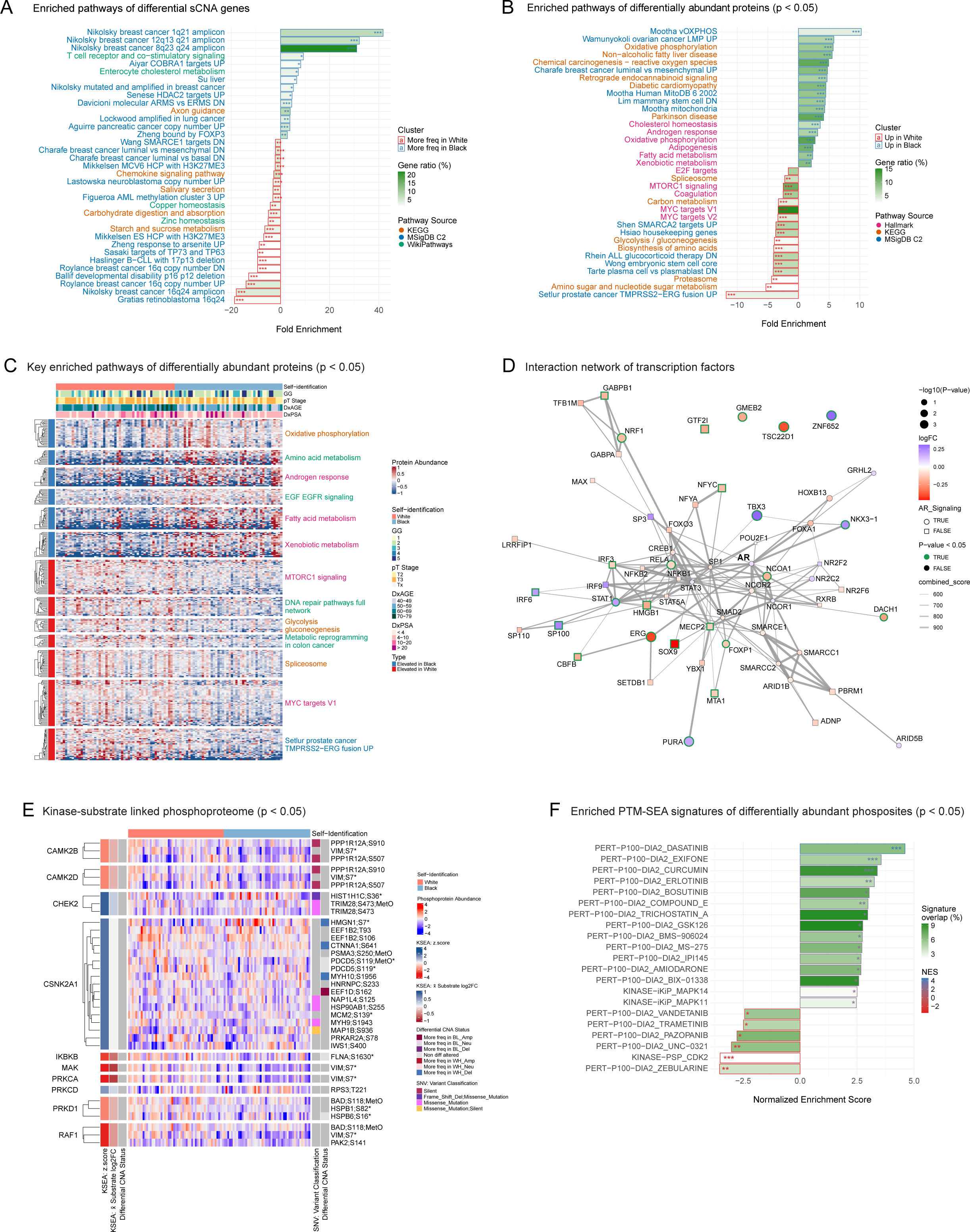
Differential pathway enrichments and interaction networks. **(A)** Top enriched pathways (adj. p < 0.05) from ancestry-associated CNA genes: bars indicate fold enrichment; fill intensity, gene ratio (%); bar outlines, race-associated enrichment; and label colors, database source. **(B)** As in **(A)**, top enriched pathways (adj. p < 0.05) for proteins differentially abundant by race (p < 0.05). **(C)** Heatmap of top enriched pathways from unadjusted proteome differences (p < 0.05). **(D)** AR-centered transcription factor network supported by STRING (score ≥ 700). Node shape represents AR pathway membership; fill color for log2FC (blue, elevated in Black; red, in White), green border for significance, edge width for interaction confidence. **(E)** Heatmap of significant kinase-substrate links for differentially abundant phosphoproteome (p < 0.05), annotated by KSEA z-score, mean substrate log_2_FC, differential CNA status of kinase and substrate, and SNV variant classification. **(F)** Top enriched pathways from PTM-SEA of phosphosites, showing ancestry-associated fold enrichment and percent signature overlap. Significance asterisks:**** (p<0.0001), ***(p<0.001), **(p<0.05), and *(p<0.1)

Proteomic pathway enrichment of differentially abundant proteins (**Figure 4B**, nominal p < 0.05) showed distinct but partially overlapping themes with the CNA results. Black patient tumors were enriched in mitochondrial and metabolic pathways (fatty acid metabolism, oxidative phosphorylation (OXPHOS), cholesterol homeostasis), xenobiotics and reactive oxygen species stress responses, and EMT/stemness pathways. White patients showed enrichment in canonical oncogenic pathways (MYC, E2F, mTORC1), the *TMPRSS2*-*ERG* signature, and biosynthetic processes (proteasome, spliceosome, embryonic stem cell signatures). Despite differences in protein counts between the adjusted and unadjusted analyses (**Figure S6A, Table S6B-C**), pathway enrichment overlap was significant (Jaccard similarity = 60.2%).

A summary of ancestry-associated pathways highlighted upregulated androgen response, EGFR signaling, and xenobiotic metabolism in Black patients, compared to enrichment in glycolysis/gluconeogenesis, mTORC1, and DNA replication/repair in White patients. Elevated DNA repair proteins in White patients include mismatch repair (MSH2/3/6, MLH1), nucleotide excision (ERCC5), base excision (APEX1, POLB), double-strand break repair (XRCC5/6, PARP1), and replication factors (MCM3/4/7). These patterns remained robust after adjusting for age and PSA (**Figures 4C, S6B, Tables S14D-E**).

Transcription factor networks further revealed distinct AR-associated hubs (**Figure 4D)**: NKX3-1, STAT1, PURA, and TBX3 were elevated in Black patients; NCOA1, FOXP1, RELA, and ERG, in White patients. Immune regulators were also featured: STAT1, IRF6, and SP100 were elevated in Black patients, alongside IRF3, CBFB, GTF2I, HMGB1, and TSC22D1 in White patients. Integration with an AR cistrome^42^ identified 65 differentially abundant AR-regulated proteins (adj. p < 0.05), with two-thirds elevated in Black patients, including CAMKK2^43^, ERBB2^44^, INPP4B^43^, and ALDH1A3^45^ (**Figures S6C-D**). In White patients, elevated AR-regulated proteins included SOX9, LDHA^46^, NDRG1^46^, and SMARCA4^47^.

Given the higher prevalence of *TMPRSS2-ERG* fusions in White patients, we evaluated fusion-associated alterations. A panel of 23 ERG-responsive proteins accurately and significantly distinguished ERG-positive from ERG-negative tumors across independent proteomic^23^ and transcriptomic^48^ cohorts (adj. p < 0.05; ρ ≥ 0.837, p < 1.00E^-4^). Sparse-partial least squares discriminant analysis confirmed this robust ERG-amplification linked proteomic classification is conserved across >600 tumors from CPDR-APOLLO and external cohorts (**Figures S6E-G)**.

### Ancestry-associated phosphoproteomic alterations and kinase signaling

To evaluate the influence of CNA on phosphoproteome dynamics, we first assessed CNAs-linked phosphorylation changes. We observed significant *trans*-effects (CNAs affecting phosphorylation of ≥5 proteins) in 4% of the genes we tested, primarily involving recurrent CNAs at 5q12, 8q22 and 13q14 (**Figure S7A**). Among CNA-phosphoprotein pairs, 5.2% showed significant correlation (|ρ| > 0.2; p < 0.05; **Figures S7B-C**) with POTEF phosphorylation showing the strongest association. Two genes, *STK39* and *ZC3H18*, showed significant *cis*-correlations where CNA had a concordant effect on both protein and phosphoprotein abundance (ρ > 0.24; p < 0.05).

Ancestry-stratified analyses of kinase and substrate genes revealed more frequent deletions of the kinase genes *GSK3B* and *CDK7*, and the substrate gene *MAP1B* in Black patients. Conversely, White patients showed increased alteration of the kinase genes *PRKDC*, *MAPK3*, and *AURKB*, and deletion of the substrate gene *HMGN1* (**Figures S7D-E**).

Kinase-Substrate Enrichment Analysis (KSEA) identified significant ancestry-associated differences in kinase activity (**Figure S7F-G**). Black patients showed higher activity of CHEK2 and CSNK2A1/CK2α, while White patients exhibited higher activity of PRKD1, CAMK2D/B, CDK1/2, and RAF1 activities. These distinctions were highlighted by CNA status; for example, increased phosphorylation of CDK2 substrates in White patients co-occurred with *CDK2* amplifications, whereas elevated HMGN1 phosphorylation in Black patients corresponded to higher CK2α activity and fewer *HMGN1* deletions. (**Figures 4E, S7H)**.

Post-translational Modification Signature Enrichment Analysis (PTM-SEA) further identified additional ancestry-specific phosphorylation signatures^49^. Black patients were enriched for signatures related to epigenetic remodeling (HDAC: MS-275, Trichostatin A; EZH2: GSK126; G9a: BIX-01338), receptor tyrosine kinase (RTK) signaling (EGFR: erlotinib, SRC: dasatinib), and inflammatory signatures (angiotensin II). Conversely, White patients were enriched in pathways involving cell cycle regulation (CDK1/2), MAPK signaling (MAPK12, trametinib, vandetinib, and pazopanib), and epigenetic plasticity (UNC-321, zebularine) (**Figures 4F, S7I, Tables S6F-G**).

### Integrative multi-omics gene-set analysis reveals ancestry-associated pathway distinctions

We used Multi-Omics Gene-Set Analysis (MOGSA)^50^ to identify the contribution of genomic, proteomic, and phosphoproteomic features to gene set enrichments. By applying generalized linear models to single-sample gene-set scores (GSS), we identified 56 significant pathways (p<0.01) that stratified into five largely ancestry-associated subtypes (**Figure 5A, Table S7A**). White-enriched subtypes (1 and 2) highlighted apoptosis, mitochondrial function, cell adhesion/migration, and epigenetic silencing (Cluster 1); RNA splicing, Polycomb mediated repression, aging and apoptotic programs (Cluster 2); and stem-like, cytoskeletal remodeling, together with innate and adaptive immune signatures, including T cell associated pathways (Cluster 3). Black-enriched subtypes (3, 4 and 5) were characterized by extracellular matrix remodeling (ECM), epithelial differentiation, and germline-associated programs (Cluster 4), and developmental signaling, DDRs, and epigenetic signatures (Cluster 5).

**Figure 5.**
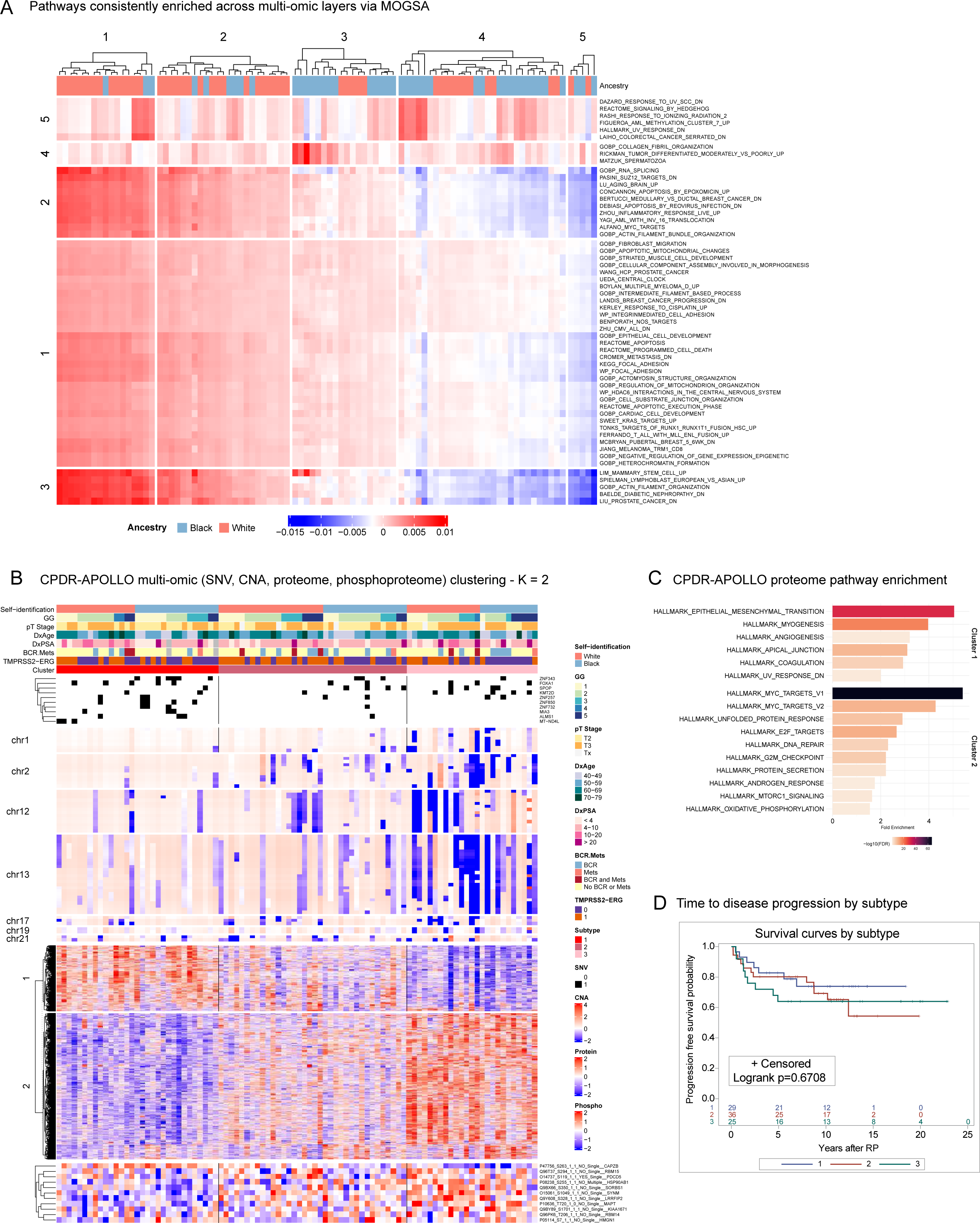
Integrated multi-omics analyses. (**A**) Heatmap of MOGSA pathways (GSS p < 0.01 in ≥50% of samples) across SNV, CNA, protein, and phosphoprotein data, hierarchically clustered based on Euclidean distance and complete linkage. Positive scores (red) indicate activation; negative scores (blue), repression. (**B**) iClusterBayes multi-omics clustering of CPDR-APOLLO prostate tumors by subtype assignments, race, clinical-pathologic, and *TMPRSS2-ERG* fusion annotations. Rows show top subtype distinguishing SNV, CNA, proteomic, and phosphoproteomic features. **(C)** Top enriched pathways per protein cluster based on fold-enrichment, colored by significance (-log_10_ adj. p < 0.05). **(D)** Univariate Kaplan-Meier survival curves for time to BCR or metastasis by molecular subtype.

### Integrative multi-omics analysis reveals distinct prostate cancer subtypes

To understand molecular heterogeneity independent of patient ancestry, we used *iClusterBayes*^51^ to jointly analyze genomic, proteomic, and phosphoproteomic data without prior filtering for differential alteration. Our analysis identified three distinct molecular subtypes (at K=2, according to Bayesian Information Criterion and deviance ratio), each with a balanced racial composition and a unique genomic and proteomic profile (**Figures 5B, S8A-B**). Subtype 1 is defined by a higher mutation frequency in *FOXA1*, *ZNF343,* and *ZNF850*. Its protein expression profile showed an enhanced mesenchymal phenotype characterized by stromal remodeling and stress-adaptive pathways but suppressed proliferation. Subtype 2 exhibited an intermediate, mixed expression profile. Subtype 3 featured more frequent *KMT2D* mutations and extensive CN losses on chromosome 13q, affecting *FOXO1, RB1*, *SETDB2*, and *RNASEH2B*. Within Subtype 3, Black patients had increased deletions at chromosome 2q (*ERCC3, LRP1B*), while White patients had 12p losses (*KRAS*, *RECQL*). Its protein profile was the inverse of Subtype 1, with highly active cell cycle, metabolic, and transcriptional programs, indicating a well-differentiated, highly proliferative tumor. (**Figure 5B-C, Tables S7B-C**). Additional subtype-defining alterations include deletions at 1q (*PTPN14, ARID4B*), 19p (*RNASEH2A*), 17p (*POLR2A* and *TP53*) and 21q (*TMPRSS2-ERG)*.

Overall, subtype distinctions were primarily driven by CNAs and protein expression, with less influence from SNVs and phosphoproteins. Excluding the phosphoproteomic data yielded nearly identical subtype classifications, though certain phosphosites (CAPZB(S263), PDCD5(S119), and SORBS1(S350)) suggested CNA-driven *trans*-effects (**Figures S8C-F, Tables S7D**). Kaplan-Meier (KM) analysis indicated faster disease progression in Subtype 3, though not statistically significant (**Figure 5D**).

We validated our findings by applying the same iClusterBayes model to publicly available TCGA PCa data, yielding three comparable subtypes (**Figures S8G-I, Tables S7E-F**). Shared CNA alterations included 2q, 13q, 17p, and 21q; TCGA additionally showed 10q (*PTEN*) deletions. Subtype-specific pathway enrichments were largely concordant across cohorts. Our Subtype 3 aligned with TCGA Subtype 2, sharing enrichment in proliferative pathways (MYC, E2F targets), DNA repair, OXPHOS, and G2M checkpoint signaling, though notably lacking androgen response signatures (**Figure S8J-K, Tables S7C**). Our Subtype 1 partially matched TCGA Subtype 1, with shared mesenchymal, stromal, and inflammatory signatures. Our Subtype 2 resembled TCGA Subtype 3, both having mixed, transitional profiles. Clinical outcomes were also consistent across cohorts, with TCGA Subtype 2 showing trends toward faster progression (**Figure S8L**).

### Prognostic relevance of CNAs

Using multivariate Cox proportional hazards (CoxPH) models, we evaluated the prognostic relevance of ancestry-specific and common CNAs, proteins, and phosphoproteins, adjusting for age, PSA at diagnosis, and percent African ancestry (**Figure S9**). Analyses of CNA identified several prognostic hotspots **(Figure S9A-B, Tables S8A-C)**. In Black patients, these included 2q22 (*LRP1B, ARHGAP15*), 3q13 (*GSK3B*), 4q26-34 (*HPGD*), and 13q13-14 (*FOXO1*). In White patients, prognostic deletions were found at 2p13 (*TET3*), 3q26 (GNB4), 8p23 (*GATA4*), 10p11 (*ZEB1*), 12q24 (*CUX2*), 16p13 (*GRIN2A*), and 16q12 (*CHD9*). We also identified progression-associated loci that were independent of ancestry including 2q14 (*ERCC3*), 2q22-37 (*ACVR2A*, *GIGYF2*), 4q28, (*FGF2*), 7q34 (*BRAF*), 8p11 (*SFRP1*, *IKBKB*), 13q12-13 (*LATS2*), and 17q11.2 (*SUZ12*). These findings were supported by either KM analyses, AUC performance, or both **(Figures 6A, S10A-C, Table S8D)**. External validation using TCGA data confirmed survival associations for two White-enriched genes (*GRIN2A*, *GNB4*) and three universally altered genes (*GIGYF*, *MTMR6*, *SMAD9*). No significant associations were observed for Black-enriched CNAs (**Figures S10D-E**), likely due to the limited number of Black patients in this dataset.

**Figure 6.**
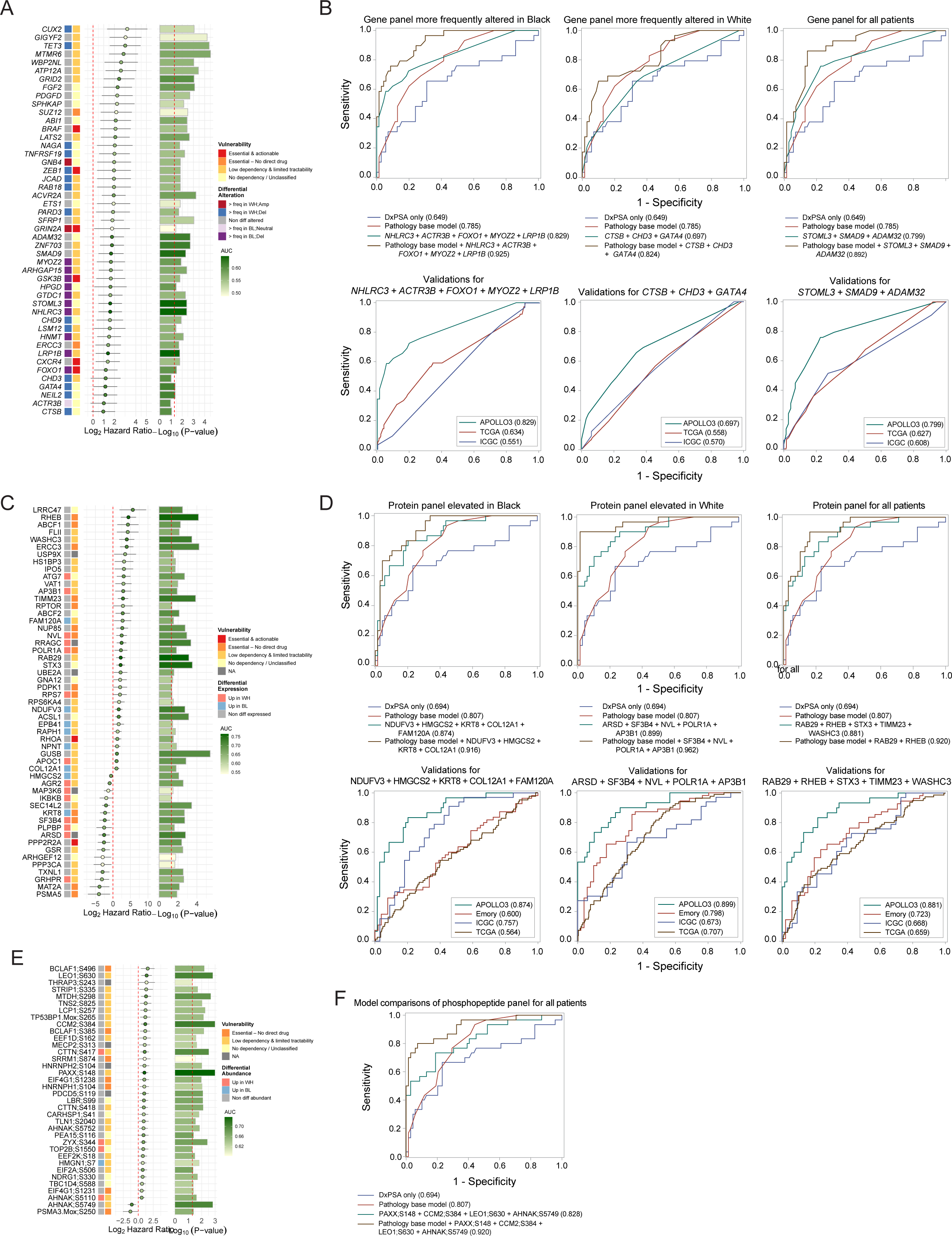
Prognostic associations of sCNA, protein, and phosphoprotein alterations. Forest plots of multivariate CoxPH models (adjusted for age, PSA, and percent African ancestry), depicting association between sCNA events (**A**), protein (**C**), or phosphoprotein (**E**) abundance and BCR or metastasis, with HRs (dotted line: Log₂ HR = 0), 95% confidence intervals, differential alteration status, targetability, and -log10 p value significance (dotted-line: p = 0.05). Fill color of points and bar-plots denotes AUC. Predictive models of sCNA **(B)**, protein **(D)**, and phosphoprotein **(F)** features selected by AUC from universally- or race-enriched subsets, compared alone or together with pathology or PSA models (top panel), and validated in independent datasets (B: TCGA and ICGC; D: Emory, ICGC, and TCGA).

To refine prognostic accuracy, we developed ancestry-specific CNA panels. The panel for Black patients (*NHLRC3*, *ACTR3B*, *FOXO1*, *MYOZ2, LRP1B*) and a diverse ancestry panel (*STOML3*, *SMAD9, ADAM32*) demonstrated high accuracy (AUC ∼ 0.88), outperforming PSA and pathology-based models. The White-specific panel achieved moderate accuracy (AUC ∼ 0.7). Validation in independent datasets (TCGA, ICGC^21^, and Emory^52^) showed reduced performance for all panels (**Figure 6B**).

#### Prognostic relevance of proteomic features

We analyzed proteomic data to identify protein sets associated with PCa progression or protection. Our findings, validated by KM survival analyses, revealed distinct sets of prognostic proteins that were either ancestry-specific or ancestry independent (**Figures 6C, S9C-D, Tables S9A-E**). Progression-associated proteins elevated in Black patients were involved in metabolism and mitochondrial function, cytoskeletal organization, membrane and vesicular trafficking, and immune modulation. Protective proteins included HMGCS2 and KRT8. Progression-associated proteins elevated in White patients regulated cell signaling (RRAGC), ribosomal biogenesis, apoptosis (BIRC6), and stress response, vesicle trafficking, and metabolism (APOC1). Protective proteins in this group were involved in metabolic balance, stress-activated signaling, and mRNA splicing (**Figures S11A-D, Table S9F**).

Proteins with similar abundance across ancestries and with prognostic relevance were involved in cell cycle and DNA repair, PI3K-AKT/mTOR signaling (RHEB), cytoskeletal regulation, and vesicle trafficking. Other notable functions included protein ubiquitination and degradation, nuclear and mitochondrial processes and diverse metabolic functions (ASCL1). Protective proteins in this group included regulators of metabolic and redox balance, cytoskeletal remodeling (ARHGEF12), protein degradation, and vesicle trafficking (**Figures S11E-F**). Validation using TCGA mRNA data confirmed many of the protein associations for White-elevated and ancestry-independent proteins, but only HMGCS2 among Black-elevated proteins (**Figures S12A-C**). While most protein and mRNA associations were concordant, some like ATG7 and MAP3K6, showed discordant effects where protein and mRNA levels had opposing prognostic values (**Figures S12D-E**).

We developed ancestry-specific and universal protein panels that achieved high predictive performance (AUC > 0.8), outperforming PSA and pathology models. Combining these panels with pathology data further improved performance (AUC > 0.9; **Figure 6D**). External validation showed moderate performance (AUC: 0.67-0.76)^23^. As expected, the White-specific (ARSD, SF3B4, NVL, POLR1A, and AP3B1) and universal (RAB29, RHEB, STX3, TIMM23, and WASHC3) panels performed best in predominantly European ancestry cohorts (Emory, TCGA), confirming their robustness across datasets.

### Prognostic relevance of phosphoproteomic alterations

Phosphoproteomic profiling identified both ancestry-specific and universally abundant progression-associated phosphosites (**Figures 6E, S9E, S13, Tables S10A-D**). Universally abundant phosphosites included PDCD5(S119), linked to apoptosis via TP53 activation^53^, PAXX(S148), mediating chemoresistance via XRCC6/Ku70 interactions^54^, and EIF4G1(S1231), a targetable node for MET inhibition^55,56^. PTM-SEA further showed overlaps with key cancer pathways and drug perturbation signatures (**Table S10E**). For example, increased phosphorylation of CTTN(S417/S418), and EEF1D(S162), indicated enhanced MAPK12 activity, while AHNAK(S5110), aligned with PI3K/mTOR inhibition, and NDRG1(S330) mapped to AKT/PKC inhibitor signatures. HNRNPH1/2(S104) phosphorylation links CDK2 signaling to cell-cycle and RNA processing, while the phosphorylation of PEA15(S116) and CARHSP1(S41) points to inflammatory regulation via angiotensin II pathways. Additional overlaps linked phosphosites to angiogenesis (LBR(S99), SRRM(S874), and AHNAK(S5752) with pazopanib) and epigenetic regulation (LEO1(S630) with UNC-0321) ^57^.

From these findings, we developed a four-phosphoprotein panel (PAXX(S148), CCM2(S384), LEO1(S630), and AHNAK(S5749)) that predicted disease progression with high accuracy (AUC = 0.83), which improved further when combined with pathology features (AUC = 0.92; **Figure 6F**). The consistent biological pathways identified through multi-omics integration ultimately highlight the ancestry-associated signaling differences driving disease progression, linking specific molecular findings to clinical outcomes.

## DISCUSSION

Our integrative proteogenomic study of localized PCa in a racially balanced, equal-access cohort addresses the critical gaps in understanding how molecular differences contribute to racial disparities in clinical outcomes. By profiling genomic and proteomic changes simultaneously, we uncovered ancestry-specific biological differences and novel molecular subtypes. These findings emphasize that uniform diagnostic and treatment approaches are insufficient; precision oncology must incorporate ancestry-associated molecular features to reduce disparities.

### Genomics

Our analysis of SNVs and CNAs revealed both established and novel ancestry-associated genomic drivers of PCa. We confirmed a higher prevalence of *SPOP* and *FOXA1* mutations^15,58–60^ and *ZFHX3* frame-shift deletions^60,61^ in tumors from Black patients, and *TP53* in those from White patients, reaffirming their etiologic roles^15,58,59,62,63^. Newly identified hotspots, including *HYAL3*, *ADAMTS2*, and several zinc-finger protein genes, suggest additional mechanisms of tumorigenesis in African-ancestry tumors. Recurrent *ADAMTS2* mutations mirror recent findings in Nigerian PCa patients^64^. Consistent with broader genomic diversity, Black patients showed higher mutation counts and transversion-driven signatures^65^. Conversely, White patients showed more aging-associated signatures, reflecting their older age at diagnosis^66^. These distinct mutational processes underscore the need for ancestry-aware biomarker development and clinical trial stratification

CNA analysis revealed both shared and divergent events. Common alterations across ancestries include 8q gains, 8p/13q deletions, with fewer *PTEN* deletions and *TMPRSS2-ERG* fusions in Black patients^14,15,59,61,62,67,68^. White patients were enriched for focal deletions at loci involving key tumor suppressors at 3p13, 8p23, 12p12, 16q22-24, and 17p11-13. Black patients more often showed focal amplifications, and recurrent deletions at 2q22, 3q13 (encompassing LSAMP, previously linked to aggressive PCa in African American men^14^), 5q12-14, and 13q14, with mixed events at 7q36 and 8q24. These results extend prior ancestry-associated observations, including 1q41-43 deletions in African Caribbean cohorts^17^, 2q22^63^ and 3q13^14,69^ deletions in Black patients, and recurrent 3p13 and 8p23 deletions in White patients^34,35,70,71^. The strong concordance across studies suggests convergence of ancestry-specific drivers with prognostic and therapeutic relevance, despite cohort and platform differences.

Ancestry-associated eQTLs introduce an inherited regulatory layer that significantly influences the tumor proteome, acting independently of somatic CNAs and SNVs. This germline layer modulates key pathways including AR axis-coupled fatty acid metabolism (ACSM1^72^) and multi-drug resistance (ABCC4^73^), carcinogen detoxification (GSTM1/3^73^), and cell migration/invasion (ARHGAP1^74^). Specifically, LGALS8, a galectin involved in cell adhesion and immune modulation, was associated with poorer prognosis, whereas FECH, a mitochondrial enzyme in heme biosynthesis and oxidative metabolism, showed a protective effect. Because these inherited regulatory effects can influence treatment response and resistance, incorporating eQTL status into risk models and clinical trial stratification may refine patient selection and guide personalized therapy. Recent large-scale WGS analyses^75^, demonstrating that ancestry-enriched germline variants can predispose tumors to acquire distinct somatic drivers, further support this approach.

### Proteomics and phosphoproteomics

Previous multi-omics studies of PCa have emphasized genomics and transcriptomics, with limited proteomic and phosphoproteomic coverage, especially in racially diverse^22,23,76^, localized disease^77,78^ cohorts. Our study extends this scope by adding proteomic and phosphoproteomic layers, providing functional resolution that capture protein and kinase activity changes not evident from genomic data alone. We identified ancestry-specific features, consistent with previous transcriptomic findings, such as those from the Durham Veterans Affairs Health Care System^79^. This included results showing enhanced androgen response, cholesterol homeostasis, EMT, and lipid-fatty acid metabolism in Black patients, while revealing novel differences such as heightened mTORC signaling and distinct inflammation-associated pathways in White patients. This concordance across molecular layers strengthens our biological findings and their clinical interpretability, highlighting the value of proteomics in translating genomic variation into functional pathways alterations with diagnostic and therapeutic potential.

### Molecular subtypes

Using iClusterBayes to integrate multi-omic data, we identified three reproducible molecular subtypes that provide a new framework for risk stratification with clinical potential. These subtypes were validated externally in and independent TCGA cohort. Subtype 1, a mesenchymal phenotype defined by stromal remodeling, EMT, and stress adaptive pathways, suggests testable hypothesis for responsiveness to EMT-inhibiting therapies^80,81^. Subtype 2 represented a transitional state with mixed features. Subtype 3 was a proliferative subtype, characterized by heightened cell cycle progression, DNA repair, and transcriptional reprogramming, with deletions on 13q, 12p, and 2q22. This subtype showed a trend toward accelerated progression (though not statistically significant). Interestingly, IND3-enriched patients who are significantly associated with disease progression risk are also characterized by cell cycle regulation and proliferative pathways. Notably, these observation aligns with recent reports that prostate tumors with elevated proliferation and cell cycle signatures, particularly E2F targets, G2M checkpoint, MYC targets, and mitotic spindle pathways, are associated with higher risk for BCR^82^ or poorer overall survival^83^. These findings reinforce the notion that highly proliferative PCas represent more aggressive disease states and may harbor vulnerabilities to DNA repair inhibitors or cell-cycle checkpoint blockade^84,85^. Importantly, ancestry contributed to subtype composition: Black patients more frequently carried 2q22 deletions within Subtype 3, while White patients exhibited 12p deletions. Although phosphoproteomics contributed modestly to subtype resolution, enrichment of specific phosphosites in Subtype 3 indicates added value for refining molecular risk. Collectively, these subtypes offer a reproducible classification framework for improved risk stratification and hypothesis-driven therapeutic exploration.

### Prognostic models

Our integrative analysis shows that ancestry-associated CNAs, proteins, and phosphoproteins are both biologically distinct and clinically informative. The ancestry-specific biomarker panels we developed significantly outperformed conventional models, highlighting the clinical value of molecular data in risk prediction. We found that White patients harbored prognostic CNA hotspots in tumor suppressors such as *TET3*, *GATA4*, and *GRIN2A*, while Black patients showed progression associated alterations in *FOXO1*, *GSK3B,* and *LRP1B*. Furthermore, alterations such as *LRP1B* deletion in Black patients and *GRIN2A* loss in White patients are not only prognostic but also predictive of therapeutic response^86–89^, suggesting their dual utility. The broader *trans*-acting effects between CNAs and proteins further suggest that prognosis is influenced by rewired regulatory networks converging on potentially druggable pathways. Our findings also highlight the importance of large, diverse cohorts to validate infrequent but potentially impactful genomic drivers such as *GIGYF2* and *MTMR6*.

Proteomics added valuable prognostic information by identifying ancestry-specific vulnerabilities. Markers of mitochondrial metabolism, ECM remodeling, and immune regulation were enriched in Black patients, while proliferative and apoptotic regulators were elevated in White patients, correlating with poor outcomes. We identified both known aggressive markers elevated in Black patients (RAPH1^90^, DOCK4^91^) and novel candidates (COL12A1, TSPAN31, CMTM6). For proteins elevated in White patients, we found new markers (RRAGC, PARD6B, and SLC25A5) and confirmed known associations such as BIRC6^92^, ATG7^92^, and APOC1^93^ with aggressive disease. We also identified a set of universally abundant prognostic proteins involved in DNA repair, PI3K–mTOR signaling, and ubiquitination, including ERCC3, CDK11A, RPS6KA4, and UBE2A, suggesting potential targets for ancestry-agnostic interventions.

Phosphoproteomics further refined risk stratification by linking phosphorylation events to disease progression. We identified phosphosites associated with chemoresistance^94–96^, chromatin remodeling^57^, and kinase-driven signaling^53^, many of which mapped to the same pathways implicated by CNAs and proteins. The enrichment of specific phosphosites in the more aggressive Subtype 3 reinforces the value this profiling for identifying high-risk patients. Crucially, several phosphosites, including those regulated by CK2α, CHEK2, and CDK2, point to potentially actionable kinase dependencies, suggesting that phosphoproteomic data can guide therapeutic targeting as well as prognostication.

### Ancestry-specific biological differences with therapeutic implications

Our integrative analysis shows that ancestry-associated molecular differences reflect not isolated events, but convergent systems-level effects on key biological pathways, revealing testable therapeutic vulnerabilities and resistance mechanisms often missed by single omics analysis.

Androgen receptor regulation and steroid hormone pathways were prominent all across omics-layers. Black patient tumors showed stronger androgen/estrogen responses and cholesterol metabolism, with more AR-regulated proteins and denser transcriptional networks, including NKX3-1, STAT1, PURA, and TBX3. White patient tumors were enriched for, NCOA1, FOXP1, RELA, and ERG networks as well as *TMPRSS2–ERG* fusions. These patterns suggest ancestry-associated differences in resistance to AR-targeted therapies, raising hypotheses that co-targeting AR with cooperating transcriptional regulators such as STAT or chromatin modulators could prevent or overcome resistance.

Metabolic reprogramming was a consistent finding. Black patient tumors displayed upregulated OXPHOS, and fatty acid metabolism, indicating potential sensitivity to mitochondrial inhibitors or 5′-AMP-activated protein kinase (AMPK) activators such as metformin, which are under clinical evaluation for PCa^97–99^. Enzymes such as FASN, ACC, ACLY, and CPT1A also emerge as promising targets for further investigation^100–103^. In contrast, White patient tumors relied more on glycolysis, ribosomal biosynthesis, and proteasomal activity, pointing to vulnerabilities like proteasome inhibitors like bortezomib^104^. These divergent metabolic states provide a strong basis for testing ancestry-specific targeted treatments.

We also observed distinct, ancestry-specific perturbations in cell-cycle and DDR pathways. The indel-based IND3 mutation signature, indicative of mismatch-repair-associated replicative stress, was enriched in White patients and correlated with aggressive disease, implicating early DDR activation. Tumors from White patients also showed MYC and E2F-driven proliferation, elevated CDK1/2 activity, and phosphorylation of key regulators, suggesting potential sensitivity to CDK or cell-cycle checkpoint inhibitors. Conversely, higher inferred CHEK2 and CK2α activities in Black patients indicated enhanced DDR signaling and pro-survival adaptation ^105–107^. While DDR alterations and MMRd-associated signatures appeared in both ancestries, White patients more frequently harbored *FANCA, PALB2,* and *RPA1* deletions, suggesting potential sensitivity to PARP inhibitors or immune checkpoint blockade^108–110^. Additional *POLR2A* deletions may further sensitize these tumors to targeted therapies^111,112^. These ancestry-linked DDR alterations align with our prior finding of enriched germline variants in DDR genes among African American men with PCa, particularly in RAD-associated pathways^113^. Recurrent 13q loses encompassing *RNASEH2B, RB1*, and *BRCA2* across ancestries may modify responses to PARP inhibition, highlighting the clinical complexity introduced by convergent genomic alterations^114^. The frequent elevated expression of DDR proteins in White patients also suggests enhanced DNA repair capacity and potential for treatment resistance^115,116^. These findings underscore the value for integrative multi-omics profiling for resolving ancestry-associated vulnerabilities. Integrating phosphoproteomic data, for example, pinpointed elevated PAXX;S148 phosphorylation and its interacting partner XRCC6/Ku70, enabling the identification of potential precision targets^54^.

Our analysis revealed convergent alterations implicating both the PI3K/AKT/mTORC and MAPK pathways. White patients frequently carried *PTEN* and *YWHAE* deletions with elevated IKBKB and YWHAZ proteins and enhanced MAPK and mTORC1 activity^96,117,118^, together with higher inferred RAF1^119^, PRKD1^120^ and CAMK2D/B activation, consistent with enhanced proliferation and motility^121^. Conversely, Black patients showed elevated ERBB2, INPP4B, CAMKK2, and GREB1, implicating PI3K/AKT-AR cross-talk^108–110^. These profiles raise hypotheses for evaluating targeted PI3K, AKT, mTOR^111,122,123^, MAPK^124^, or ERBB2^112^ inhibitors in ancestry-aware contexts.

Multi-omics integration revealed recurrent ancestry-associated disruptions in epigenetic regulation that may prime tumors for lineage plasticity even before treatment. Black patients were enriched for stem-like pathways and CNAs in *HDAC2*, *COBRA1*, *FOXP3*. Conversely, White patients displayed Polycomb and SWI/SNF repression signatures. These perturbations intersected with oncogenic signaling and stress responses, likely reshaping enhancer landscapes, reprogramming the AR cistrome, and weakening lineage-restrictive transcription^125^. Phosphoproteomics further implicated kinase activities modulating HDAC and EZH2 in Black patients, and G9a and MAPK-RTK signaling in White patients. The detection of these changes in untreated tumors suggests an intrinsic predisposition to lineage plasticity and neuroendocrine transitions before exposure to AR inhibitors^126^. These findings support hypotheses that early interception of chromatin reprogramming, directly or in combination with kinase or AR-directed therapies, could blunt lineage plasticity and therapy resistance^127^.

Our integrative analysis results consistently aligned with an ancestry-associated divergence in immunoregulation and inflammation^128,129^, suggesting these distinct immune states, encompassing both innate (interferon/viral response) and adaptive (T-cell) programs, likely influence immunotherapy response. Black patients showed enrichment of immune-related programs, including STAT1 and IRF6, together with CNA affecting FOXP3 and HDAC2. Prognostic analysis further linked immune differences to outcomes, highlighting the elevated immune-modulatory protein DOCK4^130^ and prevalent deletion of immunoregulatory protein *LRP1B*^86,131^. In contrast, White patients were enriched for T-cell associated pathways, chemokine signaling, and interferon α/γ response with elevated immune regulators like IRF3. These findings support hypotheses that tailoring immunotherapy strategies to ancestry-associated immune profiles for more effective outcomes.

Stromal/cytoskeletal remodeling and EMT signals were prominent in Black patients, supported by enriched ECM remodeling and epithelial differentiation, together with elevated COL12A1, NPNT, DOCK4, and RAPH1 expression. Conversely, White patients were enriched for proliferative epithelial and lineage plasticity programs, driven by ERG and SOX9 networks. The overlap of these patterns with iClusterBayes Subtypes suggests that EMT- and plasticity-associated states may confer therapy resistance, especially under androgen deprivation, warranting the evaluation of stromal- or EMT-targeted approaches in specific molecular subtypes.

### Limitations

This study provides one of the first racially balanced, integrative proteogenomic analysis of localized PCa, leveraging a unique equal access cohort with detailed clinical annotation and long-term follow-up. Key strengths include the equal-access, racially balanced cohort, which minimizes socio-clinical confounding; the depth of multi-omics integration, spanning genomics, proteomics, and phosphoproteomics; and the use of covariate adjustment, which reduces residual clinical confounding such as age at diagnosis. Limitations include modest sample size, which limited detection of subtle ancestry specific differences and rare drivers with a long tail distribution^132^. Despite covariate adjustment, residual confounders cannot be fully excluded. Validation was further constrained by limited matched proteomics datasets and demographic imbalances in external cohorts.

In summary, this study provides evidence that ancestry-associated proteogenomic alterations converge on key oncogenic programs in localized PCa. We delineate both universally conserved and ancestry-specific molecular mechanisms of progression and therapeutic vulnerability. By integrating multi-omics data in a racially balanced cohort, we provide a framework for ancestry informed prognostic models and precision treatments aimed at reducing outcomes disparities. Given our modest sample size, these therapeutic implications should be viewed as hypothesis-generating, requiring validation in larger, ancestrally diverse cohorts. Future research must include functional studies to elucidate the causal mechanisms of ancestry-associated differences. Additionally, the application of single-cell and spatial multi-omics approaches will be crucial to resolve intratumoral heterogeneity and clarify the link to ancestry-specific clinical outcomes.

## METHODS

### Cohort selection

Prostate specimens were collected from treatment-naïve patients who were treated by RP between 1998 and 2018 who underwent radical prostatectomy (RP) surgery at the Center for Prostate Disease Research (CPDR), Walter Reed National Military Medical Center (WRNMMC), and had provided written consent for the use of their biospecimens, clinical data, or both, for research under protocols (DBS.2020.094, WRNMMC #393738 and USUHS #GT90CM, respectively) approved by the Institutional Review Board. Patients, who comprise active-duty service members, retirees, and beneficiaries, were selected based on self-identified race (55 White and 57 Black), and matched for pathological tumor stage and grade. Whole-genome sequencing was performed on fresh-frozen specimens from 103 patients (52 Black and 51 White), using matched germline DNA from blood samples for accurate somatic mutation detection. The final proteogenomic dataset included phospho- and global-proteomics for 101 patients (48 Black, 53 White). An additional proteomic baseline was established by analyzing normal FFPE prostate tissues from a subgroup of 25 of these patients (10 Black and 15 White).

### Tissue processing at time of surgery

Histologically defined prostate tumor tissues were isolated from 6-7 µm thick sections of *ex-vivo* tumor biopsy specimens collected immediately following RP and embedded in OCT prior to freezing. Hematoxylin and Eosin (H&E)-stained sections were reviewed by a certified pathologist (I.A.S.) to determine Grade Group (GG) and percent composition of tumor regions selected for genomic and proteomic analyses. For both WGS and proteomics, defined regions of tumor and normal tissues were isolated manually from unstained tissue sections by macro-dissection using a #10 blade scalpel (Graham-Field 2975#10 and Integra #4-410). Total tissue area isolated per patient was > 0.5 cm^2^, measured by ImageJ (NIH) or NIS-Elements D (Nikon) software. Pathologic Gleason score and pathologic T stage were defined by evaluation of whole-mounted prostate sections.

### DNA isolation and library preparation

Tumor DNA was isolated from fresh frozen tumor tissues using the DNeasy Blood and Tissue Kit; germline or reference DNA, from matched peripheral mononuclear cells of whole blood using the PAXgene Blood DNA Kit (Qiagen, Gaithersburg, MD). Genomic DNA was measured by using Qubit DNA High Sensitivity Assay Kits on the Qubit Fluorometer 3.0 (Thermo Scientific). If blood was unavailable, DNA was isolated instead from pathologically defined non-tumor biopsy tissue for reference. A minimum input of 120 ng normalized gDNA was added at 55 µL volume into wells of a 96 microTUBE plate and sheared using the LE220 Focused-ultrasonicator (Covaris). Sheared DNA was assessed by a fragment analyzer (Agilent Technologies) to ensure adequate DNA insert size (350 bp insert, ≤ 2 x 101 bp). Sequencing libraries were prepared from 500 ng fragmented DNA using the Illumina TruSeq DNA PCR-free Library Preparation Kit with adaptations for automation on a Hamilton STAR Liquid Handling System. Illumina TruSeq Single Indexes (Set A 20015960, Set B 20015961) were used for barcoding. Adapter-ligated libraries were pooled at 2 µM ratios and sequencing was performed on Illumina’s GAIIx, HiSeqX, or NextSeq500 platforms. Sequences were normalized prior to analysis to avoid discrepancies between sequencing platforms. Paired-end reads were sequenced at a minimum depth of 30X mean coverage across the entire genome, and most tumor samples have at least 80X coverage.

### Whole genome sequencing sample concordance and variant calling

All WGS samples were initially processed through the Illumina Sequence Analysis Software Resequencing Workflow (version 6.19.1.403 + NSv6) as described earlier in Soltis et al.^133^. All samples were profiled for somatic mutations, rearrangements, and copy number variations (CNV) using validated pipelines. This workflow aligned sequencing reads to the human reference genome (NCBI GRCh38 with decoys) with the Isaac aligner (version Isaac-04.17.06.15)^134^ and called germline variants (SNVs and short indels) with Strelka2 (version 2.8.0)^135^, structural variants (SVs) with Manta (version 1.1.1)^136^, and copy number variants with Canvas (version 1.28.0.272 + master)^137^. Initial sample quality features assessed at this stage included total pass-fail reads, percent aligned reads, and total coverage depth (target ∼30X for germline specimens, ∼90X for tumors). We confirmed sample gender from chromosome X heterozygous to homozygous variant ratios with a support vector machine (SVM) classifier trained on DNA specimens of known gender.

We inferred sample ancestries from DNA evidence using *Peddy*^138^ and compared these to the self-declared race of patients. Briefly, principal component analysis (PCA) was performed on genotype calls at specific loci from 2,504 samples in the 1000 Genomes Project (1KGP). A support vector machine (SVM) classifier was trained on the resulting first four principal components (PCs), using known ancestries from the 1KGP’s five super-populations (African, American, East Asian, European, and South Asian) as training labels. Sample genotype calls from our cohort at these same loci were then mapped into the principal component space, and the trained SVM was used to predict underlying ancestries. To quantify racial admixture, we focused on PC1, as it captured the primary axis of variation between the European and African ancestry cohorts. Percent African and European ancestry were calculated from PC1 based on the distance to the PC1 means of these two reference populations^139^.

We then called somatic variants from matched tumor and normal specimens using the Illumina Sequence Analysis Software Tumor Normal Workflow (version 6.9.1.177 + NSv6), which called somatic SNVs and indels with Strelka2, somatic structural variants with Manta, and somatic copy number alterations with Canvas. Prior to running this workflow, we first verified that the matched sample DNA specimens were derived from the same individual. For this we performed pairwise sample genotype concordance check using SNVs on chromosome 1. Mismatched samples are removed from further analysis. This analysis also revealed that all expected sample pairs indeed derived from the same individuals. Tumor purity was estimated from sequence reads by Canvas within the Illumina Tumor Normal Workflow.

### Germline variant pathogenicity and somatic variant analysis

We used InterVar^140^ to classify all germline variants. We selected variants called Pathogenic (P) or Likely Pathogenic (LP) and occurring in genes reported as germline mutations by The Cancer Genome Atlas^141^. We further filtered these variants using ExAC^142^, retaining those with an allele frequency less than 1%.

For somatic SNVs and indels, we only retained variants passing all Strelka2 filters. To further control for potential false positives and/or caller artifacts, we used a panel of normals (PON) approach whereby additional somatic variant calls were dropped if they were detected in ‘‘pseudo somatic’’ call sets derived from the matched normal specimens. For this, we ran the Tumor Normal Workflow on every normal WGS dataset against a synthetic normal sample generated with *wgsim*^143^ and mimics an ∼30X depth WGS dataset with all base calls matching the hg38 reference genome. We ran *wgsim* with the following parameters to create a set of simulated WGS paired-end FASTQ files with no variants from the reference genome: -N 475,000,000 -d 420 -s 95 -e 0.0–1 150 -2 150 -r 0.0 -R 0.0 -X 0.0. We then ran the Re-sequencing Workflow on this synthetic dataset and used the resulting outputs as the ‘‘normal’’ specimen when running the Tumor Normal Workflow with the true normal WGS samples as the ‘‘tumors’’. Passing variants occurring in two or more pseudo-somatic samples formed a filter list. These SNV and indel variants were then filtered from tumor somatic variants in the cohort. Finally, we annotated these filtered somatic SNVs and indels with ANNOVAR^144^ against GENCODE v28 gene models^145^, to be used throughout the remainder of the study. Tumor mutational burden (TMB) was calculated as the sum of SNVs and indels per megabase.

### DNA copy number segments and gene-level copy number values

For somatic copy number alterations (CNAs), we retained segments called by Canvas that passed all caller filters (i.e., PASS variants) for gene-level analysis. For somatic SVs, we ran Break Point Inspector v1.7 on the Manta, calls which reexamines support for these calls from the tumor and normal sample BAM files directly. Somatic SVs not passing Manta and additional Break Point Inspector filters were discarded.

We used continuous log_2_(CN) – 1 values (logCN) as the numerical factors for Canvas segments for several analyses. Here, CN, or the normalized floating point copy number (e.g., 2.0 equals copy number 2), is equal to (2 * RC)/(DC), where RC is the mean read count per bin reported by Canvas for the segment and DC is the estimated value of the overall diploid coverage for the sample. For autosomes, we used log_2_(CN) – 1 values directly; for sex chromosomes, we adjusted the calculations on chromosomes X and Y to log_2_(CN) – 0.

High-resolution, allele-specific, gene-level CNA calls were further determined using the Battenberg package (v2.2.8).) The algorithm was applied to paired tumor and normal whole-genome sequencing data as described in Nik-Zainal et al.,^146^. Briefly, 1000 Genomes Project (1KGP) reference heterozygous SNPs were used for haplotype phasing and a- and b-alleles assignment. Genomic regions were segmented using piece-wise constant fitting, t-tests were conducted on segmented b-allele frequencies (BAFs) of copy number segments to determine clonal or subclonal copy number states. Subclonal copy numbers are modeled as mixtures of two copy number states estimated from the average BAF of heterozygous SNPs within each segment.

### Chromothripsis and Kataegis

Chromothripsis was determined by using the ShatterSeek (v11.1) package^147^. For a high confidence call, an observation of at least six interleaved intrachromosomal SVs, seven contiguous segments oscillating between two CN states, a passing result on the fragment joins test (q-value < 0.2), and a passing result on either the chromosomal enrichment or the exponential distribution of breakpoints test (q-value < 0.2). Alternatively, a high-confidence call was made based on the presence of inter-chromosomal SVs, implicating multiple chromosomal events that require the observation of at least three interleaved intrachromosomal SVs,, four or more inter-chromosomal SVs, seven contiguous segments oscillating between two CN states, and a passing result on the fragment joins test (q-value < 0.2). These events were confirmed by visual inspection of SV, CNV and LOH using a genome browser. Kataegis was analyzed by using the kataegis (v1)^148^ package using inter-mutation length 50,000 as a cutoff. Five samples out of 103 were found to have kataegis events (370, 573, 1941, 1739, 1369).

### Identification of differential SNV and copy number alteration

Differential SNVs frequencies between groups were first assessed using Fisher’s exact test, identifying genes with significantly different proportions (p <0.05). To account for potential confounding factors, Firth’s penalized logistic regression was subsequently applied, modeling each gene’s mutation status (mutated vs. wild-type) as a function of self -identified race, age, and PSA levels at diagnosis. Genes significant in either test (p, 0.05) were considered differentially altered and presented in the results.

To identify ancestry-specific, biologically relevant copy number alterations (CNAs), we used a two-pronged approach combining Fisher’s exact test and Elastic Net regression. Fisher’s exact test assessed the overall differences in CNA occurrences. Elastic Net regression provided a complementary analysis that accounted for multinomial event types, and covariates, like age, and PSA at diagnosis, and percent African ancestry ^149,150^. For Fisher’s exact test, we selected genes as differentially altered if they met the following criteria: a p value ≤ 0.1, a difference of at least six alterations (≥ 12%), and a relative difference (REL.DIFF = (BL–WH)/((BL+WH)/2) of ≥ 0.4 between Black and White samples. We evaluated the statistical significance of gene set overlaps using a hypergeometric test.^151^.

For the Elastic Net analysis, we applied the ***glmnet*** package in R. We created a matrix of copy number alteration values (-2, -1, 0, 1, or 2), excluding genes with low variance (<0.01). The CNA matrix (predictors) and covariates data frame were merged into a single object for modeling. We determined the optimal alpha parameter (0.005) through five-fold cross-validation, over a range of alpha values (0 to 0.5, by = 0.001), selecting the lambda value that minimized the cross-validated mean square error (CVM). We then used this optimal alpha in models employing **Stability Selection** and **Ensemble Methods**. In Stability Selection, we repeatedly fit the model on 500 random subsamples (90% of the data each) to calculate a stability constant for each gene, which was the number of times it was selected across all subsamples.. The ensemble method involved fitting the dataset through 200 models using the optimal alpha value over a range of lambda values (25 to 1). We then averaged each gene’s coefficient magnitude across all models to assess its importance. Genes were classified as differentially altered if they had a stability score > 0.25 and an absolute mean coefficient > 3.00E-4.

### Specimen preparation for proteomics

Tissues were collected into a single spherical cluster and transferred with a sterile pipette tip into a 150 µl labeled pressure microtubes (Pressure BioSciences Inc.,). Each microtube contained 10 µl of 1X collection buffer (100X Halt Phosphatase Inhibitor Cocktail, Thermo Scientific, diluted with LC-MS/MS-grade water).The microtubes were sealed with MicroCaps using a MicroCap Tool (Pressure BioSciences Inc.), placed into labeled 0.6 ml sterile PCR tubes (Axygen), briefly kept on dry ice, and finally transferred to -80°C for long term storage before pressure assisted protein digestion.

A total of 102 tissue sections on PEN membrane slides were manually scraped into pressure cycling technology (PCT) MicroTubes (Pressure Biosciences, Inc.) containing 10 µL LC-MS-grade water supplemented with 1x Halt Phosphatase Inhibitor Cocktail (ThermoFisher Scientific, Inc.) with an average area of 75 mm2. Microtubes were capped, placed in a 0.5 mL thin-walled PCR tube and stored at -80°C. Prior to digestion, sample buffer was adjusted to 100 mM triethylammonium bicarbonate (TEAB, pH 8.0), 10% acetonitrile. Samples were incubated in a thermocycler at 99⁰C for 30 minutes. Tubes were centrifuged briefly and then MicroTubes were removed from the PCR tubes. The MicroCaps were removed and discarded. Following the addition of 1 µL of SMART Digest Trypsin (Thermo Scientific), MicroTubes were capped with MicroPestles. Pressure-assisted lysis and digestion was performed in a Barocycler 2320EXT (Pressure BioSciences, Inc.) by sequentially cycling 60 times between 45 kpsi and atmospheric pressure at 50⁰C. Resulting peptide samples were transferred to 0.5 mL microcentrifuge tubes, vacuum dried, resuspended in 100 mM TEAB and peptide concentration was determined using a bicinchoninic acid assay (ThermoFisher Scientific).

Forty micrograms of peptide from each sample, along with a pooled reference sample (assembled from equivalent amounts of peptide digests pooled from individual patient samples), were aliquoted into a final volume of 100 µL of 100 mM TEAB. Peptides were labeled with a unique isobaric tandem mass tag (TMT) label according to the manufacturer’s protocol (TMT11plex™ Isobaric Label Reagent Set, ThermoFisher Scientific, Inc.). Each multiplex was fractionated by basic reverse phase liquid chromatography (1260 Infinity II, Agilent) into 96 fractions through development of a linear gradient of acetonitrile (0.69%/min). Concatenated fractions (36 pooled samples, representing 10% of the total peptide sample) were prepared for global LC-MS/MS analysis. The remaining 90% of peptides were pooled into 12 fractions for serial phosphopeptide TiO2 enrichment followed by iron immobilized metal ion affinity chromatography (Fe-IMAC). Briefly, concatenated peptide fractions were vacuum dried, re-suspended in TiO2 binding/equilibration buffer, and bound to TiO2 affinity spin columns (High-Select TiO2 Phosphopeptide Enrichment Kit, ThermoFisher Scientific, Inc.). The sample flow-through and washes were reserved for subsequent enrichment by Fe-NTA (nitrilotriacetic acid) affinity chromatography (High-Select Fe-NTA Phosphopeptide Enrichment Kit, ThermoFisher Scientific, Inc.).

Additionally, normal formalin-fixed paraffin-embedded (FFPE) prostate tissues, taken furthest from the isolated tumor section from 10 Black and 15 White patients, were used to establish a baseline protein readout.

### Mass spectrometry-based proteomics, phosphoproteomics and data analysis

Liquid chromatography-tandem mass spectrometry (LC-MS/MS) and analyses of TMT-11 multiplexes were performed as previously described^152^.We analyzed 36 RPLC fractions per TMT-11 sample plex, resuspended in 12.5 μL of 100 mM NH_4_HCO_3,_ by LC-MS/MS using a nanoflow LC system (EASY-nLC 1200, ThermoFisher Scientific) coupled online with a high field Orbitrap MS Fusion Lumos Tribrid MS (ThermoFisher Scientific). Briefly, each concatenated TMT fraction (5 μL, ∼500 ng) was loaded on a nanoflow HPLC system equipped with a reversed-phase trap column (Acclaim PepMap 100 Å C18, 20 mm, nanoViper, Thermo Scientific) and a heated (50°C) reversed-phase analytical column (Acclaim PepMap RSLC C18, 2 µm, 100 Å, 75 µm × 500 mm, nanoViper, Thermo Scientific), which was directly connected to the mass spectrometer. Peptides were eluted using a 120 min linear gradient from 2% mobile phase A (95% acetonitrile, 0.1% formic acid) to 32% mobile phase B at a constant flow rate of 250 nL/min. The electrospray source capillary voltage was set to 2.0 kV and the temperature to 275°C. High resolution (R=60,000 at *m/z* 200) broadband (*m/z* 400-1600) mass spectra (MS) were acquired. Subsequently, the top 12 most intense molecular ions from each MS scan were selected for high-energy collisional dissociation (HCD, normalized collision energy of 38%) and acquired in the orbitrap at high resolution (R=50,000 at *m/z* 200). The monoisotopic precursor selection mode was set to “Peptide” and MS1 peptide molecular ions chosen for HCD were restricted to charged states *z* = +2, +3, and +4. The radio frequency (RF) lens was set to 30% and both MS1 and MS2 spectra were collected in profile mode. Dynamic exclusion (t = 20s at a mass tolerance of 10 ppm) was enabled to minimize redundant selection of peptide molecular ions for HCD.

Analyses of TMT global and phosphoproteome data were performed as previously described^152^. Briefly, peptide identifications were generated by searching LC-MS/MS data against a publicly available, non-redundant human proteome database (Swiss-Prot, Homo sapiens, downloaded 12/01/2017) using Mascot (v. 2.6.0, Matrix Science) and Proteome Discoverer (PD; v. 2.2.0.388, ThermoFisher Scientific) with the following parameters: a precursor mass tolerance of 10 ppm, fragment ion tolerance of 0.05 Da, a maximum of two tryptic miscleavages, static modification for TMT reporter ion tags (229.1629 Da) on N-termini and lysine residues, and dynamic modifications for oxidation (15.9949 Da) on methionine residues. Resulting peptide spectral matches (PSMs) were filtered using an FDR of < 1.0% (q-value < 0.01), as determined by the Percolator module^153^ within PD. TMT reporter ion intensities were extracted using PD at a mass tolerance of 20 ppm. The average PSM identified per TMT-11 multiplex (12 TMT-11 multiplexes in total for CPDR-APOLLO) was 8,558 ± 756 (relative standard deviation (RSD) = 8.8%). LC-MS/MS performance and quality control were determined by analysis of a commercial human cell line digest (MSPE, MS Compatible Human Protein Extract, Digest, Promega) before and after data was acquired for each TMT-11 plex; RSD of MSPE PSM = 3.8%).

For PSMs mapping to multiple proteins, we assigned them to unique proteins using a mean squared error approach. We then determined global protein-level abundance by calculating the median log2 abundance ratios of unique PSMs. Protein-level abundance was calculated from normalized, median log_2_ transformed TMT reporter ion ratio abundances, requiring a minimum of two PSMs per protein accession. Normalized log_2_ protein-level abundances for each TMT-11 multiplex were merged, and protein-level abundances for proteins not quantified in all patient samples, but in ≥50% were imputed using a k-nearest neighbor (k-NN) strategy using the *impute* R-package^154^.

The abundances of phosphorylated (phospho)-PSMs were assembled at the level of discrete phosphosites that map to a unique protein using a tiered strategy as previously described^133^. Normalized log_2_-transformed protein-specific phosphosite abundances were calculated from the median abundance of phospho-PSMs sharing the same phosphosite and the methionine oxidation state. For phosphosites redundantly quantified as both low-and high-confidence versions, only high-confidence phosphosites were selected for downstream analyses. For phosphosites co-identified in companion global proteomic data, median log_2_-transformed, protein-specific phosphosite abundances were additionally normalized to the total protein abundance quantified in global proteome analyses. Phosphosites co-quantified in at least 75% of the cohort were imputed by k-NN using the *impute* R-package^154^.

#### Identification of proteins with significant differential CN alterations

Differential expression analysis of the global proteomics, phosphoproteomics, and validation datasets was performed using Limma (v3.52.2) in R (v4.2.1). Pathway analysis was performed using clusterProfiler (v3.14.3)^155^ with proteins differentially expressed in Black versus White groups (adj. p < 0.05). Inference of kinase activity was generated using the KSEA App (v0.99.0; PSP&NetworKIN Kinase Substrate Dataset Curated July2016 reference database)^156^ for all imputed features in the phosphoproteomics datasets. PTM-SEA pathway analysis for phosphosites was performed using ssGSEA2 (v1.0.0) and the v2.0.0 PTMsigDB database. Differential analysis adjusted for age and PSA was described by Law et al.^157^ and was performed on the subset of the cohort with available age and PSA data (n99). ERG positivity status was established by a consensus *TMPRSS2-ERG* fusion call by using MANTA, Canvas, and Genoquake from whole genome sequencing data and by a positive ERG IHC staining of the index tumor.

^158^Data from Sinha et al.,^23^ pre-processed similarly to the global proteomics dataset by imputing features with more than 50% quantification in the cohort and converting to log2 space—and Setlur et al.,^48^ was used in the validation of the ERG status differential analysis. A Spearman rank correlation was calculated for log2FC between co-significant (adj. p < 0.05) features between datasets. Sparse Partial Least Squares Discriminant Analysis (sPLS-DA; mixOmics v6.20.0; caret v6.0.93) was used to assess classification performance of ERG status for 23 co-significant features^159–161^. Model parameters were first optimized on a z-score normalized 70%:30% split of the global proteomics dataset prior to training with the full dataset and performance assessed in the z-score normalized validation datasets. An average AUROC was calculated for the global proteomics training dataset as an average Mahalanobis distance to the predicted centroids over 1000 iterations on 2 components (seed 189). Prediction of performance for the validation datasets was performed with pROC (v1.18.0)^162^.

### Identification of candidate driver genes using connectivity map analysis

To identify candidate genes driving cellular responses to copy number alterations, we employed the Connectivity Map (CMAP) analysis data set (GSE92742)^38,39^, adapting the method of Krug. et al.^163^. Specifically, we first identified 1943 genes exhibiting CNA events in at least 12 samples and were associated with 20 or more significantly *trans*-correlated proteins (Pearson correlation |ρ| > 0.3). For each *trans*-correlated gene-protein pair, we defined “up” or “down” protein expression signatures by comparing protein levels in CN-altered samples to those with copy number neutral samples. Genes were classified as deleted or amplified based on a frequency difference of at least five samples in their respective frequencies. We then queried these gene signatures against the Library of Integrated Network-based Cellular Signatures (LINCS) shRNA knockdown database, evaluating the connectivity scores and enrichment p values to select candidate driver genes. We assessed the proportion of *trans*-correlated genes among 407 candidates and determined how many could be directly explained by gene expression changes. Candidate driver genes were ultimately identified based on normalized connectivity scores (NCS) meeting two criteria: (1) an outlier NCS (beyond 1.5 IQR), enriched for cis-located genes (Fisher’s exact test, BH-FDR corrected p < 0.1); and (2) a significant positive cis-correlation. This approach yielded eight candidate driver genes.

To confirm that our identified candidate driver genes were not the result of chance, we conducted a permutation test^37^. For our analysis involving 407 queries, we replaced the actual *trans*-genes with randomly selected genes and repeated the CMAP enrichment analysis. To estimate the false discovery rate (FDR), each of our 10 permutation runs was treated as a Poisson sample with rate λ, representing the count of identified candidate driver genes. Due to the small number of permutations (n=10) and λ, we calculated a Score confidence interval^164^, using its midpoint to estimate the expected number of false positives. Based on these 10 random permutations, the overall FDR was determined to be 0.25, with a 95% confidence interval of (0.1423, 0.3557).

### Identification of ancestry-associated expression quantitative trait loci

Ancestry-associated expression quantitative trait loci (eQTL) were identified as previously described^41^. This process integrates protein abundance, genetic variation, and population allele frequency data to find candidate eQTLs with distinct allele frequency differences between populations. Briefly, proteins differentially abundant between self-identified Black and White patient prostate adenocarcinoma (PRAD) tumors (LIMMA adjusted p < 0.05) were mapped to eQTLs from the PancanQTL^40^. Population allele frequencies for European and African superpopulations were obtained from the gnomAD (v4.1)^165,166^ and 1000 Genomes Project (1KGP; vPhase3)^27^. Allele frequencies (AF) were extracted for male-only populations eQTLs from gnomAD (African: AF_afr_XY; European: AC_fin_XY, AC_nfe_XY, AN_fin_XY, AN_nfe_XY) and 1KGP (African: AFR_AF; European: EUR_AF). Germline variant call files (VCFs) were processed and filtered using bcftools (v1.17)^167^ to extract genotype and variant details (chromosome, position, ref allele, alt allele and dbSNP ID) for eQTLs of interest. eQTLs showing ≥ 40% AF difference between African and European populations and between the study cohort’s allele frequencies, calculated from available germline-level genotype data, were prioritized as ancestry-associated candidates. Candidates were further filtered to those with significant altered protein abundances between genotype groups (Mann-Whitney U test p < 0.05) and concordant median fold change directionality with PanCanQTL expression beta values. Transcript-level data for significantly altered protein candidates mapping to putative ancestry-associated eQTL events were further evaluated in in TCGA PRAD tumors (White: n = 413; Black, n = 57; FDR < 0.1)^20^, downloaded from LinkedOmics. This workflow was performed in R v.4.3.1 using the following packages: vcfR (v.1.15.0)^168^, limma (v3.60.6)^169^, ggplot2 (v.3.5.2)^170^, and patchwork (v.1.3.0). Code is available at: https://github.com/GYNCOE/APOLLO3.

### Role in cancer and targetability annotation

To annotate genes with a role in cancer or a driver function, we consolidated information from several databases: the COSMIC Cancer Gene Census (release v100, May 21, 2025)^171,172^, the OncoKB Cancer Gene List (03/28/2025 update)^173,174^, and recent pan-cancer studies^175,176^. We defined four unified clinical actionability levels, by curating data from OncoKB, COSMIC, and CanSar.ai^177^. For each gene, we selected the highest actionability level, prioritizing evidence from prostate cancer and other solid tumors over pan-cancer annotations. From OncoKB, we selected the highest clinical evidence level, excluding annotations for resistance in blood cancers. For the COSMIC actionability dataset (Version 15, Release Date: February 27, 2025), we chose the highest available rank for each gene, selecting one representative entry if multiple shared the top rank. Finally, we assigned each gene one of four CanSAR targetability levels: Level 1 (directly druggable; ≥50% similarity), Level 2 (near a druggable target), Level 3 (active screened compounds), and Level 4 (structurally druggable).

Clinical actionability Level 1 include alterations supported by strong clinical evidence, including regulatory approvals (e.g., FDA) or professional guidelines for the use of approved drugs in the same cancer type It combines OncoKB Levels 1–2 and COSMIC Ranks 1–2. Level 2 includes alterations under active clinical investigation or drug repurposing efforts, based on late-phase clinical trial data or established drug targets without direct disease-specific validation. Sources for this level are OncoKB Level 3, COSMIC Rank 3, and CanSAR Level 1. Level 3 comprises targets with emerging preclinical or clinical support, such as case reports or early-phase investigational data but without established clinical standards of care. Sources include OncoKB Level 4, COSMIC Rank 4, and CanSAR Level 2. Level 4 represents early-stage drug discovery opportunities that include genes identified through preclinical compound screening or structural predictions, without clinical validation This level corresponds to **CanSar.ai** Levels 3–4.

### Gene effect and dependency levels and therapeutic vulnerability annotations

Median gene effect scores (Chronos) and gene dependency probabilities from DepMap CRISPR-Cas9 screens (24Q4 release)^178^ of nine prostate adenocarcinoma cell lines were each classified into four levels of functional essentiality. Gene effect scores were grouped as: Level 1 (< –0.7), Level 2 (–0.7 to –0.5), Level 3 (–0.5 to 0), and Level 4 (> 0). Dependency probabilities were grouped as: Level 1 (≥ 0.7), Level 2 (0.6–0.7), Level 3 (0.5–0.6), and Level 4 (< 0.5). These levels were interpreted as: strongly essential (Level 1), moderately essential (Level 2), low dependency (Level 3), and no dependency (Level 4), reflecting decreasing impact of gene knockout on cell viability. To integrate the two metrics, each gene was assigned the more severe (lower) of the two scores, ensuring that strong evidence from either source would define the final essentiality classification.

Genes were classified into four therapeutic vulnerability tiers by integrating prostate cancer-specific functional essentiality scores and unified clinical actionability information. This combined framework prioritized genes based on both their biological importance to tumor cell viability and the feasibility of therapeutic targeting. Tier 1 (Essential and actionable) included genes with high tractability (Levels 1– 2) and moderate to strong essentiality (Scores 1–3). Tier 2 (Essential – No direct drug) comprised genes with limited or unknown tractability (Level 3 – 4 or missing) but high essentiality (Scores 1 – 2). Tier 3 (Low dependency and limited tractability) included genes with low essentiality (Score 3) and limited tractability. Remaining genes were assigned to Tier 4 (No dependency/Unclassified). Each tier was also assigned a numeric score to facilitate downstream prioritization and visualization.

### Multi-omics gene set analysis

Multi-Omics Gene Set Analysis (MOGSA) was conducted using the MOGSA R package (v1.380)^50^. Single-sample gene-set scores (GSS) were computed for curated pathway collections, including KEGG, WikiPathways, Gene Ontology (GO), MSigDB Hallmark, and C2 gene sets, across omics layers comprising differential copy number alterations (n = 1,631 genes; identified by Fisher’s Exact Test and elastic net models), age- and PSA-adjusted differentially mutated SNVs (n = 10), and differentially abundant global proteomic (n = 313) and phosphoproteomic substrate (n = 63) features. To identify pathways enriched in predefined sample groups, we applied a two-step filtering strategy: first, we retained pathways comprising 5 to 500 genes that showed per-sample GSS p< 0.01 in at least 50% of samples; second, we applied generalized linear models (GLMs) to the GSS matrix and selected pathways with a false discovery rate p < 0.01.

#### Integrative clustering using iClusterBayes

Integrative clustering was performed using the iClusterBayes function from the iClusterPlus R package (v1.42) to jointly model tumor profiles across four molecular layers: somatic SNVs, CNAs, global proteome, and phosphoproteome data^51^. This Bayesian multi-omics clustering approach, which identifies molecular subtypes based on shared latent variation across data layers, provides significant improvement over earlier genomics-only^20,23^ or proteomics/phosphoproteomics-only^76^ classification of molecular subtypes. SNVs were encoded as binary variables and filtered to retain mutations with a minimum frequency of 2%. To reduce the dimensionality of Battenberg-derived copy number segmentation data, we applied *CNregions* function using epsilon = 0.005 to control segment merging. A fixed threshold was used, known germline copy number variants were excluded, and at least 50% overlap with the CNV reference was required. This yielded a reduced matrix of ∼800 consensus regions across all samples for downstream analysis. Gene-level log2 copy number ratios were subsequently computed by mapping segmented regions to gene coordinates. CNA features with low variance (<0.01) were excluded. All continuous datasets (CNA, proteome, and phosphoproteome) were scaled to unit variance and modeled using Gaussian assumptions. Model fitting was conducted across a range of latent dimensionalities (K = 2 to 6). Regularization parameters were specified as prior.gamma values of 0.1 (SNV), 0.25 (CNA), 0.01 (proteome), and 0.2 (phosphoproteome). Posterior inference was carried out using 25,000 burn-in iterations, followed by 20,000 draws with thinning every 5 steps. The optimal model was selected based on the lowest Bayesian Information Criterion (BIC). Although the selected model was defined with K = 2 latent variables, three distinct sample clusters were identified by applying Gaussian mixture modeling to the posterior latent variable matrix (meanZ). Clustering results were visualized using heatmaps of posterior gamma scores for features retained across omics layers, with samples ordered by their cluster assignments.

To assess generalizability, the analysis was repeated using the TCGA prostate cancer cohort filtered for patients of African and European ancestry (393 European ancestry; 58 African Ancestry) and, substituting mRNA expression data for the proteome layer. SNV and CNA data were processed as described above, including dimensionality reduction of segmented CNA data using the CNregions function with epsilon = 0.003, resulting in approximately 800 consensus regions. mRNA features were filtered to retain the top 8,000 genes with variance ≥0.05. The model was fit using prior.gamma values of 0.1 (SNV), 0.25 (CNA), and 0.001 (mRNA), with K ranging from 2 to 5. Additionally, a sensitivity analysis was conducted by excluding the phosphoproteome layer from the original four-layer dataset and re-running the integrative clustering using the same preprocessing steps and tuning parameters.

### Survival analysis

Cox proportional hazards regression models were applied to evaluate the associations between disease progression and gene-level alterations, including CNVs, protein and phosphoprotein expression levels. Protein and phosphoprotein expression levels were analyzed both as continuous and as categorical variables, with cutoff points determined using the Youden Index. Univariable analyses were first conducted to identify candidate predictors. We observed that age and PSA values at diagnosis, and patient’s percent African ancestry were significant predictors, and these were included in the multivariable Cox models to estimate hazard ratios (HRs) and 95% confidence intervals (CIs). Survival probabilities were calculated using the Kaplan–Meier (KM) method, defining time-to-event was as the interval from radical prostatectomy to either BCR or metastasis. For KM analyses using the TCGA dataset, progression-free interval (PFI) was used as the primary outcome endpoint^179^. Patients who did not experience an event during the study period were censored at the date of their last follow-up. Kaplan–Meier survival curves were generated for groups defined by gene-level alterations, as well as by protein and phosphoprotein expression levels. Differences between survival curves were assessed using the log-rank test. Receiver operating characteristic (ROC) curve analysis was performed to compare the discriminatory performance of predictive models. The areas under the curve (AUCs) were calculated for each model. All statistical tests were two-sided, and p values ≤ 0.05 were considered significant unless otherwise specified. Analyses were performed using SAS software (v9.4, SAS Institute, North Carolina, USA)

## Acknowledgement

We acknowledge all patients who participated in the study. We also thank Kerri Cronin, Jaime Boone, Susan Kwon, and Christopher Jamieson for their program support. Funding for this study was provided in part by the Uniformed Services University of the Health Sciences (USUHS) through the Henry M. Jackson Foundation for the Advancement of Military Medicine, Inc.

## Disclaimer

The contents of this publication are the sole responsibility of the author(s) and do not necessarily reflect the views, opinions or policies of Uniformed Services University of the Health Sciences (USUHS), the Henry M. Jackson Foundation for the Advancement of Military Medicine, Inc., the Department of Defense (DoD) or the Departments of the Army, Navy, or Air Force. Mention of trade names, commercial products, or organizations does not imply endorsement by the U.S. Government.

## APOLLO Research Network

Jordan A. Driscoll, Satishkumar Ranganathan Ganakammal, George L. Maxwell, Xiaoying Lin, Hai Hu, Sean Hanlon

## Declaration of generative AI and AI-assisted technologies in the writing process

During the preparation of this work, the authors used a large language model for grammar and language refinement in the writing process. After using this tool, the authors reviewed and edited the content as needed and take full responsibility for the final publication.

