## SupplementaryFigures for "Integrated proteogenomics uncovers ancestry-specific and shared molecular drivers in localized prostate cancer"

**Schafer et. al.**

#### SUPPLEMENTARY FIGURES

**Figure S1**

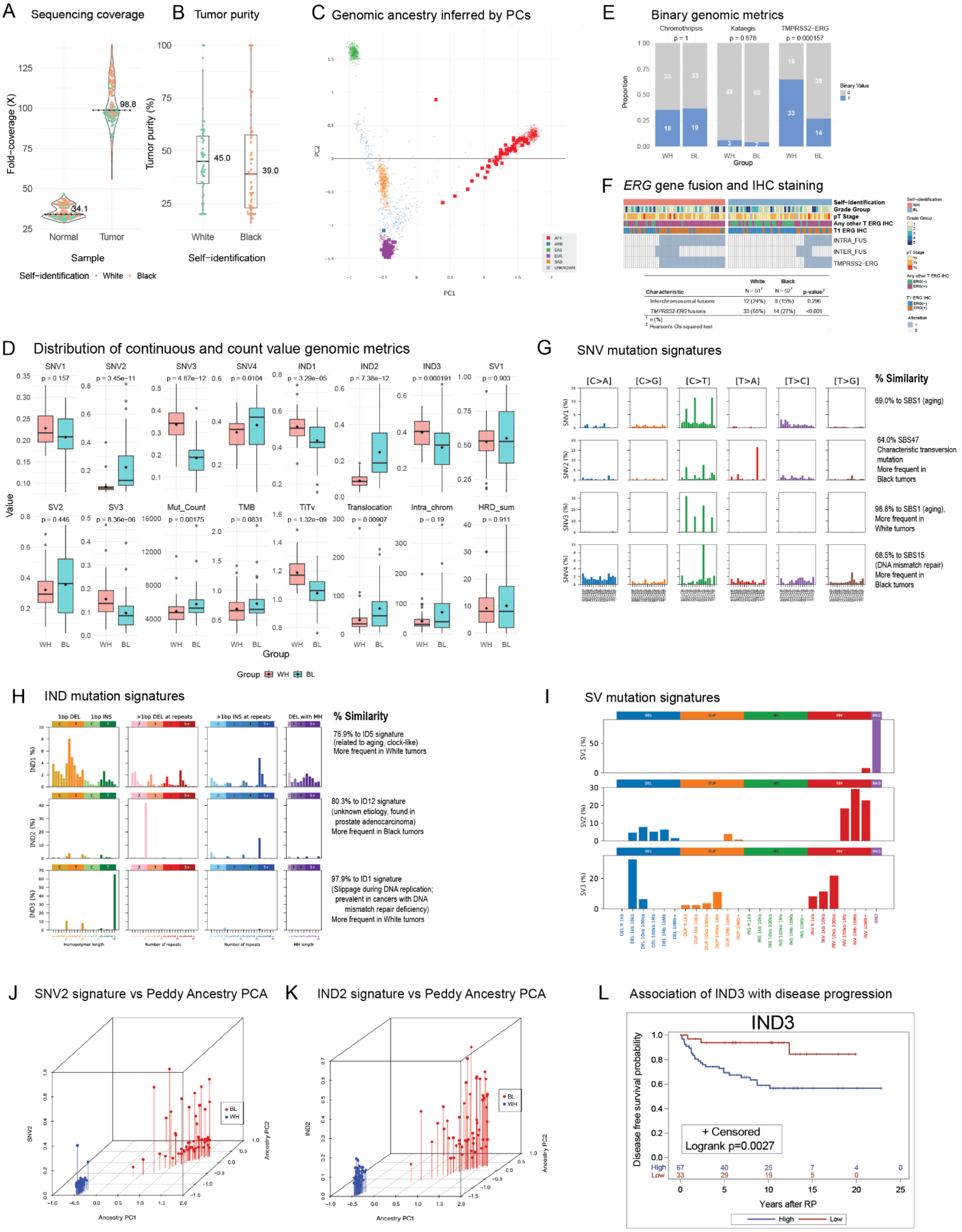

A Sequencing coverage
B Tumor purity
C Genomic ancestry inferred by PCs
E Binary genomic metrics
F ERG gene fusion and IHC staining
D Distribution of continuous and count value genomic metrics
G SNV mutation signatures
H IND mutation signatures
I SV mutation signatures
J SNV2 signature vs Peddy Ancestry PCA
K IND2 signature vs Peddy Ancestry PCA
L Association of IND3 with disease progression
Fold-coverage (X)
Normal
Tumor
Sample
Self-identification White Black
Tumor purity (%)
45.0
39.0
98.8
34.1
Self-identification
PC1
PC2
Chromothripsis p = 1
Kataegis p = 0.678
TMPRSS2-ERG p = 0.000157
Proportion
Group
Binary Value
Self-identification
Grade Group
pT Stage
Any other T ERG IHC
T1 ERG IHC
INTRA_FUS
INTER_FUS
TMPRSS2-ERG
Characteristic
White N = 51
Black N = 52
p-value
Interchromosomal fusions 12 (24%) 8 (15%) 0.296
TMPRSS2-ERG fusions 33 (65%) 14 (27%) <0.001
n (%)
Pearson's Chi-squared test
SNV1 p = 0.157
SNV2 p = 3.45e-11
SNV3 p = 4.87e-12
SNV4 p = 0.0104
IND1 p = 3.29e-05
IND2 p = 7.38e-12
IND3 p = 0.000191
SV1 p = 0.903
SV2 p = 0.446
SV3 p = 8.36e-06
Mut_Count p = 0.00175
TMB p = 0.0831
TiTv p = 1.32e-09
Translocation p = 0.00907
Intra_chrom p = 0.19
HRD_sum p = 0.911
Value
Group WH BL
[C>A] [C>G] [C>T] [T>A] [T>C] [T>G]
% Similarity
69.0% to SBS1 (aging)
64.0% SBS47 Characteristic transversion mutation More frequent in Black tumors
98.8% to SBS1 (aging), More frequent in White tumors
68.5% to SBS15 (DNA mismatch repair) More frequent in Black tumors
1bp DEL 1bp INS >1bp DEL at repeats >1bp INS at repeats DEL with MH
76.9% to ID5 signature (related to aging, clock-like) More frequent in White tumors
80.3% to ID12 signature (unknown etiology, found in prostate adenocarcinoma) More frequent in Black tumors
97.9% to ID1 signature (Slippage during DNA replication; prevalent in cancers with DNA mismatch repair deficiency) More frequent in White tumors
Homopolymer length
Number of repeats
MH length
DEL DUP INS INV BND
SNV2
IND2
Ancestry PC1
Ancestry PC2
IND3
Disease free survival probability
+ Censored
Logrank p=0.0027
High 67 40 25 7 4 0
Low 33 29 19 5 0
Years after RP
High Low

**Figure S1**. **Genomic landscape and patient demographics in a cancer cohort** **(A)** Whole genome sequencing of DNA from normal and tumor samples achieved median coverage depths of 34.1 X and 98.8X, respectively. (**B**) The median tumor purity of tumors from White patients and Black patients were 45% and 39%, respectively. (**C**) Patient ancestry was classified into five groups based on self-identified and *Peddy* predicted ancestry matching ancestry groups from the Human Genome Diversity Project and the 1000 Genomes Project: Africa (AFR), the Americas (AMR), East Asia (EAS), Europe (EUR) and South Asia (SAS). The AFR super population includes the African Ancestry in SW USA (ASW) population. (**D**) Comparison of genomic metrics with continuous and count values between Black and White patients, with assessed Wilcoxon rank-sum test p-values. (**E**) Comparison of binary genomic values (e.g., presence or absence) between Black and White patients with significance assessed by Fisher's exact test p-values. (**F**) Cases with intra-chromosomal (*TMPRSS2-ERG*) and inter-chromosomal *ERG* fusions, annotated by their corresponding ERG IHC staining status. Somatic mutational signatures for single-base substitutions (SNV; **G**), insertion–deletions (IND; **H**), and structural variants (SV; **I**). For (**G**) and (**H**), the closest-matching COSMIC reference signatures are indicated (v3.2, March 2021). (**J-K**) Principal-component analysis (PCA) of tumor-level exposures to SNV2 (**J**) and IND2 (**K**) show these signatures trend with greater variability in germline genome among Black patients. (**L**) Univariate Kaplan-Meier survival curves showing time to biochemical recurrence **(**BCR) and/or metastasis for IND3.

### Figure S2

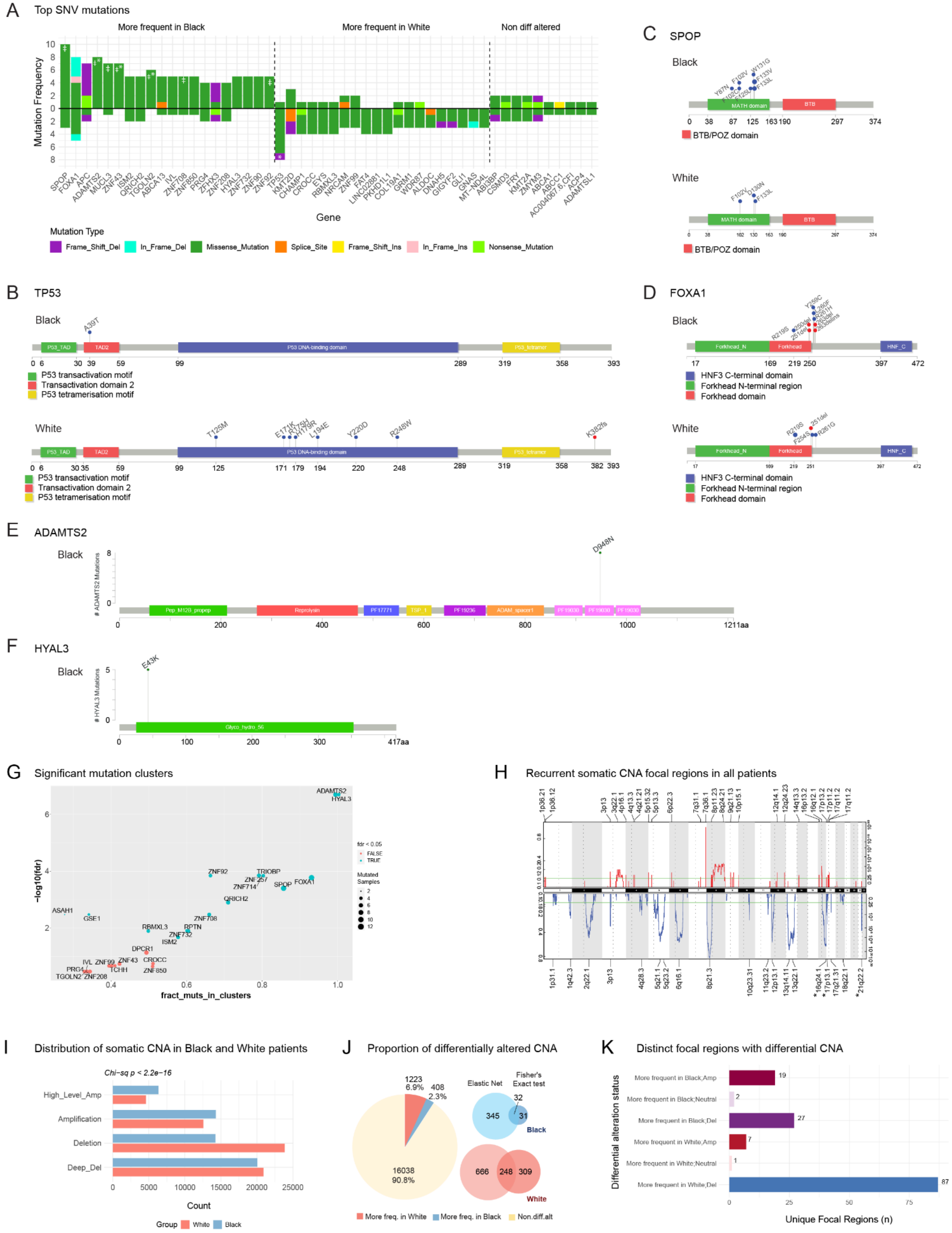

A
Top SNV mutations
More frequent in Black
More frequent in White
Non diff altered
Mutation Frequency
Gene
Mutation Type
Frame_Shift_Del
In_Frame_Del
Missense_Mutation
Splice_Site
Frame_Shift_Ins
In_Frame_Ins
Nonsense_Mutation
B
TP53
Black
White
P53 transactivation motif
Transactivation domain 2
P53 tetramerisation motif
C
SPOP
Black
White
MATH domain
BTB
BTB/POZ domain
D
FOXA1
Black
White
HNF3 C-terminal domain
Forkhead N-terminal region
Forkhead domain
E
ADAMTS2
Black
F
HYAL3
Black
G
Significant mutation clusters
-log10(fdr)
fract_muts_in_clusters
H
Recurrent somatic CNA focal regions in all patients
I
Distribution of somatic CNA in Black and White patients
Chi-sq p < 2.2e-16
High_Level_Amp
Amplification
Deletion
Deep_Del
Count
Group
White
Black
J
Proportion of differentially altered CNA
Elastic Net
Fisher's Exact test
More freq. in White
More freq. in Black
Non.diff.alt
K
Distinct focal regions with differential CNA
Differential alteration status
More frequent in Black;Amp
More frequent in Black;Neutral
More frequent in Black;Del
More frequent in White;Amp
More frequent in White;Neutral
More frequent in White;Del
Unique Focal Regions (n)

**Figure S2. Somatic SNV mutation and copy number alteration profiles.** (**A**) Top differentially mutated genes, ordered by frequency of occurrence. SNVs above the midline represent mutations in Black patients, below the midline, White patients. Asterisk (*) denotes Fisher's exact test $p < 0.05$; diesis (ǂ), Firth's $p < 0.05$ (adjusted for age and PSA). **(B-F)** Lollipop plots showing locations of SNV mutation detected in Black vs White tumors for *TP53* (**B**), *SPOP* (**C**), and *FOXA1* (**D**). Mutational hotspots were in *ADAMTS2* (**E**) and *HYAL3* (**F**) and were detected only in Black tumors. (**G**) Plot of log FDR vs fraction of mutation clusters of recurrent mutations. **(H)** Recurrent somatic copy number altered focal regions in all patients called by GISTIC2 ($q < 0.25$). Top red tracks indicate copy gain. Asterisks denote significantly differentially altered focal regions (Wilcoxon rank-sum $p < 0.05$). (**I**) Distribution of somatic copy number alterations in Black and White patients. (**J**) Differential sCNA identified by Fisher's exact and by Elastic Net, showing distinct and shared calls. (**K**) Distribution of unique focal regions harboring differentially altered genes between groups.

### Figure S3

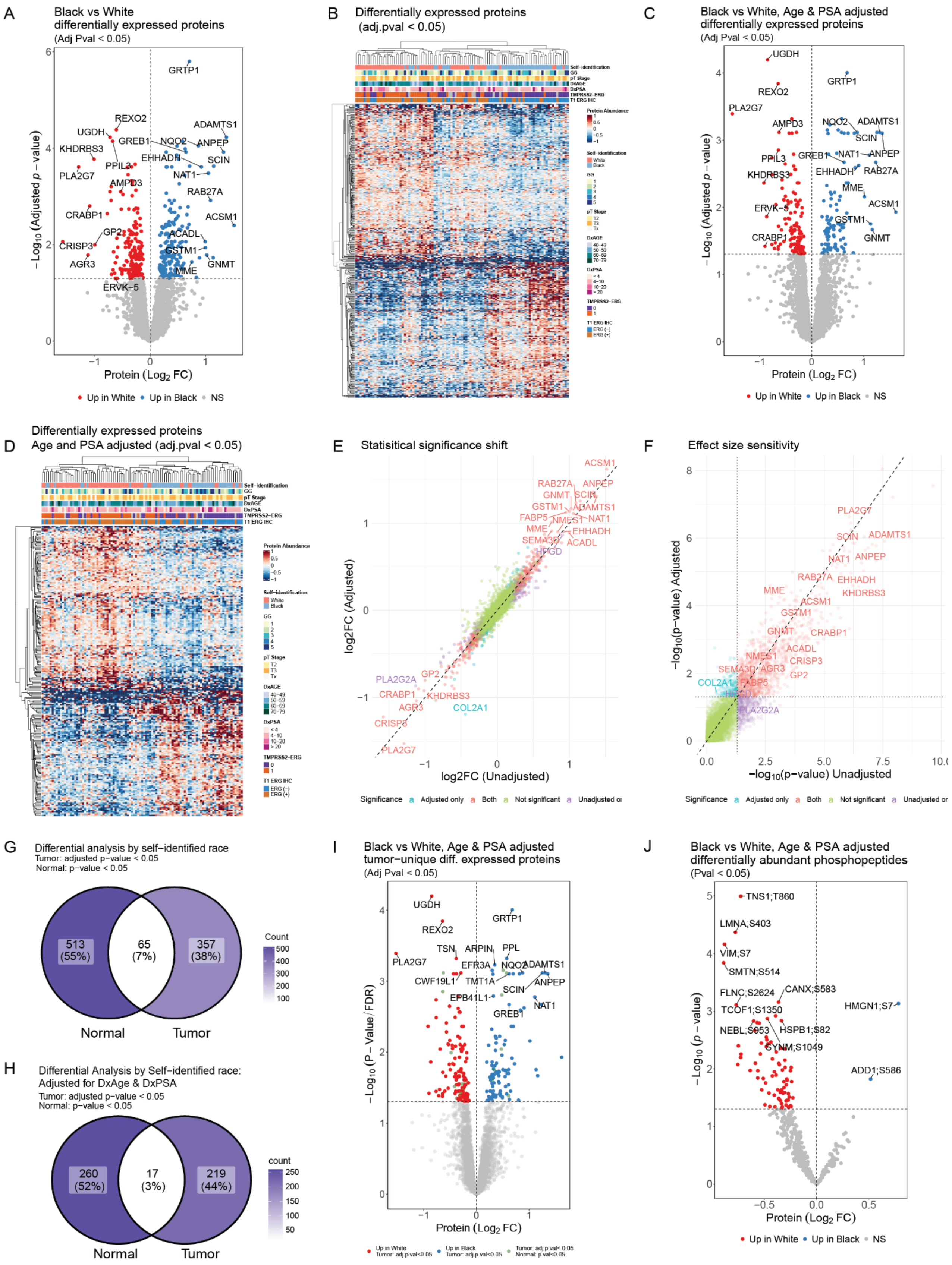

A
Black vs White
differentially expressed proteins
(Adj Pval < 0.05)
GRTP1
REXO2
ADAMTS1
UGDH
GREB1
NQO2
ANPEP
KHDRBS3
EHHADH
SCIN
PPIL3
PLA2G7
NAT1
AMPD3
RAB27A
CRABP1
ACSM1
GP2
ACADL
CRISP3
GSTM1
AGR3
GNMT
MME
ERVK-5
−Log10 (Adjusted p − value)
Protein (Log2 FC)
Up in White
Up in Black
NS
B
Differentially expressed proteins
(adj.pval < 0.05)
Self−identification
GG
pT Stage
DxAGE
DxPSA
TMPRSS2−ERG
T1 ERG IHC
Protein Abundance
C
Black vs White, Age & PSA adjusted
differentially expressed proteins
(Adj Pval < 0.05)
UGDH
GRTP1
REXO2
PLA2G7
AMPD3
NQO2
ADAMTS1
SCIN
PPIL3
GREB1
NAT1
ANPEP
KHDRBS3
EHHADH
RAB27A
MME
ERVK−5
ACSM1
GSTM1
CRABP1
GNMT
−Log10 (Adjusted p − value)
Protein (Log2 FC)
Up in White
Up in Black
NS
D
Differentially expressed proteins
Age and PSA adjusted (adj.pval < 0.05)
Protein Abundance
Self−identification
White
Black
GG
pT Stage
DxAGE
DxPSA
TMPRSS2−ERG
T1 ERG IHC
E
Statisitical significance shift
ACSM1
RAB27A
ANPEP
GNMT
SCIN
GSTM1
ADAMTS1
FABP5
NMES1
NAT1
MME
EHHADH
SEMA3D
ACADL
HPGD
PLA2G2A
GP2
CRABP1
KHDRBS3
AGR3
COL2A1
CRISP3
PLA2G7
log2FC (Adjusted)
log2FC (Unadjusted)
Significance
Adjusted only
Both
Not significant
F
Effect size sensitivity
PLA2G7
SCIN
ADAMTS1
NAT1
ANPEP
RAB27A
EHHADH
MME
KHDRBS3
ACSM1
GSTM1
GNMT
CRABP1
ACADL
NMES1
CRISP3
SEMA3D
AGR3
GP2
COL2A1
FABP5
PLA2G2A
−log10(p−value) Adjusted
−log10(p−value) Unadjusted
Significance
Adjusted only
Both
Not significant
G
Differential analysis by self-identified race
Tumor: adjusted p-value < 0.05
Normal: p-value < 0.05
513
(55%)
65
(7%)
357
(38%)
Count
Normal
Tumor
H
Differential Analysis by Self−identified race:
Adjusted for DxAge & DxPSA
Tumor: adjusted p-value < 0.05
Normal: p-value < 0.05
260
(52%)
17
(3%)
219
(44%)
count
Normal
Tumor
I
Black vs White, Age & PSA adjusted
tumor−unique diff. expressed proteins
(Adj Pval < 0.05)
UGDH
GRTP1
REXO2
TSN
ARPIN
PPL
PLA2G7
EFR3A
NQO2
ADAMTS1
CWF19L1
TMT1A
SCIN
ANPEP
EPB41L1
NAT1
GREB1
−Log10 (P − Value/FDR)
Protein (Log2 FC)
Up in White
Tumor: adj.p.val<0.05
Up in Black
Tumor: adj.p.val<0.05
Tumor: adj.p.val< 0.05
Normal: p.val<0.05
J
Black vs White, Age & PSA adjusted
differentially abundant phosphopeptides
(Pval < 0.05)
TNS1;T860
LMNA;S403
VIM;S7
SMTN;S514
FLNC;S2624
CANX;S583
TCOF1;S1350
HMGN1;S7
NEBL;S953
HSPB1;S82
SYNM;S1049
ADD1;S586
−Log10 (p − value)
Protein (Log2 FC)
Up in White
Up in Black
NS

**Figure S3. Effect of covariate adjustment (age and PSA at diagnosis) on global, tumor-unique proteome, and phospho-proteomes.** All group comparisons are between Black and White patient tumors; unless noted, significance is adj. $p < 0.05$. **Volcano plots** for the unadjusted (**A**) and age and PSA adjusted (**C**) global proteome. Significantly differentially abundant proteins are highlighted. Red and blue denote proteins more abundant in White and Black tumors, respectively. **Heatmaps** show the hierarchical clustering of proteins significant in the unadjusted (**B**) and adjusted models (**D**). **Model concordance:** (**E**) Scatter plot comparing $\log_2$ fold-changes ($\log_2$FC) from unadjusted (x-axis) vs. adjusted (y-axis) analyses. Points near the diagonal indicate stable effects sizes; color encodes significance status across models. (**F**) Scatter plot of -log10 (p-values) from unadjusted (x-axis) and adjusted (y-axis) models. Dotted lines mark the nominal $p = 0.05$ threshold. Points above both lines are consistently significant; shifts across quadrants indicate sensitivity to covariate adjustment. **Tumor vs normal (global proteome):**Venn diagram summarizing the global tumor- versus normal-proteome ($p < 0.05$) comparison. Counts indicate proteins uniquely altered in tumors: **357** in the unadjusted analysis (**G**) and **219** after covariate adjustment (**H**). **Tumor-unique proteome:** (**I**) Volcano plot for proteins detected only in tumor samples, after age and PSA adjustment. Red and blue denote proteins more abundant in White and Black tumors, respectively. Significant proteins are labeled. **Phosphoproteome:** (**J**) Volcano plot after age and PSA adjustment; red and blue denote phosphosites more abundant in White and Black tumors, respectively ($p < 0.05$).

**Figure S4**

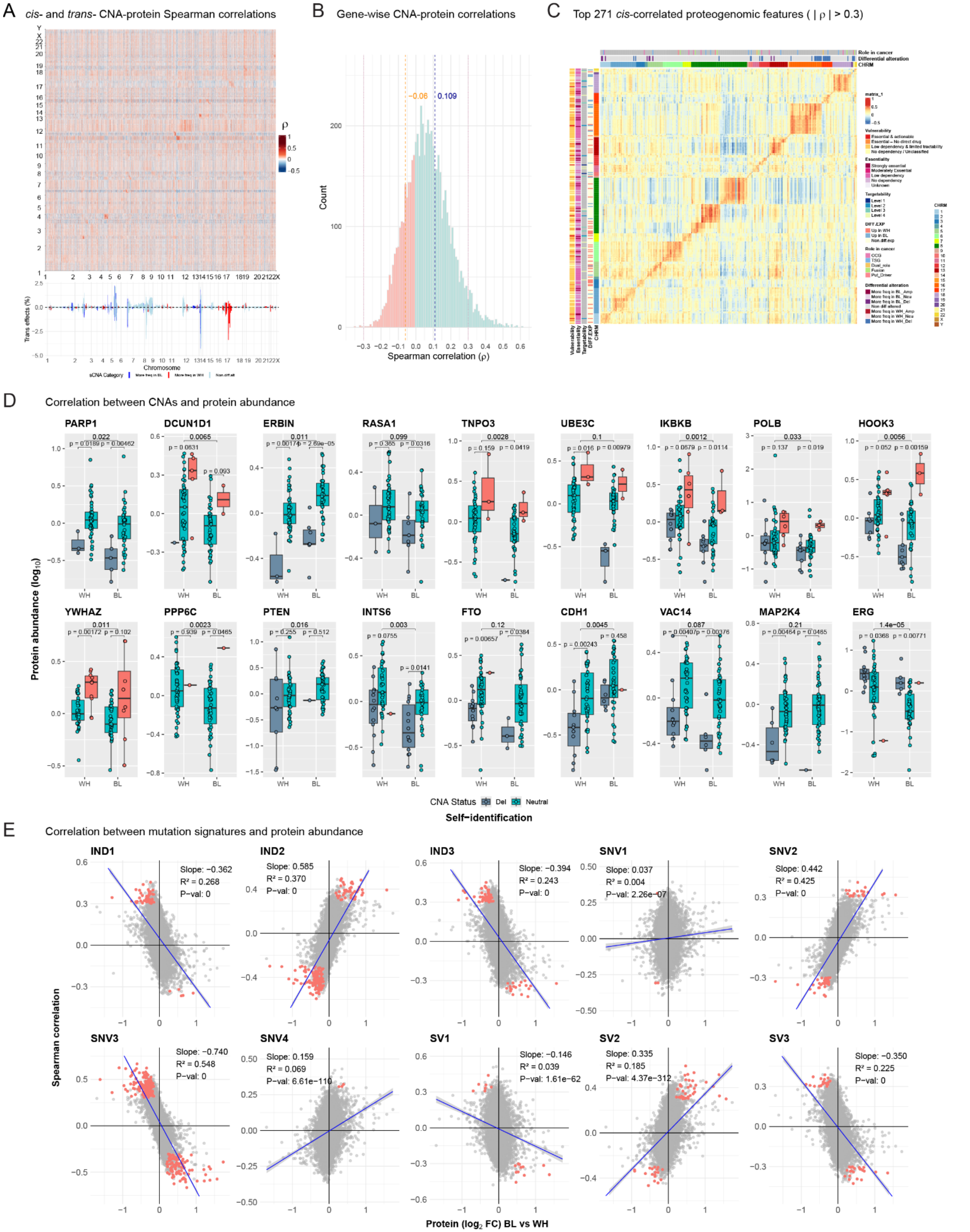
A cis- and trans- CNA-protein Spearman correlations
B Gene-wise CNA-protein correlations
C Top 271 cis-correlated proteogenomic features ( | ρ | > 0.3)
D Correlation between CNAs and protein abundance
E Correlation between mutation signatures and protein abundance
Spearman correlation
Count
Chromosome
Trans effects (%)
Protein abundance (log10)
CNA Status Del Neutral
Self−identification
Protein (log2 FC) BL vs WH
PARP1
DCUN1D1
ERBIN
RASA1
TNPO3
UBE3C
IKBKB
POLB
HOOK3
YWHAZ
PPP6C
PTEN
INTS6
FTO
CDH1
VAC14
MAP2K4
ERG
IND1
IND2
IND3
SNV1
SNV2
SNV3
SNV4
SV1
SV2
SV3

**Figure S4. Proteogenomic correlates of Copy Number Alterations (A) Global and gene-specific CNA-protein correlations.** Spearman correlations between CNA (x-axis) -protein abundance (y axis) for all genes. *Cis*-effects appear as a prominent diagonal line, while *trans*-effects form vertical patterns. The histogram shows the fraction of significant *trans*-effects for each CNA gene. (**B**) **Overall distribution of all gene-wise CNA-protein correlations.** The blue, and orange dashed lines, mark the median positive ($\rho = 0.109$) and negative ($\rho = -0.06$) correlations, respectively. (**C**) Heatmap displaying the significant *cis*-correlations (Spearman's $|\rho|>0.3$, adj. $p < 0.05$) for the top 271 proteogenomic features. The x-axis represents CNA; the y-axis, the matched proteins. Column annotations indicate the genes' differential alteration status and their known functional roles in cancer. Row annotations describe protein differential expression, druggability, functional essentiality (from DepMap), and vulnerability (from tractability and essentiality). **(D) Protein abundance changes with CNA status.** Boxplots illustrating how protein abundance changes with different CNA statuses (deletions, neutral, and amplifications) for select genes. Protein abundance is $\log_{10}$ transformed. Genes are arranged by their chromosomal location. Statistical significance is annotated using asterisks: a Wilcoxon test was used for two-level comparisons, while a Kruskal-Wallis test was used for three-level comparisons. Asterisks denote statistical significance: **** ($p < 0.0001$), *** ($p < 0.001$), ** ($p < 0.01$), * ($p < 0.05$), ns ($p > 0.05$). **(E) Correlation between mutation signatures and protein abundance**. Scatter plots show linear regression analysis of Spearman's coefficient ($\rho$) against $\log_2$ fold change ($Log_2FC$) of proteins that are differentially abundant between Black and White patient tumors. Proteins with a positive regression slope and a log2FC>0 are upregulated in tumors from Black patients; proteins with a negative slope are upregulated in tumors from White patients. The coefficient of determination ($R^2$) indicates the proportion of variance in Spearman $\rho$ explained by $\log_2$FC. Colored points highlight proteins that are both significantly differentially expressed (adj. $p < 0.05$) and have a significant Spearman correlation ($| \rho | > 0.3$).

**Figure S5**

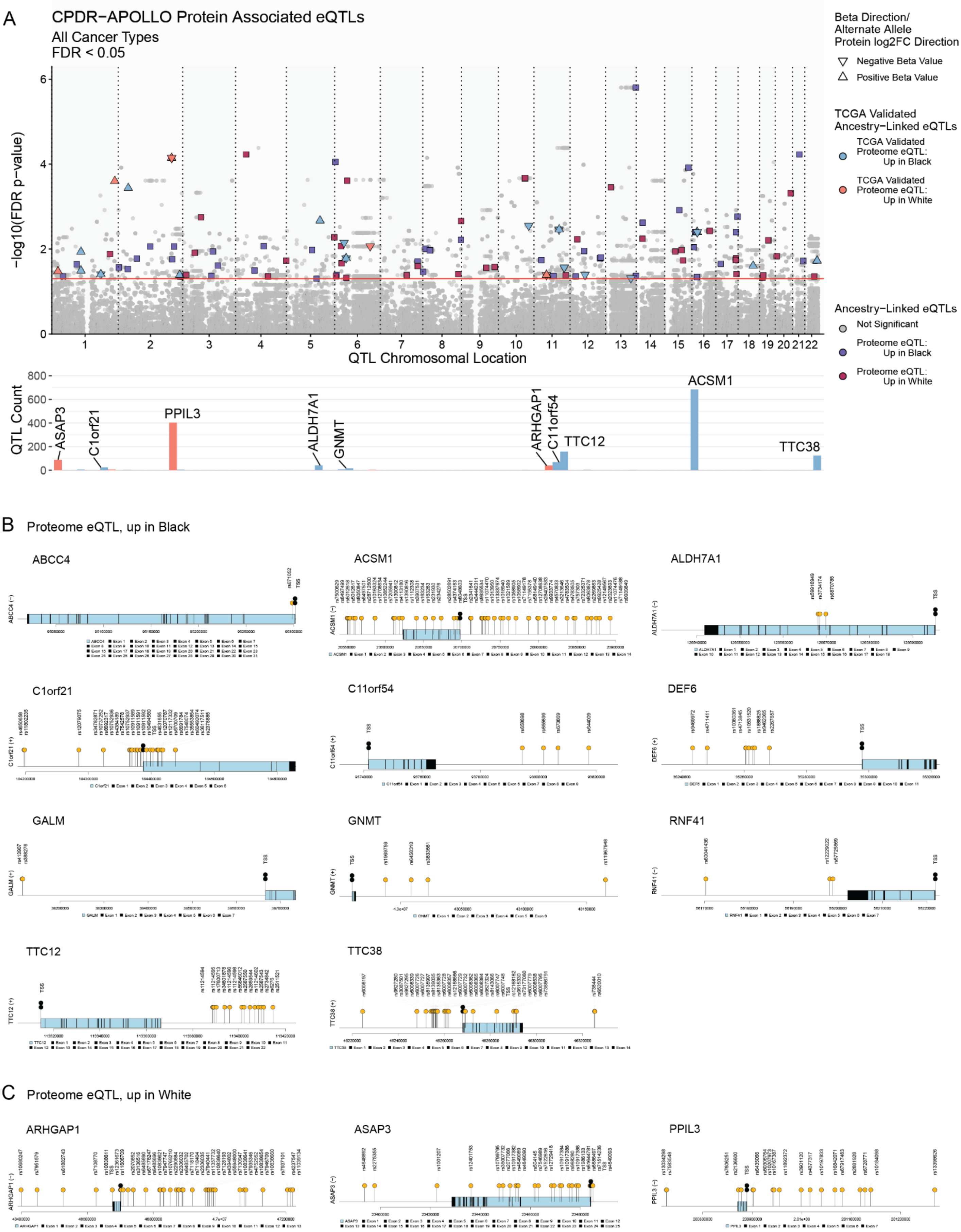


**Figure S5. Ancestry-associated genetic variants correlate with differences in protein abundance.**

**(A) Prioritized candidate ancestry-associated eQTL associated with proteins differentially abundant between Black and White patients.** Top: Plot details eQTL events (x-axis) and their associated proteins. The eQTLs were prioritized from the Pan-cancer quantitative trait loci (PancanQTL) resource and linked to proteins with proteins with significantly altered abundance (adj. $p < 0.05$, y-axis) in tumors from Black and White patients. Ancestry-associated eQTL candidates were defined by an allele frequency difference of at least 40% between African and European superpopulations from gnomAD, 1KGP, and the CPDR-APOLLO cohort. Further prioritization required a concordant expression direction between eQTL beta values and protein abundance, comparing heterozygous alternate and homozygous reference genotype groups (Mann-Whitney U test $p < 0.05$). Proteins associated with ancestry-associated eQTLs in Black patients are indicated by purple squares; those in White patients, by maroon squares. Protein-level findings were compared to transcript data from self-described Black (n=57) and White (n=413) TCGA prostate cancer patients. Candidates with significant alterations (adj. $p < 0.1$) and concordant fold-change differences at the transcript level are annotated as blue triangles for proteins elevated in Black patients and as coral triangles for proteins elevated in White patients. The direction of the triangle corresponds to the concordant beta value or protein abundance of the alternate allele expression. Bottom: Bar plot shows the number of eQTLs associated with TCGA-validated genes. Labeled genes have a minimum of ten associated eQTLs. (**B**) and (**C**) **eQTLs associated with proteins significantly elevated in Black patient tumors and in White patient tumors, respectively.**

### Figure S6

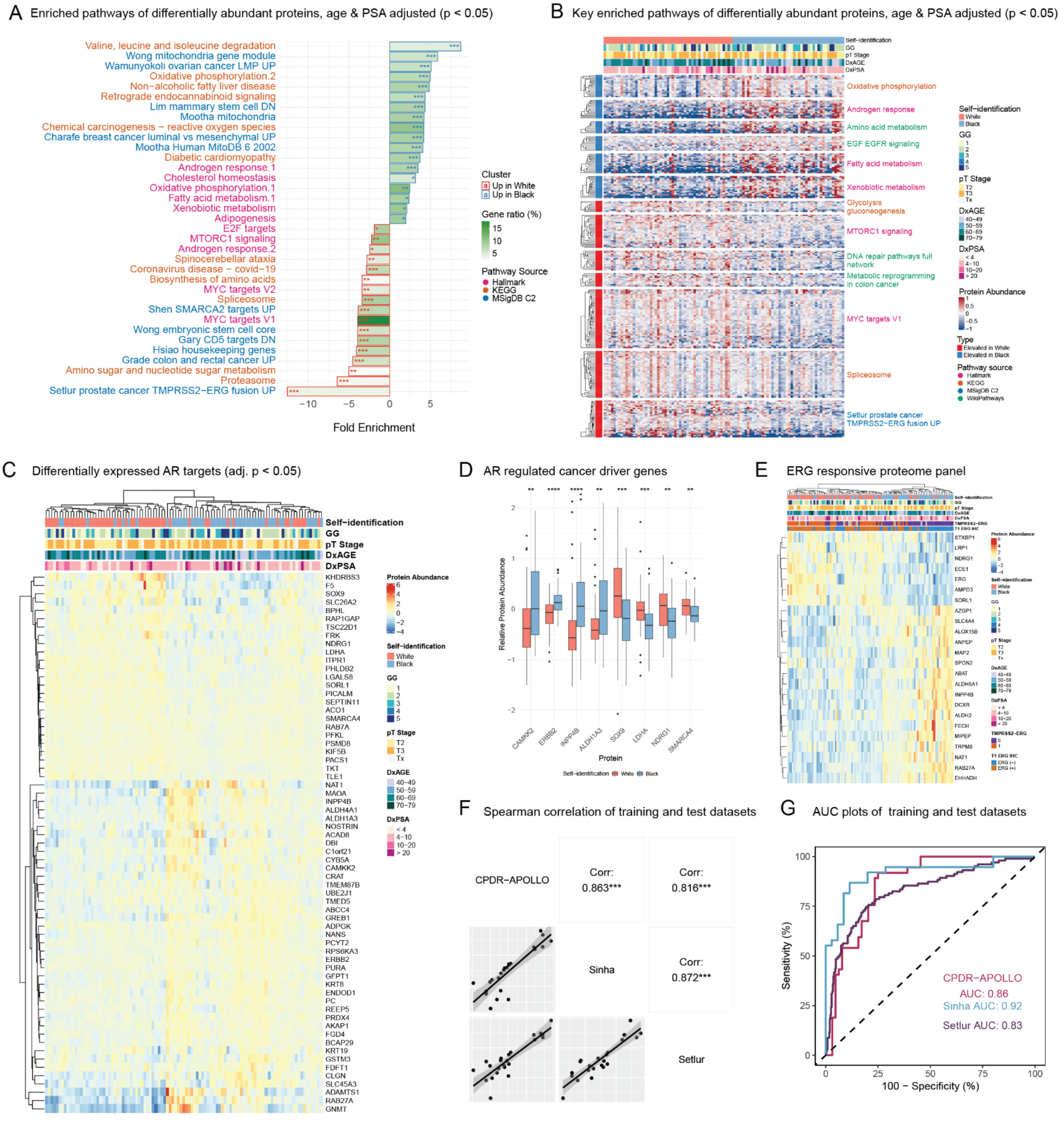

A Enriched pathways of differentially abundant proteins, age & PSA adjusted (p < 0.05)
Valine, leucine and isoleucine degradation
Wong mitochondria gene module
Wamunyokoli ovarian cancer LMP UP
Oxidative phosphorylation.2
Non−alcoholic fatty liver disease
Retrograde endocannabinoid signaling
Lim mammary stem cell DN
Mootha mitochondria
Chemical carcinogenesis − reactive oxygen species
Charafe breast cancer luminal vs mesenchymal UP
Mootha Human MitoDB 6 2002
Diabetic cardiomyopathy
Androgen response.1
Cholesterol homeostasis
Oxidative phosphorylation.1
Fatty acid metabolism.1
Xenobiotic metabolism
Adipogenesis
E2F targets
MTORC1 signaling
Androgen response.2
Spinocerebellar ataxia
Coronavirus disease − covid−19
Biosynthesis of amino acids
MYC targets V2
Spliceosome
Shen SMARCA2 targets UP
MYC targets V1
Wong embryonic stem cell core
Gary CD5 targets DN
Hsiao housekeeping genes
Grade colon and rectal cancer UP
Amino sugar and nucleotide sugar metabolism
Proteasome
Setlur prostate cancer TMPRSS2−ERG fusion UP
Fold Enrichment
Cluster
Up in White
Up in Black
Gene ratio (%)
Pathway Source
Hallmark
KEGG
MSigDB C2
B Key enriched pathways of differentially abundant proteins, age & PSA adjusted (p < 0.05)
Self-identification
GG
pT Stage
DxAGE
DxPSA
Oxidative phosphorylation
Androgen response
Amino acid metabolism
EGF EGFR signaling
Fatty acid metabolism
Xenobiotic metabolism
Glycolysis gluconeogenesis
MTORC1 signaling
DNA repair pathways full network
Metabolic reprogramming in colon cancer
MYC targets V1
Spliceosome
Setlur prostate cancer TMPRSS2−ERG fusion UP
Protein Abundance
Type
Elevated in White
Elevated in Black
Pathway source
Hallmark
KEGG
MSigDB C2
WikiPathways
C Differentially expressed AR targets (adj. p < 0.05)
Self−identification
GG
pT Stage
DxAGE
DxPSA
KHDRBS3
F5
SOX9
SLC26A2
BPHL
RAP1GAP
TSC22D1
FRK
NDRG1
LDHA
ITPR1
PHLDB2
LGALS8
SORL1
PICALM
SEPTIN11
ACO1
SMARCA4
RAB7A
PFKL
PSMD8
KIF5B
PACS1
TKT
TLE1
NAT1
MAOA
INPP4B
ALDH4A1
ALDH1A3
NOSTRIN
ACAD8
DBI
C1orf21
CYB5A
CAMKK2
CRAT
TMEM87B
UBE2J1
TMED5
ABCC4
GREB1
ADPGK
NANS
PCYT2
RPS6KA3
ERBB2
PURA
GFPT1
KRT8
ENDOD1
PC
REEP5
PRDX4
AKAP1
FGD4
BCAP29
KRT19
GSTM3
FDFT1
CLGN
SLC45A3
ADAMTS1
RAB27A
GNMT
Protein Abundance
White
Black
40−49
50−59
60−69
70−79
< 4
4−10
10−20
> 20
D AR regulated cancer driver genes
Relative Protein Abundance
CAMKK2
ERBB2
INPP4B
ALDH1A3
SOX9
LDHA
NDRG1
SMARCA4
Protein
Self-identification White Black
E ERG responsive proteome panel
STXBP1
LRP1
NDRG1
ECE1
ERG
AMPD3
SORL1
AZGP1
SLC4A4
ALOX15B
ANPEP
MAP2
SPON2
ABAT
ALDH6A1
INPP4B
DCXR
ALDH2
FECH
MIPEP
TRPM8
NAT1
RAB27A
EHHADH
F Spearman correlation of training and test datasets
CPDR−APOLLO
Corr: 0.863***
Corr: 0.816***
Sinha
Corr: 0.872***
Setlur
G AUC plots of training and test datasets
Sensitivity (%)
100 − Specificity (%)
CPDR−APOLLO AUC: 0.86
Sinha AUC: 0.92
Setlur AUC: 0.83

**Figure S6. Ancestry-associated differences in molecular pathways and protein signatures (A) Differential pathway enrichment based on race-associated proteome abundance.** Differential pathway enrichment by race-associated age and PSA adjusted proteome abundance**.** Bar plot showing the top significantly enriched pathways (adjusted $p < 0.05$) associated with genes more frequently altered in tumors from Black (blue outline) or White (red outline) patients. Bars show fold enrichment (x-axis), fill color intensity corresponds to the percentage of genes or protein in input set relative to the pathway gene set (Gene ratio %). Asterisks indicate adjusted p-value significance levels. Pathway labels are color coded by database source: ***($p < 0.001$ "), **($p < 0.05$), *($p < 0.1$), $^{ns}$($p > 0.1$). **(B) Heatmap of top enriched pathways from differentially expressed proteins**. Top enrich pathways of age and PSA adjusted differentially expressed proteins ($p < 0.05$). (**C**) **Global proteome of AR-regulated genes.** Heatmap of significantly differentially abundant (Limma adjusted $p < 0.05$) global proteome of AR regulated genes without covariate adjustment. (**D**) **Relative protein abundance of key AR-regulated proteins in Black versus White patients.** Asterisks denote significance of adjusted Wilcoxon rank-sum test (see legend below). **(E) Heatmap of ERG-responsive proteins that discriminate fusion *TMPRSS2-ERG* fusion and ERG protein expression status**. ERG is included as a reference. (**F**) **Correlation of ERG-responsive protein expression between training and validation datasets**. Spearman's rank correlation of the log2FC ERG positive versus ERG negative significant features between the CPDR-APOLLO training dataset with two validation datasets:  proteomics (Sinha et al., 2019) and microarray (Setlur et al., 2008) derived datasets. (**G**) **Discriminatory power of the ERG-responsive protein panel.** Receiver operating characteristic (ROC) curves for sensitivity and specificity of ERG responsive proteins panel in discriminating *TMPRSS2-ERG* fusion and ERG status in CPDR-APOLLO training and the Sinha and Setlur validation datasets by sparse partial least squares discriminant analysis (sPLS-DA).

#### Figure S7

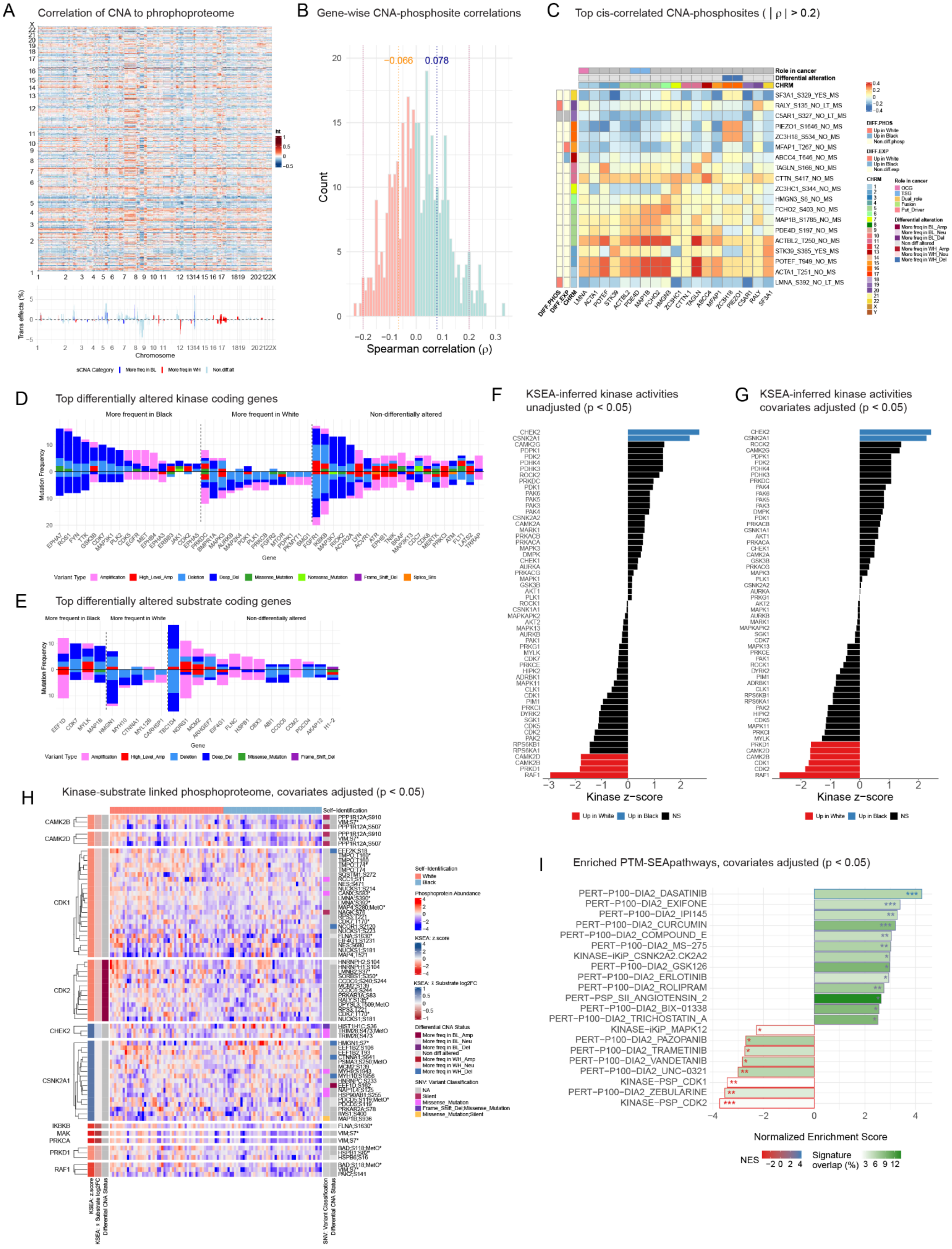

A Correlation of CNA to phrophoproteome
Chromosome
Trans effects (%)
sCNA Category More freq in BL More freq in WH Non.diff.alt
B Gene-wise CNA-phosphosite correlations
−0.066
0.078
Count
Spearman correlation (ρ)
C Top cis-correlated CNA-phosphosites ( | ρ | > 0.2)
Role in cancer
Differential alteration
CHRM
SF3A1_S329_YES_MS
RALY_S135_NO_LT_MS
C5AR1_S327_NO_LT_MS
PIEZO1_S1646_NO_MS
ZC3H18_S534_NO_MS
MFAP1_T267_NO_MS
ABCC4_T646_NO_MS
TAGLN_S166_NO_MS
CTTN_S417_NO_MS
ZC3HC1_S344_NO_MS
HMGN3_S6_NO_MS
FCHO2_S403_NO_MS
MAP1B_S1785_NO_MS
PDE4D_S197_NO_MS
ACTBL2_T250_NO_MS
STK39_S385_YES_MS
POTEF_T949_NO_MS
ACTA1_T251_NO_MS
LMNA_S392_NO_LT_MS
DIFF.PHOS DIFF.EXP CHRM
D Top differentially altered kinase coding genes
More frequent in Black
More frequent in White
Non-differentially altered
Mutation Frequency
Gene
Variant Type Amplification High_Level_Amp Deletion Deep_Del Missense_Mutation Nonsense_Mutation Frame_Shift_Del Splice_Site
E Top differentially altered substrate coding genes
More frequent in Black
More frequent in White
Non-differentially altered
Mutation Frequency
Gene
Variant Type Amplification High_Level_Amp Deletion Deep_Del Missense_Mutation Frame_Shift_Del
F KSEA-inferred kinase activities unadjusted (p < 0.05)
Kinase z-score
Up in White Up in Black NS
G KSEA-inferred kinase activities covariates adjusted (p < 0.05)
Kinase z-score
Up in White Up in Black NS
H Kinase-substrate linked phosphoproteome, covariates adjusted (p < 0.05)
CAMK2B
CAMK2D
CDK1
CDK2
CHEK2
CSNK2A1
IKBKB
MAK
PRKCA
PRKD1
RAF1
KSEA: z.score
KSEA: ε Substrate log2FC
Differential CNA Status
SNV: Variant Classification
Self-Identification
I Enriched PTM-SEApathways, covariates adjusted (p < 0.05)
PERT−P100−DIA2_DASATINIB
PERT−P100−DIA2_EXIFONE
PERT−P100−DIA2_IPI145
PERT−P100−DIA2_CURCUMIN
PERT−P100−DIA2_COMPOUND_E
PERT−P100−DIA2_MS−275
KINASE−iKiP_CSNK2A2.CK2A2
PERT−P100−DIA2_GSK126
PERT−P100−DIA2_ERLOTINIB
PERT−P100−DIA2_ROLIPRAM
PERT−PSP_SII_ANGIOTENSIN_2
PERT−P100−DIA2_BIX−01338
PERT−P100−DIA2_TRICHOSTATIN_A
KINASE−iKiP_MAPK12
PERT−P100−DIA2_PAZOPANIB
PERT−P100−DIA2_TRAMETINIB
PERT−P100−DIA2_VANDETANIB
PERT−P100−DIA2_UNC−0321
KINASE−PSP_CDK1
PERT−P100−DIA2_ZEBULARINE
KINASE−PSP_CDK2
Normalized Enrichment Score
NES
Signature overlap (%)

**Figure S7. Associations of CNA with phosphopeptide abundance, inferred kinase activity, and pathway enrichment. Correlations between somatic copy number alteration (CNA) and phosphopeptide abundance.** (**A**) Heatmap showing *cis*- and *trans*-correlations between somatic CNA alteration (x-axis) to phospho-peptide abundance (y-axis). Of the 359 CNA-phosphoprotein pairs, 184 (51.3%) were positively correlated. Positive correlations are red and negative are blue ($p < 0.05$). *Cis*-effects appear as a red diagonal and *trans*-effects as vertical stripes. The histogram shows the percentage of significant *trans*-effects for each CNA gene. (**B**) Gene-wise correlations between CNA and phosphosite abundance. The blue and orange dashed lines indicate the median of positive (0.078) and negative (-0.066) correlations, respectively. (**C**) Top *cis*-correlated CNA-phosphoproteomic features. Annotations describe differential alteration status and functional roles for both genes and phosphopeptides (**D-E**): **Differential altered kinase and phosphoproteome alterations by race.** Bar charts showing the top differentially altered kinases **(D)** and phosphoproteome substrates **(E)**, ordered by frequency. Counts above the midline represent a higher frequency in Black patients; those below the midline, a higher frequency in White patients. (**F-I**) **Kinase activity and PTM-SEA pathway enrichment.** Bar plots show Kinase-Substrate Enrichment Analysis (KSEA) z-scores for kinases without (**F**) or with covariate adjustment (**G**). Each bar represents an enrichment z-score for kinases with at least two substrates significantly upregulated in Black patients (blue) or White patients (red) ($p <0.05$). Non-significant scores are in black. Analysis was performed with PhosphoSitePlus and NetworKIN datasets, with a NetworKIN score cutoff of 5. **(H)** Heatmap showing imputed, global-normalized phosphoproteome abundance (age and PSA adjusted), clustered by race, for significant kinase-substrate links. Row annotations provide details on KSEA enrichment, CNA, and SNV mutation status for the kinase and substrate genes.
(**I**) Bar plot illustrates the top significantly enriched pathways ($p < 0.05$) from PTM-SEA analysis of phosphosites.
The bars represent fold enrichment, and fill color intensity indicates the proportion of input phosphosites in the pathway's signature set. Asterisks denote statistical significance: *** ($p < 0.001$) , ** ($p < 0.05$), * ($p < 0.1$), and ns ($p > 0.1$).

#### Figure S8

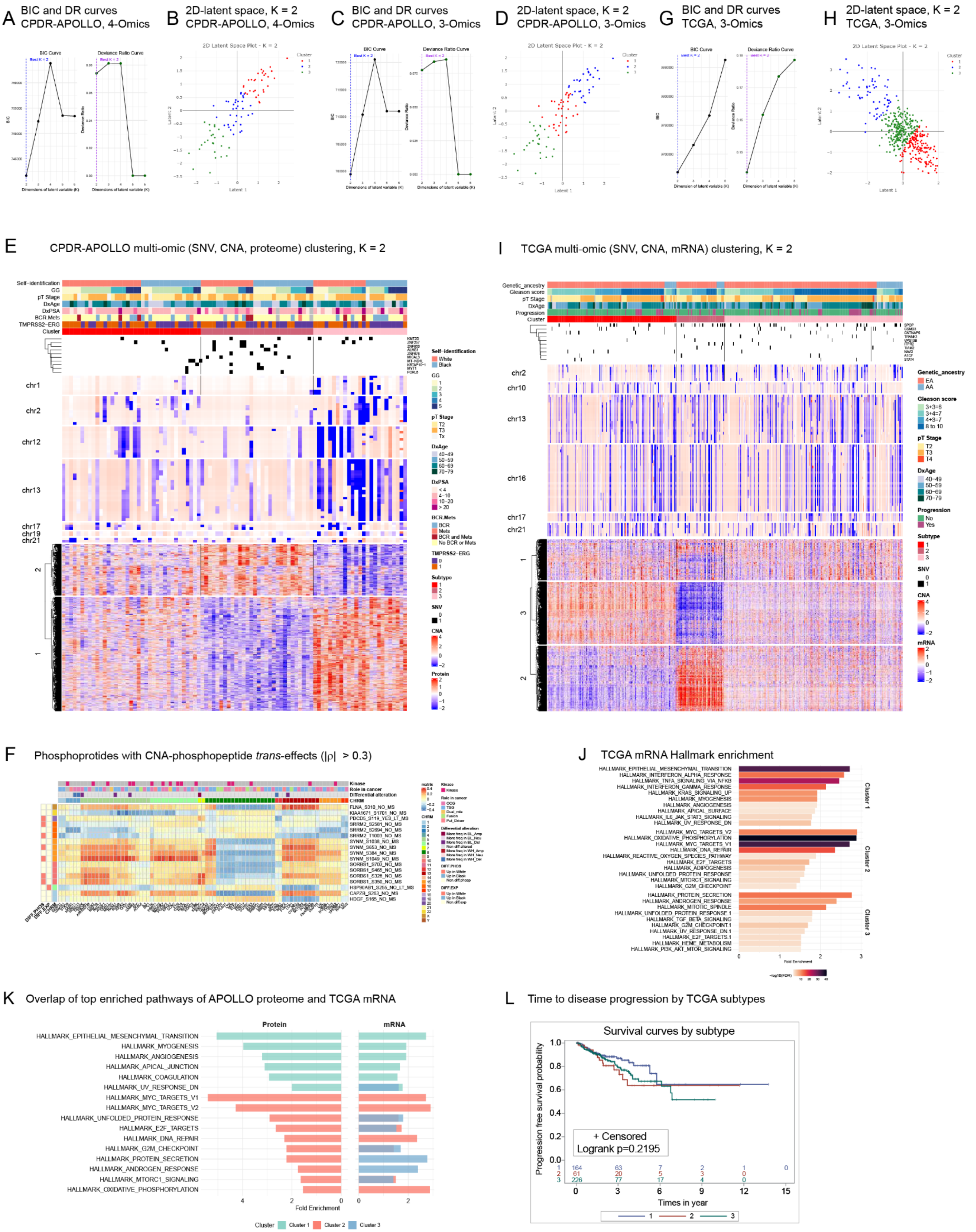

A BIC and DR curves CPDR-APOLLO, 4-Omics
B 2D-latent space, K = 2 CPDR-APOLLO, 4-Omics
C BIC and DR curves CPDR-APOLLO, 3-Omics
D 2D-latent space, K = 2 CPDR-APOLLO, 3-Omics
G BIC and DR curves TCGA, 3-Omics
H 2D-latent space, K = 2 TCGA, 3-Omics
E CPDR-APOLLO multi-omic (SNV, CNA, proteome) clustering, K = 2
I TCGA multi-omic (SNV, CNA, mRNA) clustering, K = 2
F Phosphoprotides with CNA-phosphopepetide trans-effects (|ρ| > 0.3)
J TCGA mRNA Hallmark enrichment
K Overlap of top enriched pathways of APOLLO proteome and TCGA mRNA
Protein
mRNA
HALLMARK_EPITHELIAL_MESENCHYMAL_TRANSITION
HALLMARK_MYOGENESIS
HALLMARK_ANGIOGENESIS
HALLMARK_APICAL_JUNCTION
HALLMARK_COAGULATION
HALLMARK_UV_RESPONSE_DN
HALLMARK_MYC_TARGETS_V1
HALLMARK_MYC_TARGETS_V2
HALLMARK_UNFOLDED_PROTEIN_RESPONSE
HALLMARK_E2F_TARGETS
HALLMARK_DNA_REPAIR
HALLMARK_G2M_CHECKPOINT
HALLMARK_PROTEIN_SECRETION
HALLMARK_ANDROGEN_RESPONSE
HALLMARK_MTORC1_SIGNALING
HALLMARK_OXIDATIVE_PHOSPHORYLATION
Fold Enrichment
Cluster
Cluster 1
Cluster 2
Cluster 3
L Time to disease progression by TCGA subtypes
Survival curves by subtype
Progression free survival probability
+ Censored
Logrank p=0.2195
Times in year

**Figure S8. Integrated molecular and pathway-based subtyping.** (**A-F**): **Molecular subtyping of discovery cohort (CPDR-APOLLO)**. We performed integrated clustering using 4-omics (SNV, CNA, proteomics, and phosphoproteomics), and 3-omics (SNV, CNA, and proteomics) data to identify molecular subtypes. (**A**, **C**) Model fit evaluation for the 4-omics and 3-omics data, respectively, showing that a latent dimension of K = 2 is optimal based on Bayesian Information Criterion (BIC) and deviance ratio plots. (**B, D**) Latent space visualization at K=2, with unsupervised clustering revealing three distinct clusters. (**E**) Heatmap of the 3-omics clustering, displaying patient clusters (columns), and molecular features (rows), with annotations for race, clinical-pathological features, and *TMPRSS2-ERG* status. (**F**) *Trans*-effects heatmap showing correlation between CNAs of driver genes/kinases and phosphoproteomic features. (**G-L**): **Integrated Analysis of validation cohort (TCGA)**. We applied the same *iClusterBayes* analysis to TCGA's 3-omics data (SNV, CNA, and mRNA expression) to validate our findings. (**G**) **Model selection metrics** confirm K=2 as the optimal latent dimension. (**H**) **Latent space visualization** showing three distinct clusters. (**I**) Heatmap of the TCGA clustering, displaying patient clusters and molecular features, annotated by race and clinical-pathological features. (**J**) **mRNA pathway enrichment** showing the top 10 significantly enriched Hallmark pathways for each cluster. (**K**) **Shared pathways enrichment** across protein (CPDR-APOLLO) and mRNA (TCGA) clusters, highlighting significantly enriched pathways in both cohorts. (**L**) Univariate Kaplan-Meier curves comparing the progression-free interval (PFI) among the three assigned patient clusters.

### Figure S9

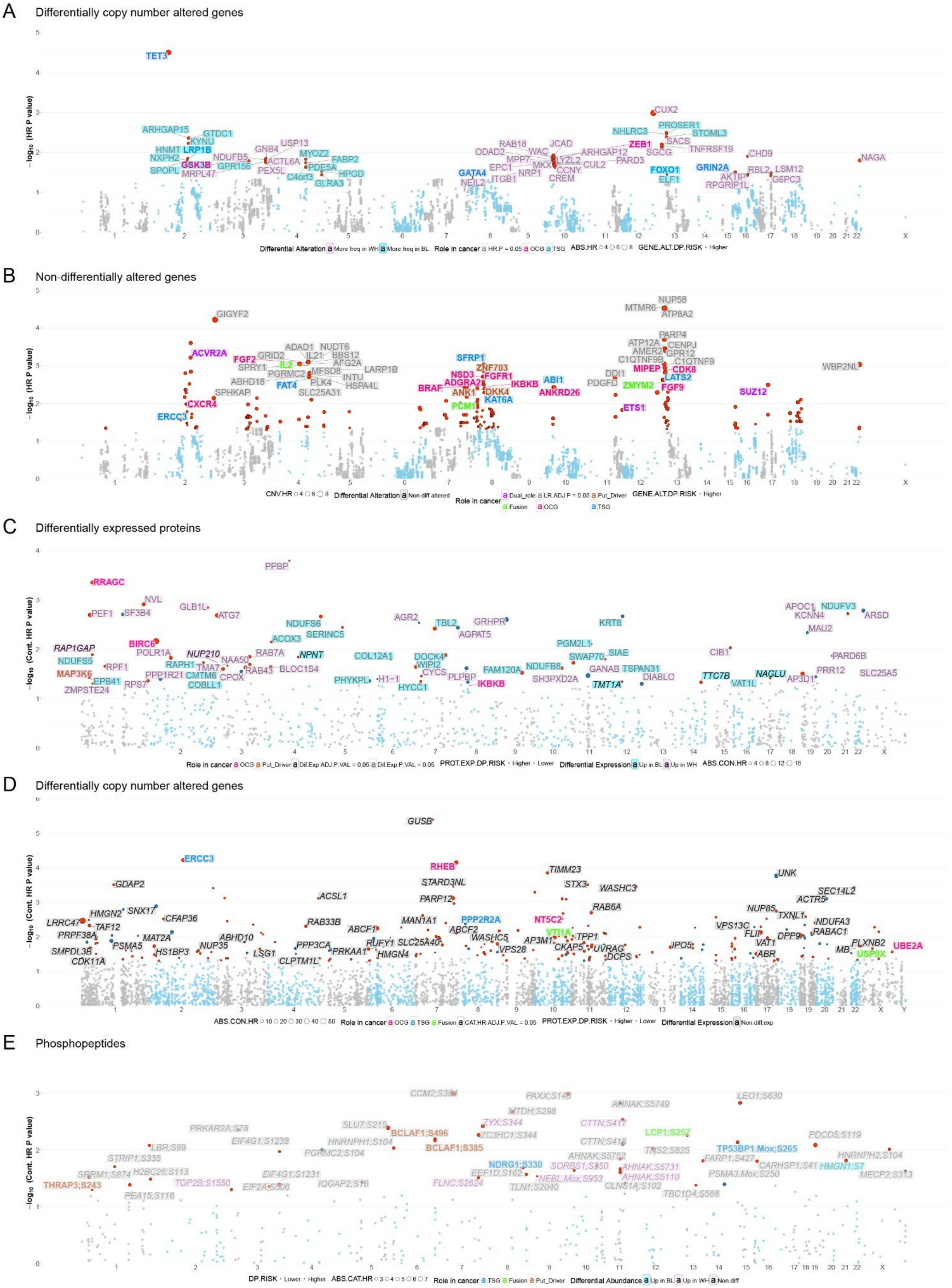
A
Differentially copy number altered genes
-log10 (HR P value)
TET3
CUX2
ARHGAP15
GTDC1
KYNU
HNMT
LRP1B
NXPH2
GSK3B
SPOPL
MRPL47
NDUFB5
GPR156
PEX5L
GNB4
USP13
ACTL6A
MYOZ2
FABP2
PDE5A
C4orf3
HPGD
GLRA3
GATA4
NEIL2
RAB18
ODAD2
MPP7
EPC1
ITGB1
WAC
MKX
NRP1
JCAD
LYZL2
CCNY
CREM
ARHGAP12
CUL2
PARD3
NHLRC3
ZEB1
FOXO1
ELF1
PROSER1
SACS
SGCG
STOML3
TNFRSF19
GRIN2A
AKTIP
RPGRIP1L
CHD9
RBL2
LSM12
G6PC3
NAGA
Differential Alteration
More freq in WH
More freq in BL
Role in cancer
HR.P = 0.05
OCG
TSG
ABS.HR
GENE.ALT.DP.RISK
Higher
B
Non-differentially altered genes
-log10 (HR P value)
GIGYF2
MTMR6
NUP58
ATP8A2
ACVR2A
FGF2
GRID2
ADAD1
IL21
NUDT6
BBS12
AFG2A
SPRY1
IL2
PGRMC2
MFSD8
LARP1B
INTU
ABHD18
FAT4
PLK4
HSPA4L
SPHKAP
SLC25A31
CXCR4
ERCC3
SFRP1
ZNF703
NSD3
FGFR1
ADGRA2
BRAF
ANK1
DKK4
IKBKB
ABI1
ANKRD26
KAT6A
PCM1
PARP4
ATP12A
CENPJ
AMER2
GPR12
C1QTNF9B
C1QTNF9
DDI1
PDGFD
MIPEP
CDK8
LATS2
ZMYM2
FGF9
ETS1
SUZ12
WBP2NL
CNV.HR
Differential Alteration
Non diff altered
Role in cancer
Dual_role
LR.ADJ.P = 0.05
Put_Driver
Fusion
OCG
TSG
GENE.ALT.DP.RISK
Higher
C
Differentially expressed proteins
-log10 (Cont. HR P value)
PPBP
RRAGC
NVL
PEF1
SF3B4
GLB1L
ATG7
APOC1
NDUFV3
KCNN4
ARSD
MAU2
AGR2
TBL2
GRHPR
AGPAT5
NDUFS6
SERINC5
KRT8
RAP1GAP
BIRC6
NDUFS5
POLR1A
NUP210
ACOX3
RAB7A
NPNT
COL12A1
DOCK4
WIPI2
CYCS
PGM2L1
SIAE
CIB1
PARD6B
MAP3K6
RPF1
RAPH1
TMA7
NAA50
RAB43
BLOC1S4
FAM120A
NDUFB8
SWAP70
GANAB
TSPAN31
TTC7B
NAGLU
PRR12
SLC25A5
EPB41
PPP1R21
CMTM6
CPOX
PHYKPL
H1-1
PLPBP
IKBKB
SH3PXD2A
DIABLO
VAT1L
AP3D1
ZMPSTE24
RPS7
COBLL1
HYCC1
TMT1A
Role in cancer
OCG
Put_Driver
Dif.Exp ADJ.P.VAL = 0.05
Dif.Exp P.VAL = 0.05
PROT.EXP.DP.RISK
Higher
Lower
Differential Expression
Up in BL
Up in WH
ABS.CON.HR
D
Differentially copy number altered genes
-log10 (Cont. HR P value)
GUSB
ERCC3
RHEB
STARD3NL
PARP12
TIMM23
STX3
WASHC3
RAB6A
UNK
SEC14L2
ACTR5
NUP85
TXNL1
NDUFA3
RABAC1
GDAP2
ACSL1
HMGN2
SNX17
CFAP36
LRRC47
TAF12
PRPF38A
MAT2A
SMPDL3B
PSMA5
HS1BP3
CDK11A
NUP35
ABHD10
LSG1
RAB33B
ABCF1
PPP3CA
PRKAA1
RUFY1
SLC25A40
HMGN4
CLPTM1L
MAN1A1
ABCF2
PPP2R2A
WASHC5
VPS28
NT5C2
VTI1A
AP3M1
CKAP5
TPP1
UVRAG
DCPS
IPO5
VPS13C
FLII
VAT1
DPP9
ABR
MB
PLXNB2
USP9X
UBE2A
ABS.CON.HR
Role in cancer
OCG
TSG
Fusion
CAT.HR.ADJ.P.VAL = 0.05
PROT.EXP.DP.RISK
Higher
Lower
Differential Expression
Non.diff.exp
E
Phosphopeptides
-log10 (Cont. HR P value)
CCM2;S384
PAXX;S148
AHNAK;S5749
LEO1;S630
MTDH;S298
ZYX;S344
ZC3HC1;S344
CTTN;S417
SLU7;S215
BCLAF1;S496
BCLAF1;S385
PRKAR2A;S78
EIF4G1;S1238
HNRNPH1;S104
PGRMC2;S104
LBR;S99
STRIP1;S335
H2BC26;S113
SRRM1;S874
THRAP3;S243
PEA15;S116
TOP2B;S1550
EIF4G1;S1231
EIF2A;S506
IQGAP2;S16
FLNC;S2624
NDRG1;S330
EEF1D;S162
TLN1;S2040
SORBS1;S350
NEBL.Mox;S953
CTTN;S418
AHNAK;S5752
LCP1;S257
TNS2;S825
AHNAK;S5731
AHNAK;S5110
CLNS1A;S102
TBC1D4;S588
FARP1;S427
TP53BP1.Mox;S265
PSMA3.Mox;S250
PDCD5;S119
CARHSP1;S41
HMGN1;S7
HNRNPH2;S104
MECP2;S313
DP.RISK
Lower
Higher
ABS.CAT.HR
Role in cancer
TSG
Fusion
Put_Driver
Differential Abundance
Up in BL
Up in WH
Non.diff

**Figure S9. Manhattan plots showing the association molecular features with progression to biochemical recurrence (BCR) or metastasis in the CPDR -APOLLO cohort.** The x-axis represents chromosomal location, and the y-axis shows -log10 transformed hazard ratio (HR) p-values, reflecting the strength of association with disease progression. Chromosomes are alternately colored gray and sky blue. Point size scales with the absolute HR magnitude. Statistical analyses were adjusted for age and PSA levels at diagnosis. (**A-B**) **Association of CNA events with disease progression:** (**A**) shows differentially altered genes, (**B**) non-differentially altered genes. Genes with significant associations (HR $p < 0.05$, absolute HR > 1.3) are indicated by larger red points. Labels highlight notable cancer-relevant genes: the text color indicates the gene's role in cancer, and the background color represents its differential alteration category. Genes without an established cancer role but with a significant association are shown in plain text. (**C, D, and E**): **Proteins and Phosphopeptides .** The plots illustrate changes in protein **(C, D)** and phosphopeptide **(E)** abundance. Highlighted points represent features with significant associations based on absolute HR > 1.5, continuous HR $p < 0.05$, categorical $p < 0.05$, or adjusted categorical $p < 0.05$ and differential protein or phosphopeptide abundance (age- and PSA-adjusted Limma $p < 0.05$). The point color denotes the direction of risk: **red** indicates higher risk with elevated abundance or phosphorylation, and **blue** indicates higher risk with decreased abundance or phosphorylation. Labels highlight notable cancer-relevant molecular features. The label's **text color** indicates the molecular feature's established cancer role, while its background color denotes the differential abundance or phosphorylation category. Features lacking an established cancer role are shown in italic or plain text based on their adjusted p-value thresholds

**Figure S10**

A CNAs prevalent in Black; increased risk

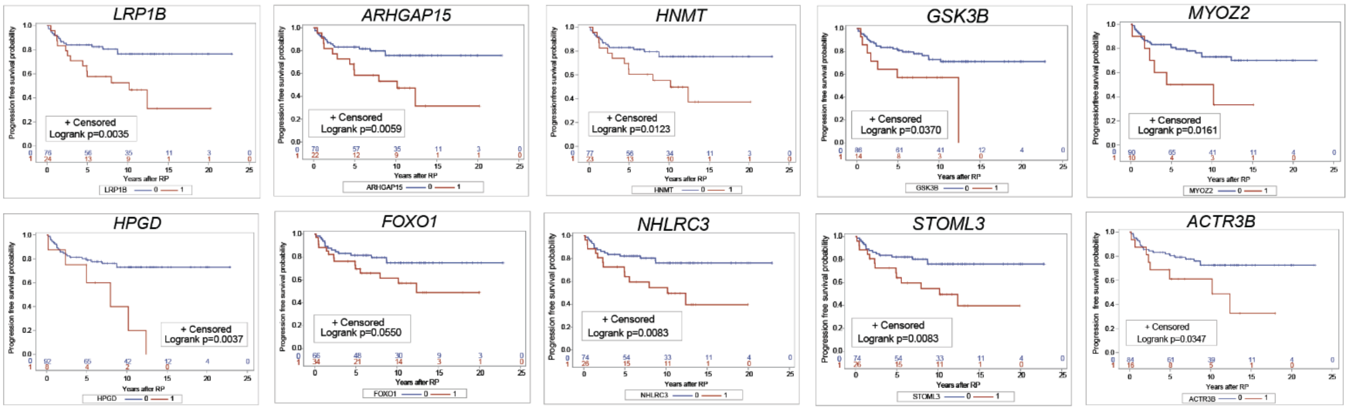


B CNAs prevalent in White; increased risk

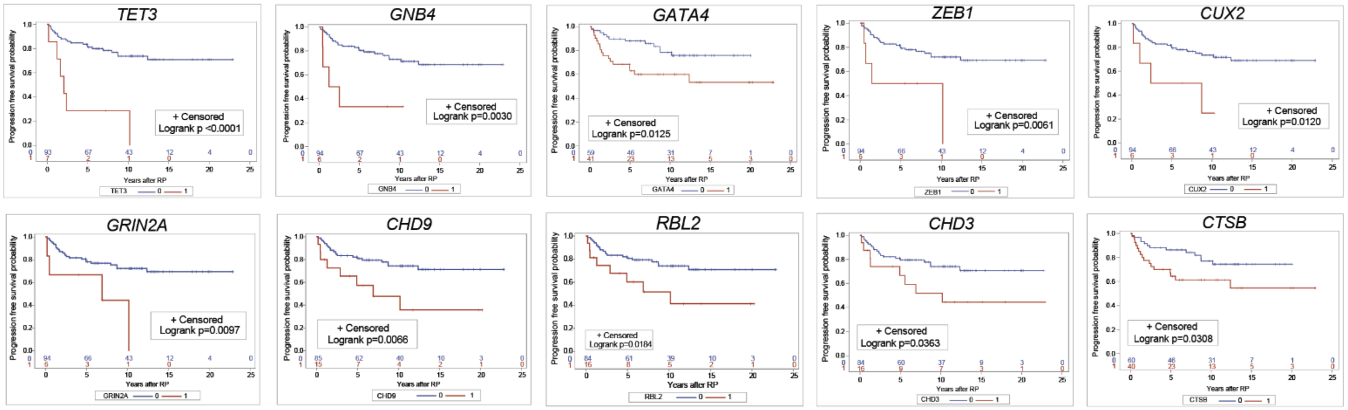


C Non-differentially altered CNAs; increased risk

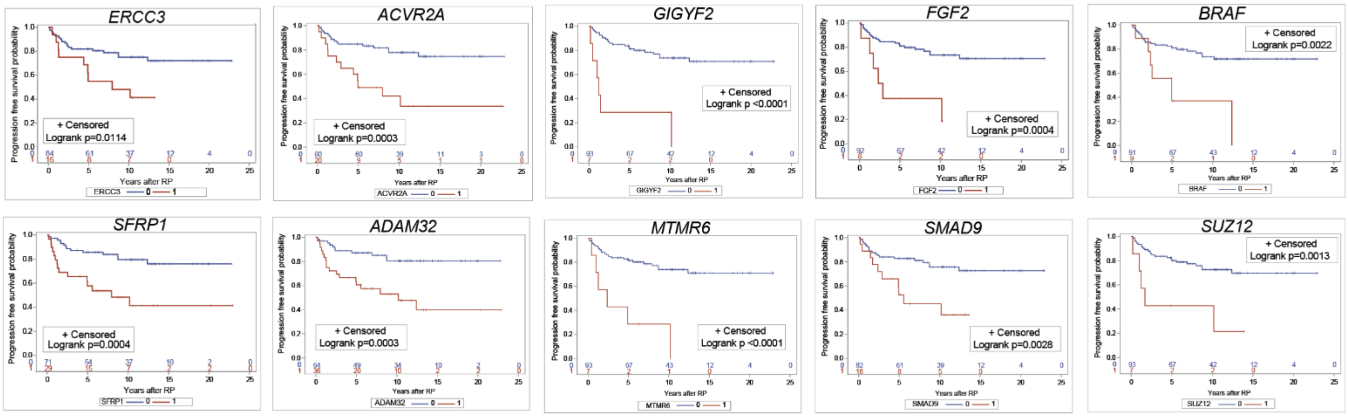


D TCGA validation, prevalent in White (Concordant)

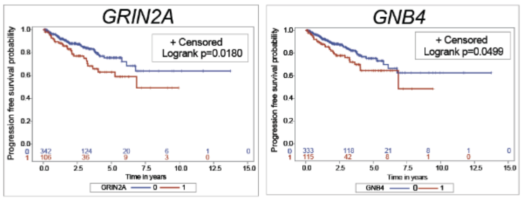


E TCGA validation, Non-differentially altered (Concordant)

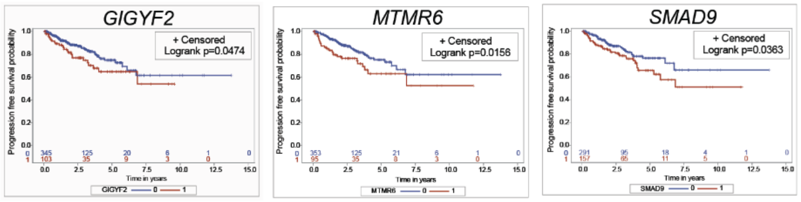

**Figure S10. Progression-free survival by copy number alteration (CNA) status.** Unadjusted univariate Kaplan-Meier curves show the time from surgery to BCR or metastasis, stratified by gene alteration status. The top panels (**A-C**)show results from the primary cohort, while the bottom panels (**D-E**) show external validation using the TCGA cohort. (**A**) CNAs frequently altered in Black patients and associated with increased risk. **(B)** CNAs frequently altered in White patients and associated with increased risk. **(C)** CNAs with similar frequency across ancestries and associated with increased risk. External validation curves for prognostic CNAs more prevalent in White patients (**D**), and for non-differentially altered CNAs (**E**).

**Figure S11**

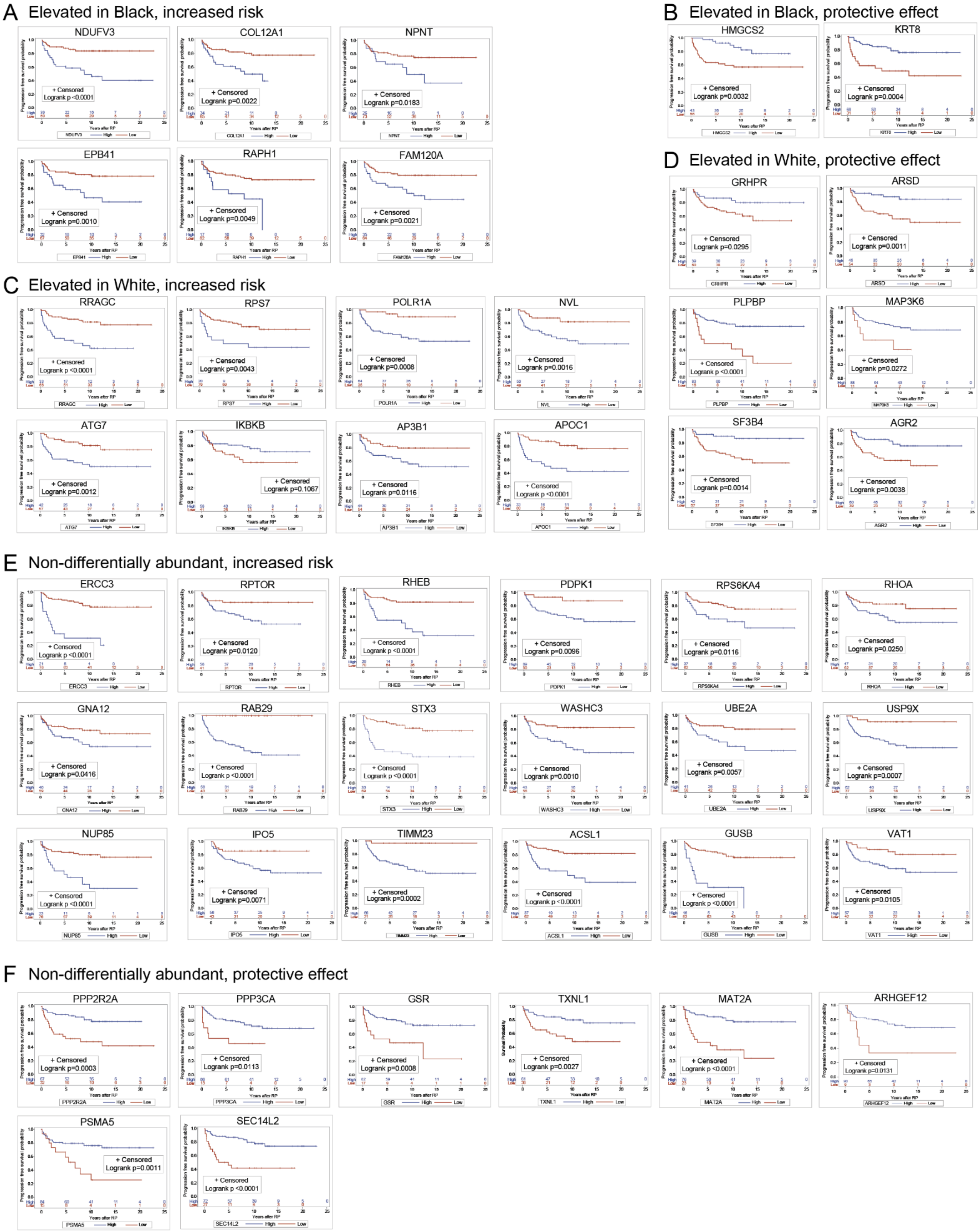

A Elevated in Black, increased risk
NDUFV3
Logrank p<0.0001
COL12A1
Logrank p=0.0022
NPNT
Logrank p=0.0183
EPB41
Logrank p=0.0010
RAPH1
Logrank p=0.0049
FAM120A
Logrank p=0.0021
B Elevated in Black, protective effect
HMGCS2
Logrank p=0.0032
KRT8
Logrank p=0.0004
C Elevated in White, increased risk
RRAGC
Logrank p<0.0001
RPS7
Logrank p=0.0043
POLR1A
Logrank p=0.0008
NVL
Logrank p=0.0016
ATG7
Logrank p=0.0012
IKBKB
Logrank p=0.1067
AP3B1
Logrank p=0.0116
APOC1
Logrank p<0.0001
D Elevated in White, protective effect
GRHPR
Logrank p=0.0295
ARSD
Logrank p=0.0011
PLPBP
Logrank p<0.0001
MAP3K6
Logrank p=0.0272
SF3B4
Logrank p=0.0014
AGR2
Logrank p=0.0038
E Non-differentially abundant, increased risk
ERCC3
Logrank p<0.0001
RPTOR
Logrank p=0.0120
RHEB
Logrank p<0.0001
PDPK1
Logrank p=0.0096
RPS6KA4
Logrank p=0.0116
RHOA
Logrank p=0.0250
GNA12
Logrank p=0.0416
RAB29
Logrank p<0.0001
STX3
Logrank p<0.0001
WASHC3
Logrank p=0.0010
UBE2A
Logrank p=0.0057
USP9X
Logrank p=0.0007
NUP85
Logrank p<0.0001
IPO5
Logrank p=0.0071
TIMM23
Logrank p=0.0002
ACSL1
Logrank p<0.0001
GUSB
Logrank p<0.0001
VAT1
Logrank p=0.0105
F Non-differentially abundant, protective effect
PPP2R2A
Logrank p=0.0003
PPP3CA
Logrank p=0.0113
GSR
Logrank p=0.0008
TXNL1
Logrank p=0.0027
MAT2A
Logrank p<0.0001
ARHGEF12
Logrank p=0.0131
PSMA5
Logrank p=0.0011
SEC14L2
Logrank p<0.0001
+ Censored
Progression free survival probability
Years after RP
High
Low

**Figure S11. Progression-free survival by protein abundance.** Unadjusted univariate Kaplan-Meier curves depict time from surgery to BCR or metastasis, stratified into high and low expression groups based on the Youden index cutoff. (**A-B**) Progression-free survival curves for proteins significantly elevated in Black patients, indicating association with increased risk **(A)** and those offering a protective effect (**B**). (**C-D**) Progression-free survival for proteins significantly elevated in White patients, with **(C)** showing increased risk, and (**D**) a protective effect. (**E-F**) Progression-free survival for proteins that are equally abundant across ancestries, with (**E**) associated with increased risk and (**F**) demonstrating a protective effect**.**

**Figure S12**

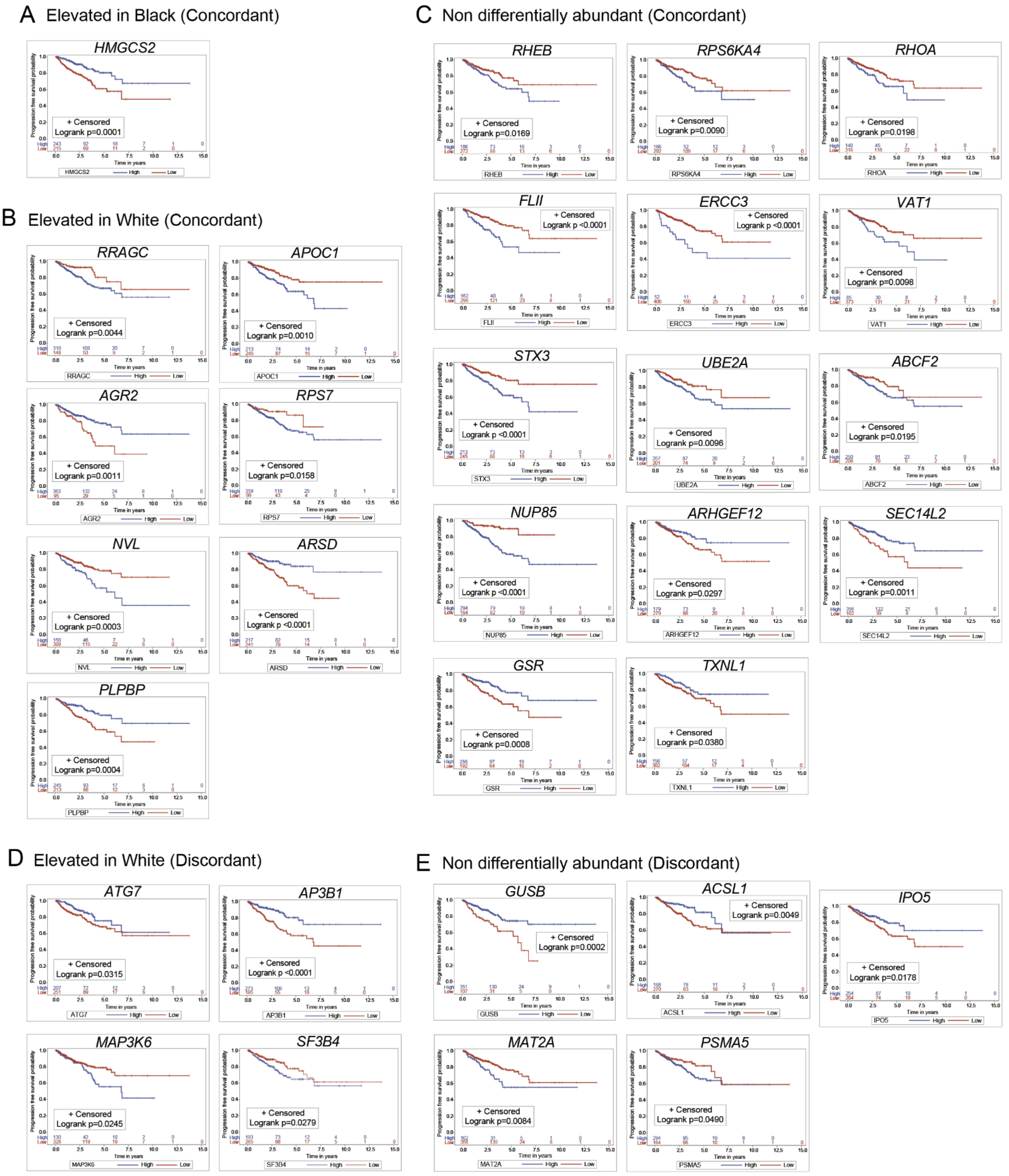
A Elevated in Black (Concordant)
HMGCS2
Logrank p=0.0001
B Elevated in White (Concordant)
RRAGC
Logrank p=0.0044
APOC1
Logrank p=0.0010
AGR2
Logrank p=0.0011
RPS7
Logrank p=0.0158
NVL
Logrank p=0.0003
ARSD
Logrank p <0.0001
PLPBP
Logrank p=0.0004
C Non differentially abundant (Concordant)
RHEB
Logrank p=0.0169
RPS6KA4
Logrank p=0.0090
RHOA
Logrank p=0.0198
FLII
Logrank p <0.0001
ERCC3
Logrank p <0.0001
VAT1
Logrank p=0.0098
STX3
Logrank p <0.0001
UBE2A
Logrank p=0.0096
ABCF2
Logrank p=0.0195
NUP85
Logrank p <0.0001
ARHGEF12
Logrank p=0.0297
SEC14L2
Logrank p=0.0011
GSR
Logrank p=0.0008
TXNL1
Logrank p=0.0380
D Elevated in White (Discordant)
ATG7
Logrank p=0.0315
AP3B1
Logrank p <0.0001
MAP3K6
Logrank p=0.0245
SF3B4
Logrank p=0.0279
E Non differentially abundant (Discordant)
GUSB
Logrank p=0.0002
ACSL1
Logrank p=0.0049
IPO5
Logrank p=0.0178
MAT2A
Logrank p=0.0084
PSMA5
Logrank p=0.0490

**Figure S12. External validation for prognostic proteins using mRNA dataset from the TCGA cohort.** Unadjusted univariate Kaplan-Meier curves depict time from surgery to BCR or metastasis, stratified into high and low expression groups based on the Youden index cutoff. (**A**) Progression-free survival curves for a protein significantly elevated in Black patients, showing concordance in association with a protective effect (HMGCS2). (**B**) Progression-free survival curves for proteins significantly elevated in White patients, showing concordance in association with an increased risk (RRAGC, APOC1, AGR2, RPS7, NVL) or with a protective effect (ARSD, PLPBP). (**C**) Progression-free survival curves for equally abundant proteins, showing concordance in association with an increased risk (RHEB, RPS6KA4, RHOA, FLI1, ERCC3, VAT1, STX3, UBE2A, ABCF2, NUP85) or with a protective effect (ARHGEF12, SEC14L3, GSR, TXNL1). (**D**) Progression-free survival curves for proteins significantly elevated in White patients, showing discordance in association with an increased risk (ATG7, AP3B1) or with a protective effect (MAP3K6, SF3B41). (**E**) Progression-free survival curves for equally abundant proteins, showing discordance in association with an increased risk (GUSB, ASCL1, IPO5) or with a protective effect (MAT2A, PSMA5).

### Figure S13

A Phosphopeptides elevated in Black patients

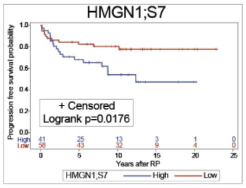


B Phosphopeptides elevated in White patients

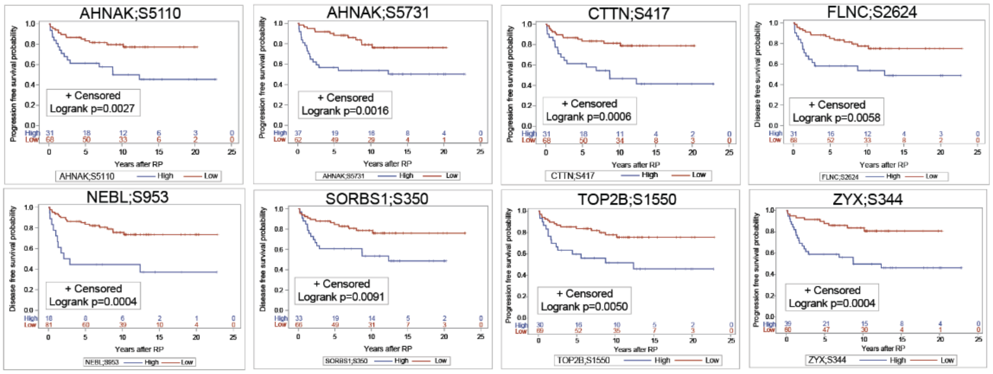


C Non-differentially abundant phosphopeptides

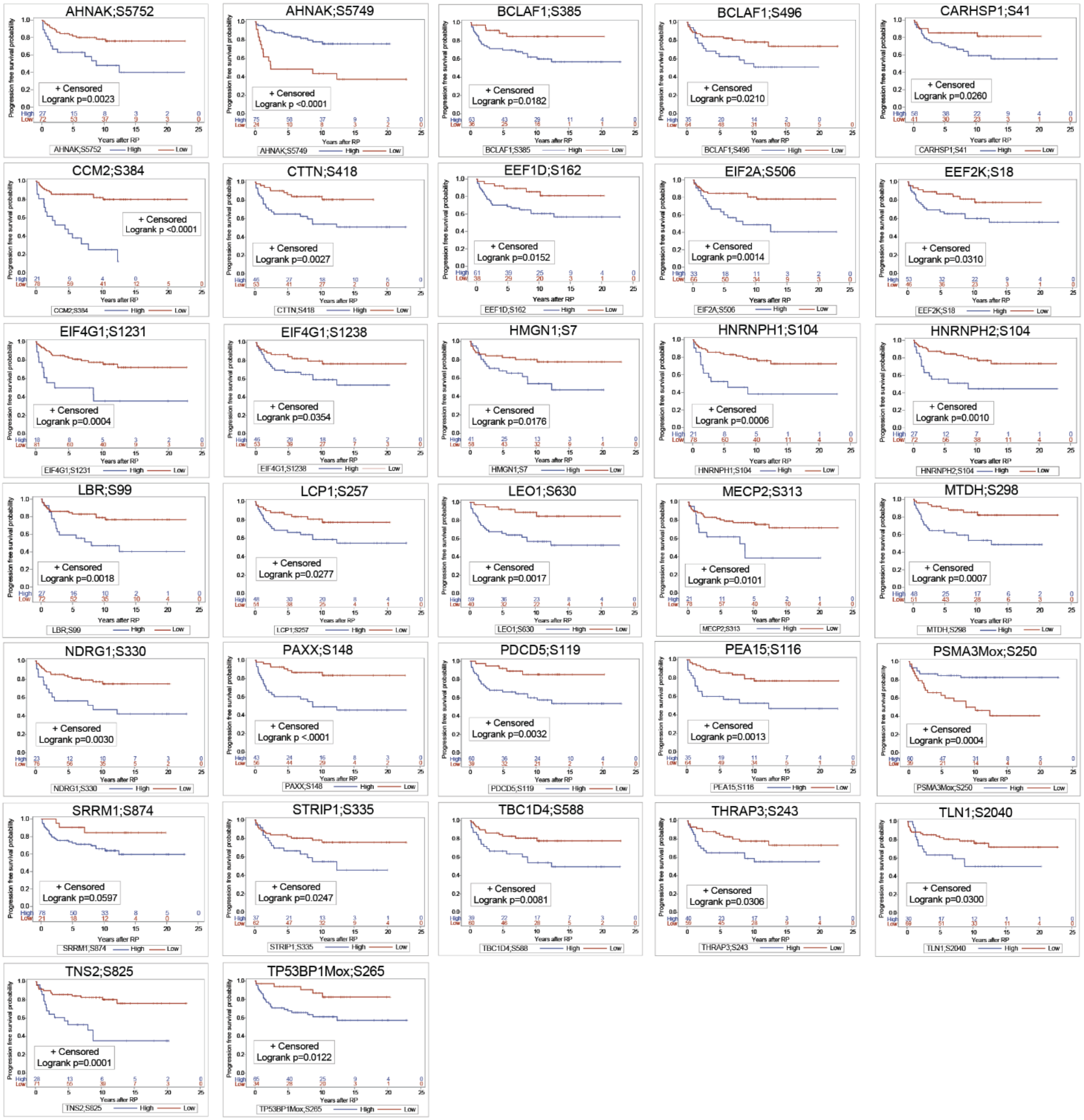

**Figure S13. Progression-free survival by phosphoprotein abundance.** Unadjusted univariate Kaplan-Meier curves illustrate time from surgery to BCR or metastasis, stratified by high and low abundance groups based on the Youden index cutoff. (**A**-**B**) Progression-free survival for phosphopeptides associated with increased risk and significantly elevated in Black and White patients, respectively. (**C**) Progression-free survival for phosphopeptides equally abundant across ancestries (listed alphabetically). With the exception of AHNAK(S5749) and PSMA3(S250), which demonstrate a protective effect, increased abundance of the remaining phosphopeptides in this figure were associated with an increased risk for disease progression.
